# Low-cost scalable discretization, prediction and feature selection for complex systems

**DOI:** 10.1101/720441

**Authors:** S. Gerber, L. Pospisil, M. Navandar, I. Horenko

## Abstract

Finding reliable discrete approximations of complex systems is a key prerequisite when applying many of the most popular modeling tools. Common discretization approaches (for example, the very popular K-means clustering) are crucially limited in terms of quality and cost. We introduce a low-cost improved-quality Scalable Probabilistic Approximation (SPA) algorithm, allowing for simultaneous data-driven optimal discretization, feature selection and prediction. Cross-validated applications of SPA to a range of large realistic data classification and prediction problems reveal drastic cost and performance improvements. For example, SPA allows the unsupervised next-day surface temperature predictions for Europe with the mean crossvalidated one-day prediction error of 0.75°C on a common PC (being around 40% better in terms of errors and five to six orders-of-magnitude cheaper than the next-day surface temperature predictions calculated on supercomputers and provided by the weather services).

**One Sentence Summary:** Introduced computational tool allows obtaining drastic cost and quality gains for a broad range of science applications.

---

Computers are finite discrete machines. Computational treatment and practical simulations of real world systems rely on the approximation of any given system’s state X(t) (where t=1,…,T) in terms of a finite number K of discrete states S={S_1_,…, S_K_}^*1,2*^. Of particular importance are discretization methods that allow the representation of the system’s states X(t) as a vector of K probabilities for the system to be in some particular state S_i_ at the instance t. Components of such a vector - 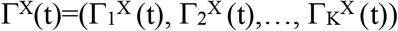 - sum-up to one and are particularly important since they are necessary for Bayesian and Markovian modeling of these systems^*3–5*^.

Bayesian and Markovian models belong to the most popular tools for mathematical modeling and computational data analysis problems in science (with over one million literature references each, according to Google Scholar). They were applied to problems ranging from social and network sciences^6^ to a biomolecular dynamics and drug design^7–9^, fluid mechanics^10^ and climate^11^. These models dwell on the law of the total probability, saying that the exact relation between the given probabilistic representations Γ^Y^(t) and Γ^X^(t) of any two processes Y and X is given as a linear model:

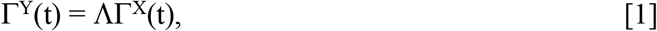

where Λ is a stochastic matrix of conditional probabilities between the discrete states of Y and X. This linear model is exact in a probabilistic sense – meaning that it does not impose a modeling error, even if the underlying dynamics of X and Y is arbitrarily complex and nonlinear. If Λ is known, then [1] provides the best relation between the two information sources Γ^Y^(t) and Γ^X^(t)^2–4^.

A particular – and very important - case of the Bayesian models [1] emerges when choosing Y(t) as X(t+1), where t is a time index. The relation matrix Λ is then a left-stochastic square matrix of transition probabilities between two discrete states, formally known as transfer operator. A Bayesian model [1] in this particular case is called a Markov model^2–4^. Besides of their direct relation to the exact law of total probability, another reason for their popularity - especially in the natural sciences - is the fact that these models automatically satisfy important physical conservation laws. They e.g. exactly conserve probability and herewith lead to stable simulations^2,7,9^. Various efficient computational methods allow to estimate conditional probability matrices Λ for real-world systems^7–15^.

In practice, all these methods require a priori availability of discrete probabilistic representations. Obtaining such representations/approximations Γ^X^(t) by means of common methods from the original system’s states X(t), is subject to serious quality and cost limitations. For example, applicability of grid discretization methods - covering original system’s space with a regular mesh of boxes {S_1_,…, S_K_} is limited in terms of cost - since the required number of boxes K grows exponentially with the dimension n of X(t)^1^.

Because of that, the most common approaches for tackling these kinds of problems are so-called meshless methods. They attempt to find a discretization by means of grouping the states X(t) into K clusters according to some similarity criteria. The computational costs for popular clustering algorithms^16^ as well as for most mixture models^17^ scale linearly with the dimensionality *n* of the problem and the amount of data, *T*. This cheap computation made clustering methods the most popular meshless discretization tools – even despite of the apparent quality limitations they entail. For example, K-means clustering (the most popular clustering method, with over 3 million Google Scholar citations) can only provide probabilistic approximations with binary (zero/one) Γ^X^(t) elements, excluding any other approximations and not guaranteeing optimal approximation quality. Mixture models are subject to similar quality issues when the strong assumptions that they impose (like Gaussianity in Gaussian Mixture Models) are not fulfilled.

Closely related to clustering methods are various approaches for matrix factorization – like the non-negative matrix factorization methods (NMF) that attempt to find an optimal approximation of the given (non-negative) n-times-T data matrix X with a product of the n-times-K matrix S and the K-times-T matrix Γ^X^^18–24^.

In situations where K is smaller than T, such non-negative reduced approximations SΓ^X^ are computed by means of the fixed-point-iterations^19,21^ or by alternating least-squares algorithms and projected gradient methods^*22*^. However, due to the computational cost issues, probabilistic factorizations (i.e., such approximations SΓ^X^ that the columns of Γ are probability distributions) are either excluded explicitly^22^ or they are obtained by means of the spectral decomposition of the data similarity matrices (like the X^T^X matrix in the Euclidean space TxT). Such probabilistic NMF variants like the Left-Stochastic Decomposition (LSD)^24^ – as well as the closely-related spectral decomposition methods^25^ and the robust Perron cluster analysis^8,12^ – are subject to cost limitations.

These cost limitations are induced by the fact that even the most efficient tools for eigenvalue problem computations (where all these methods rely on) scale polynomial with the similarity matrix dimension T. If the similarity matrix does not exhibit any particular structure (i.e., if it is not sparse), the overall numerical cost of the eigenvalue decomposition scales as O(T^3^). For example, considering twice as much data will lead to a two-to three-fold increase of cost.

Similar scalability limitations are also characteristic for the density-based clustering methods (such as the mean shifts^42^, the DBSCAN^43^ and the algorithms based on t-SNE^44^), having an iteration complexity in the orders between O(T*log*(T)) and O(T^2^). Practical applicability of such methods is restricted to relatively-small systems - or relies on the ad hoc data reduction steps (i.e., T cannot routinely exceed 10,000 or 20,000 when working on commodity hardware, see for example the green surface in the Fig. 1a)^9,44^.

**Fig. 1.**
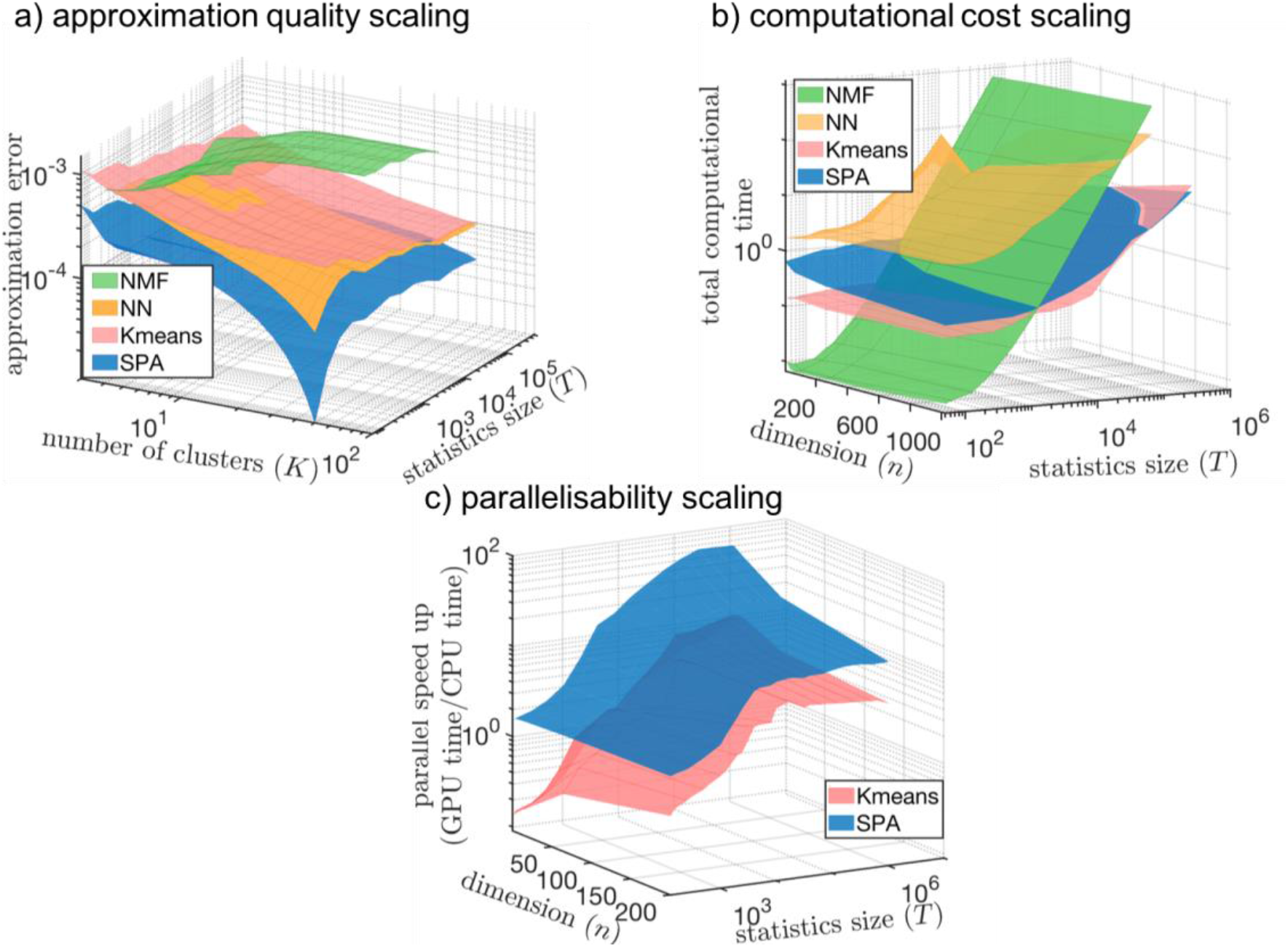
Comparing computational cost (a), discretization quality (b) and parallelizability (c) for SPA (blue surfaces) and for common discretization methods: K-means clustering^16,17^ (red), Nonnegative Matrix Factorisation^19–24^ (in its probabilistic variant called Left-Stochastic Decomposition^24^ (LSD), green surfaces) and the Self-Organising Maps^33^ (SOM, a special form of unsupervised neuronal networks used for discretization, orange surfaces). For every combination of data dimension *n* and the data statistics length *T*, methods are applied to 25 same randomly-generated data sets and the results in each of the curves represent averages over these 25 problems. Parallel speed-up in (c) is measured as the ratio of the average times *time(GPU)*/*time(CPU)* needed to reach the same relative tolerance threshold of 10^−5^ on a single Graphics Processing Unit (GPU, ASUS TURBO-GTX1080TI-11G, with 3584 CUDA cores) for *time (GPU)* versus a single CPU core (Intel Core i9-7900X CPU) for *time(CPU)*. Further comparisons can be found in the Fig. S2 from the Supplement. MATLAB script *Fig1_reproduce.m* reproducing these results is available in the repository *SPA* at *https://github.com/SusanneGerber*

Cost and quality comparison for the probabilistic approximation methods is shown in the Fig. 1. Cost factor becomes decisive when discretizing very large systems, for example in biology and geosciences, leading to the necessity of some ad hoc data pre-processing, by means of computationally-cheap methods like K-means, Principal Component Analysis (PCA) and other pre-reduction steps^26,27^.

In the following, we present a method not requiring such ad hoc reductional data pre-processing, having the same leading order computational iteration complexity O(nKT) as the cheap K-means algorithm, and allowing simultaneously finding discretizations that are optimal for models [1].

## Cost, quality and parallelizability in Scalable Probabilistic Approximation (SPA)

Construction and derivation of many computational methods can be frequently approached by casting the problem into the optimization framework. For example, an approximation quality of a discretization can be expressed as a sum of all distances distS(X(t), Γ^X^(t)) between the original state X(t) and its probabilistic discrete representation Γ^X^(t) that is obtained for S={S_1_,…, S_K_}. For example, minimizing the sum of the squared Euclidean distances 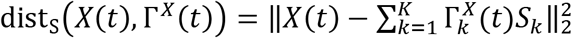 with respect to Γ and *S* for a fixed given X would allow to find the optimal probabilistic approximations 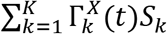 of the original n-dimensional data points X(t) in the Euclidean space^18–24^. *S_k_* is an n-dimensional vector with coordinates of the discrete state *k* and 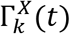 is the probability that X(t) belongs to this discrete state (referred to also as a “cluster k” or a “box k” in the following).

To the resulting expression measuring the approximation error we can add another quality measure, for example the Φ_S_(S) (that measures a quality of discrete states S) and the Φ_Γ_(Γ^X^), measuring the quality of Γ^X^ For example, persistence of the obtained discretization can be controlled by 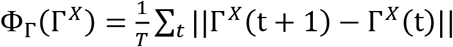^30–32^, whereas Φ_S_(S) can be chosen as a discrepancy between the actual S and some a priori available knowledge about it^28,29^.Then, the best possible probabilistic approximation can be approached by a minimization of the following quality function L with respect to the variables S and Γ^X^:

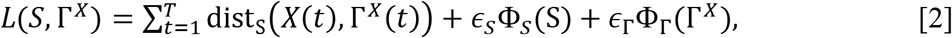

subject to the constraints that enforce that the approximation Γ^X^ is probabilistic

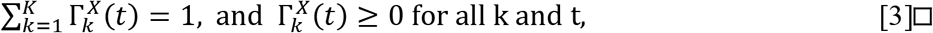

where ϵS, ϵ_Γ_≥0 regulate the relative importance of the quality criteria Φ_S_ and Φ_Γ_ with respect to the approximation quality.

As proven in the Theorem 1 in the Supplement, minima of problem [2-3] can be found in linear time by means of an iterative algorithm alternating optimization for variables Γ^X^ (with fixed S) and for variables S (with fixed Γ^X^). In the following we provide a summary of the most important properties of this algorithm. Detailed mathematical proofs of these properties can be found in the Theorems 1-3 (as well as in the Lemma 1-15 and in the Corollaries 1-11) from the Supplement.

In terms of cost, it can be shown that the computational time of the average iteration for the proposed algorithm grows linearly with the size T of the available data statistics in X – if Φ_Γ_(Γ^X^) is an additively separable function (meaning that it can be represented as 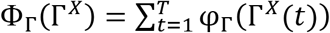. We will refer to the iterative methods for minimization of [2-3] - satisfying this property - as Scalable Probabilistic Approximations (SPA). Further, if the distance metrics distS(X(t), Γ^X^(t)) is either an Euclidean distance or a Kullback-Leibler divergence, then the overall iteration cost of SPA grows as O(nKT) (where n is system’s original dimension and K is the number of discrete states). In another words, computational cost scaling of SPA is the same as the cost scaling of the computationally-cheap K-means clustering^16^(please see a Corollary 6 in the Supplement for a proof). Moreover, in such a case it can be shown that the amount of communication between the processors in the case of the Euclidean distance distS(X(t), Γ^X^(t)) during one iteration in a parallel implementation of SPA will be independent of the size T of system’s output – and will change proportionally to O(nK) and to the number of the used computational cores. Fig. 2 illustrates these properties and shows a principal scheme of the SPA parallelization.

**Fig. 2.**
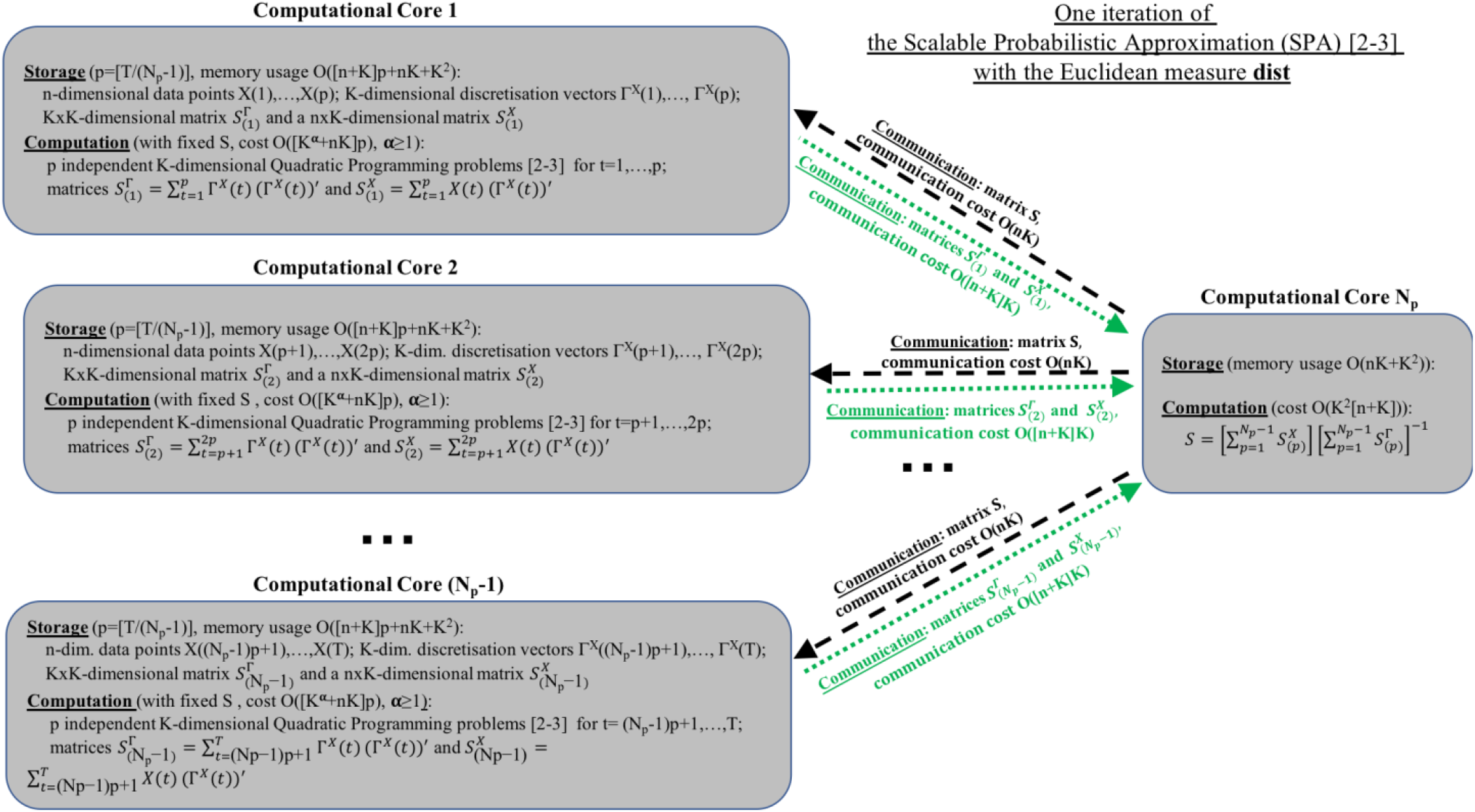
Parallelization of the Scalable Probabilistic Approximation (SPA) algorithm: communication cost of SPA for every channel is independent of the data size T and is linear with respect to the data dimension n.

In terms of quality, it is straightforward to validate that several of the common methods are guaranteed to be sub-optimal when compared to SPA – meaning that they cannot provide approximations better than SPA on the same system’s data X. This can be shown rigorously for example for different forms of K-means^16^ (please see Corollary 1 from the Supplement) and for the different variants of Finite Element clustering Methods on multivariate Autoregressive Processes with external factors (FEM-VARX, in the Corollary 2 from the Supplement)^30–32^.

Fig. 1 shows a comparison of SPA (blue surfaces) to the most common discretization methods, for a set of artificial benchmark problems of different dimensionality n and size T (please see the Supplement for a detailed description of the benchmarks). In comparison with K-means, these numerical experiments illustrate that SPA has the same overall cost scaling (Fig.1b), combined with the significantly better approximation quality and parallelizability scalings (Fig.1a and Fig.1c).

## Computing optimal discretization for Bayesian and Markovian models

Common fitting of Bayesian or Markovian models [1] relies on the availability of discrete probabilistic representations Γ^Y^(t) and Γ^X^(t) – and requires prior separate discretization of X and Y. There is no guarantee that providing any two of such discretization Γ^Y^(t) and Γ^X^(t) as an input for any of the common computational methods^7–15^ for Λ identification would result in an optimal model [1]. In another words, Bayesian and Markovian models obtained with common methods^7–15^ are only optimal for a particular choice of the underlying discrete representations Γ^Y^(t) and Γ^X^(t) (that are assumed to be given and fixed for these methods) – and are not generally optimal with respect to the change of these discretization.

As proven in the Theorem 2 from the Supplement, optimal discretization of the continuous variables X and Y for the model [1] can be obtained jointly, from the family of SPA solutions by minimizing the function *L* [2-3] for the transformed variable 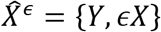. This variable is built as a concatenation (a merge) of the original variables *Y* and *∊X* (where *X* is multiplied with a tunable scalar parameter *∊* > 0). For any combination of parameter *∊* and the discrete dimension K in some pre-defined range, this SPA-optimization [2-3] is performed with respect to the transformed variables 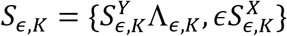 and 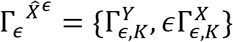.

Then, the optimal combination of *∊* and K – and the optimal discretization for the models [1] - can be found applying standard model selection criteria^34^ (for example, using information criteria or approaches like multiple cross-validation) to the obtained set of solutions 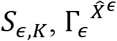.

## Sensitivity analysis and feature selection with SPA

After the discretization problem is solved, an optimal discrete representation Γ^X^(t) can be computed for any continuous point X(t). Obtained vector Γ^*X*(*t*)^ contains K probabilities 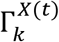 for a point X(t) to belong to each particular discrete state S_k_ – and allows to compute the reconstruction X^rec^(t) of the point X(t) as X^rec^(t)=SΓ^X^(t). In this sense, procedure [2-3] can be understood as the process of finding an optimal discrete probabilistic data compression, such that the average data reconstruction error (measured as a distance between X(t) and X^rec^(t)) is minimized.

In the following, we will refer to the particular dimensions of X as features – and consider a problem of identifying sets of features that are most relevant for the discretization. Importance of any feature/dimension j of X for the fixed discrete states S can be measured as an average sensitivity of the obtained continuous data reconstructions X^rec^(t) with respect to variations of the original data X(t) along this dimension j. For example, it can be measured by means of the average derivative norm 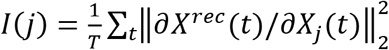. For every dimension j of X(t), this quantity *I*(*j*) probes an average impact of changes in the dimension j of X(t) on the resulting data reconstructions X^rec^(t). Dimensions j that have the highest impact on discretization will have the highest values of *I*(*j*), whereas the dimensions j that are irrelevant for assigning to discrete states will have *I*(*j*) close to zero.

At a first glance, direct computation of the sensitivities *I*(*j*) could seem to be too expensive for realistic applications with large data statistics size T and in high problem dimensions, also due to the a priori unknown smoothness of the derivates 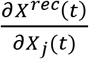 in the multidimensional space of features. However, as proven in the Theorem 3 from the Supplement, in the case of discretizations obtained by solving the problem [2-3] for the Euclidean distance measure *dist*, respective derivatives 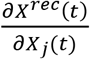 are always piecewise-constant functions of *X_j_*(*t*) if the statistics size T is sufficiently large. This nice property of derivatives allows a straightforward numerical computation of *I*(*j*) for *K* >2 – and an exact analytical computation of *I* for *K* = 2. It turns out that for *K* = 2 the importance of every original data dimension j can be directly measured as 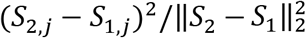. In another words, discretization sensitivity *I*(*j*) for the feature *j* is proportional to the squared difference between the discretization box coordinates *S*_1,*j*_ and *S*_2,*j*_ in this dimension *j*. The smaller the difference between the cluster coordinates in this dimension – the less is the impact of this particular feature *j* on the overall discretization.

It is straightforward to verify (please see Corollary 9 and Theorem 3 in the Supplement for a proof) that the feature sensitivity function *I* = ∑_*j*_ *I*(*j*) has a quadratic upper bound *I* ≤ ∑_*j*,*k*_1_,*k*_2__(*S*_*k*_1__(*j*) − *S*_*k*_2__(*j*))^2^. Setting Φ_*S*_(S) in [2] as Φ_*S*_(S) = ∑_*j*,*k*_1_,*k*_2__(*S*_*k*_1__(*j*) − *S*_*k*_2__(*j*))^2^, for any given combination of integer K and scalar *∊_S_* ≥ 0, minimizing [2-3] would then result in a joint simultaneous and scalable solution of the optimal discretization and feature selection problems. Overall numerical cost of this procedure will be again O(nKT). Changing *∊_S_* will control the number of features: the larger is *∊_S_* the fewer features (i.e., particular dimensions of the original data vector X) will be remaining relevant in the obtained discretization. Optimal value of *∊_S_* can again be determined by means of standard model validation criteria^34^. In the SPA results from Fig. 3 and Fig. 4 (blue curves) we use this form of Φ_*S*_(S) and deploy the multiple cross-validation – a standard model selection approach from machine learning – to determine the optimal *∊_S_* and an optimal subset of relevant features for any given number K of discrete states (clusters).

**Fig. 3.**
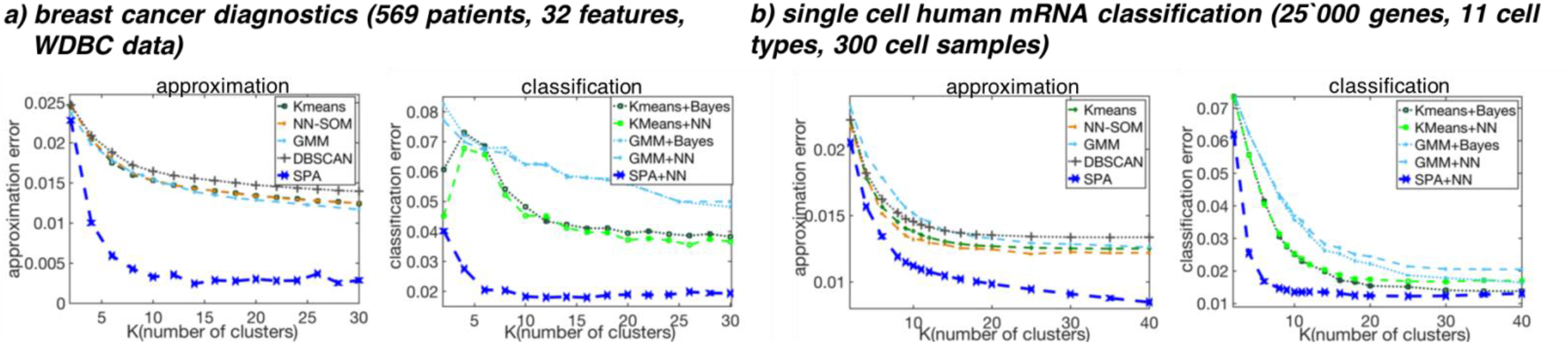
<u>Classification problems</u>: comparing approximation and classification performances of SPA (blue curves) to the common methods on biomedical applications^36,37^. Common methods include K-means clustering (dotted lines), Self-Organising Maps (SOM, brown), pattern recognition Neuronal Networks (NNs, dashed), Gaussian Mixture Models (GMMs, cyan), density-based DBSCAN clustering (dotted with circles) and Bayesian models [1] (Bayes, dotted lines). Approximation error is measured as the multiply cross-validated average squared *Euclidean norm* of difference between the true and the discretized representations for validation data, classification error is measured as the multiply cross-validated average *Total Variation norm (TV)* between the true and the predicted classifications for validation data.

**Fig. 4.**
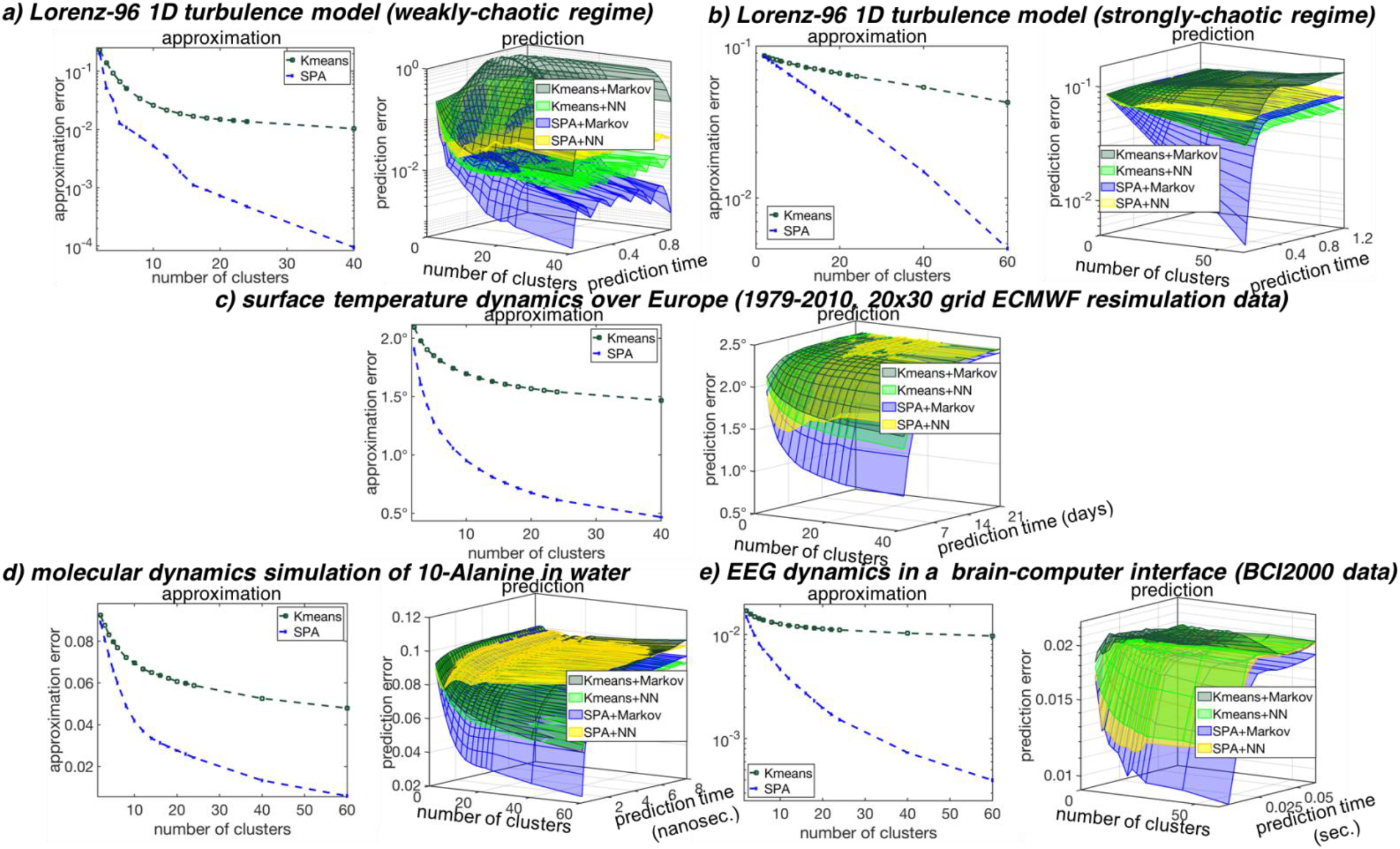
**<u>Prediction problems in time series analysis</u>: comparing approximation and prediction performances of SPA (blue curves) to the common methods on open-source data sets**^38,39,15,40^: common methods include K-means clustering (dark green) in combinations with pattern recognition Neuronal Networks (yellow and light green) and Markov models [1] (dark green). Approximation and the prediction errors are measured in the average squared *Euclidean norm* of deviations between the true and the predicted system states for the validation data not used in the model fitting.

## Applications to classification and prediction problems from natural sciences

Next, we compare the discretization performance of SPA by comparing its approximation errors to the approximation errors of the common methods, including such hard-clustering methods as K-means^16^, soft clustering methods based on Bayesian mixture models^17^ (like Gaussian Mixture Models), density-based clustering^43^ (DBSCAN) and neuronal network discretization methods (Self-Organising Maps)^33^. To compare the performances of these methods, obtained discretizations are used in parametrization of the Bayesian/Markovian models [1] – as well as in parametrization of neuronal networks^33^ - on several classification and time series analysis problems from different areas. To prevent overfitting, we deploy the same multiple-cross validation protocol^34,35^ adopted in machine learning for all of the tested methods. Hereby, the data is randomly subdivided into the training set (75% of the data) where the discretization and classification/prediction models are trained – and performance quality measures (approximation, classification and prediction errors) are then measured on the remaining 25% of validation data (not used in the training). For each of the methods this procedure of random data subdivision, training and validation is repeated 100 times, Fig. 3 and Fig. 4 provide the resulting average performance curves for each of the tested methods. MATLAB scripts reproducing these results are available in a repository *SPA* at *https://github.com/SusanneGerber*. Fig. 3 shows a comparison of approximation and classification performances for two problems of labelled data analysis from biomedicine and bioinformatics: (a) for a problem of breast cancer diagnostics based on X-ray image analysis^36^, and (b) for a problem of single cell human mRNA classification^37^. In these problems variable X(t) is continuous (and real-valued) set of collected features that have to be brought in relation to the discrete set of labels Y(t). In the case of the breast cancer diagnostics example^36^ (a), index t denotes patients and goes from 1 to 569, X(t) contains 32 image features and Y(t) can take two values ‘benign’ or ‘malignant’. In the case of the single cell human mRNA classification^37^ (b), index t goes from 1 to 300 (there are 300 single cell probes), X(t) contains genetic expression levels for 25’000 genes and Y(t) is a label denoting one of the 11 cell types (e.g., ‘blood cell’, ‘glia cell’, etc.).

Fig. 4 summarizes results for five benchmark problems from time series analysis and prediction: for the Lorenz-96 benchmark system^38^ modeling turbulent behavior in 1D, in a weakly-chaotic (a) and in the strongly-chaotic (b) regimes; (c) for the dynamics of historical surface temperatures over Europe^39^, provided by the European Centre for Medium-Range Weather Forecasts (ECMWF); for the biomolecular dynamics of a 10-alanine peptide molecule in water^15^; and for the electrical activity of the brain measured in various Brain-Computer Interaction (BCI) regimes obtained with the 64 channel Electroencephalograph and provided for open access by the BCI2000-consortium^40^.

As can be seen from the Fig. 3 and 4, application of popular discretization methods achieve quality plateaus for all of the considered applications. In another words, increasing the number K of discrete states (clusters) – and increasing the overall computational cost - does not improve the resulting performance accordingly. In contrast, application of methods involving SPA results in drastically improved performances – with a performance improvement factor ranging from two to four (for breast cancer diagnostics example, for single cell mRNA classification, for the temperature data over Europe and for the molecular dynamics application). For the Lorenz-96 turbulence applications^38^ and for the brain activity application^40^, discretization obtained by SPA are ten to hundred times better than the discretization from common methods – being at the same level of computational cost as the popular K-means clustering.

Evaluating a prediction performance of different models for a particular system, it is important to compare it with the trivial prediction strategies called *mean-value prediction* and *persistent prediction*. The *mean-value prediction* strategy predicts the next state of the system to be an expectation value over the previous already observed states – and is an optimal prediction strategy for stationary independent and identically distributed processes like the Gaussian process. The *persistent prediction* strategy is predicting the next state of the system to be the same as its current state: this strategy is particularly successful and is difficult to be beaten for the systems with more smooth observational time series, like for example for the intraday surface temperature dynamics. As it can be seen from the Fig. S3 from the Supplement, among all other considered methods (K-means, neuronal networks, SOM, mixture models) only the SPA discretization combined with the Markov models [1] allow outperforming both the *mean-value* and the *persistent predictions* for all of the considered systems.

## Summary

Computational cost becomes a limiting factor when dealing with big systems. An exponential growth in the hardware performance observed over the last 60 years (the Moore’s law) is expected to come to an end in the early 2020’s^41^. More advanced machine learning approaches (e.g., neuronal networks) exhibit the cost scaling that grows polynomial with the dimension and with the size of the statistics – making some form of ad hoc preprocessing and pre-reduction with more simple approaches (e.g., clustering methods) unavoidable for the big data situations. However, such ad hoc pre-processing steps might impose a significant bias that is not easy to quantify. At the same time, lover cost of the method typically goes hand-in-hand with the lover quality of the obtained data representations (see Fig.1). Since the amounts of collected data in most of the natural sciences are expected to continue their exponential growth in the foreseeable future, a pressure on a computational performance (quality) and a scaling (cost) of algorithms will increase.

Instead of solving discretization, feature selection and prediction problems separately, the introduced computational procedure (a Scalable Probabilistic Approximation, or SPA) solves them simultaneously. The iteration complexity of SPA scales linearly with data size. The amount of communication between processors in the parallel implementation is independent of the data size and is linear with the data dimension – making it appropriate for big data applications. Hence, SPA did not require any form of data pre-reduction for any of the considered applications. As demonstrated in the Fig. 1, having essentially the same computational cost scaling as the very popular and computationally very cheap K-means algorithm^16–17^, SPA allows achieving significantly higher approximation quality and a much higher parallel speed-up with the growing size T of the data.

Applications to large benchmark systems from natural sciences (Fig. 3 and 4) reveal that these features of SPA allow a drastic improvement of approximation and prediction qualities – combined with a massive reduction of computational cost. For example, computing the next-day surface temperature predictions for Europe (e.g., at the European Centre for Medium-Range Weather Forecasts, ECMWF) currently relies on solving equations of atmosphere motion numerically - performed on the supercomputers^39^. Discretization and prediction results for the same online daily temperature data provided in the Fig. 4c were obtained on a standard Mac PC, exhibiting a cross-validated mean error of 0.75 degree Celsius for the one-day-ahead surface air temperature predictions (approximately 40% smaller than the current next-day temperature prediction errors by weather services).

Such probability-preserving and stable predictions Γ^Y^(t) can be done very cheaply with the Bayesian or Markovian model [1] from the available SPA discretization [2,3] – just by computing the product of the obtained Bayesian matrix Λ with the discretization vector Γ^X^(t). The cost of this whole prediction operation scales linearly – resulting in orders of magnitude speed-up as compared to the predictions based on the whole system’s simulations. These results indicate a potential to pave the ways to massively-parallel data-driven and error-controlled descriptive models for a robust automated classification and prediction in complex systems.

## Acknowledgments

We thank Giovanni Ciccotti (La Sapienza Rome), Martin Weiser (ZIB Berlin), David L. Donoho (U Stanford), Michael Wand (JGU Mainz) and Patrick Gagliardinin (USI Lugano) for helpful comments about the manuscript.

## Author contributions

SG and IH have designed research, wrote the main manuscript and produced results in the Fig. 3 and Fig. 4; LP and IH have produced results in the Fig. 1 and proven Theorems 1-3 in the Supplement; MN has prepared the data and participated in the analysis of single cell mRNA data (Fig. 3b).

## Data and materials availability

The description of the model systems used for Fig. 1 is provided in the Supplement. Data sets used for computations in the Fig. 3 and Fig. 4 were published^36,37,38,39,15,40^ and are also available in an open access in the directory *Data* of the repository *SPA* at *https://github.com/SusanneGerber*

## Model and algorithms availability

We used the standard MATLAB functions *kmeans()*, *fitgmdist()*, *patternnet()* and *som()* to compute the results of the common methods (K-means, GMM, NN and SOM) in the Figs.1+3–4. To avoid getting trapped in the local optima and to enable a unified comparison, of all methods we used 10 random initializations and selected the results with the best approximation quality measure for the training sets. In case of the pattern recognition neuronal networks (NN, evaluated in the classification and prediction performance subfigures of the Fig. 3–4), in each of the instances of the multiple cross-validation procedure we repeated the network fitting for the numbers of neurons in the hidden layer ranging between 1 and 15 and selected the results with the best classification/prediction performances for the training set. The left-stochastic discretization algorithm (LSD) from Fig. 1 was implemented by us in MATLAB according to the literature description^24^ and is provided for an open access. SPA algorithms developed and used during the current study are also available in an open access as MATLAB code at *https://github.com/SusanneGerber*

- **Description of the synthetic data problems**
- **General Scalable Probabilistic Approximation (SPA) formulation**

<u>Lemma 1</u> - SPA algorithm generates nonincreasing objective function
<u>Lemma 2</u> - sufficient condition for solvability of S subproblem
<u>Lemma 3</u> - sufficient condition for solvability of Γ subproblem
<u>Lemma 4</u> - about separability
<u>Theorem 1</u> - properties of SPA algorithm
<u>Corollary 1</u> - suboptimality of K-means
<u>Corollary 2</u> - suboptimality of FEM-BV and FEM-H1
- **SPA in the Euclidean space**

<u>Lemma 5</u> - non-unique solution of SPA_2_

∘ **Optimality conditions**
∘ **The solution of** *S* **subproblem**
<u>Lemma 6</u> - analytical solution of *S*-problem
<u>Lemma 7</u> - computational and memory complexity of *S*-problem
<u>Corollary 3</u> - computational and memory complexity of *S*-problem in K-means
<u>Lemma 8</u> - regularization of *S*-problem
<u>Lemma 9</u> - uniqueness of reconstruction with fixed Γ
<u>Lemma 10</u> - derivative of solution with fixed Γ
<u>Corollary 4</u> - stability of solution in K-means

∘ **The solution of** Γ **subproblem**

<u>Lemma 11</u> - about separability of QP problems
<u>Lemma 12</u> - computational and memory complexity of Γ-problem
<u>Corollary 5</u> - computational and memory complexity of Γ-problem in K-means
<u>Lemma 13</u> - complexity of one iteration of (SPA_2_)
<u>Corollary 6</u> - comparison of leading order complexity scalings for K-means and for (SPA_2_)
<u>Lemma 14</u> - Γ solution is continuous piecewise linear function in *X*
<u>Corollary 7</u> - derivative of reconstruction is continuous piecewise constant
<u>Lemma 15</u> - analytical solution for *K* = 2
<u>Lemma 16</u> - uniqueness of reconstruction with fixed *S*
- **Computing optimal discretisations for Bayesian and Markovian models**

<u>Theorem 2</u> - the combination of optimal discretization with Markov model
- **Feature selection with SPA in the Euclidean space**

<u>Lemma 17</u> - the estimation of Γ subproblem solution stability
<u>Corollary 8</u> - the consistency of change of reconstruction and original data
<u>Corollary 9</u> - projection onto optimal polytope
<u>Theorem 3</u> - *S* subproblem regularization and feature selection
<u>Corollary 10</u> - connection between regularization and feature selection
<u>Corollary 11</u> - numerical estimation of reconstruction derivative

## Description of the synthetic data problems (used in the Figure 1 of the main manuscript)

This section provides the description of the benchmark, whose results are presented in Manuscript in the Figure 1. For a given number of data points *T* > 0 and a data dimension (number of features) *n* ≥ 2, we generate the random data *X* = [*x*_1_,…,*x_T_*] ∈ ℝ^*n,T*^ from multivariate normal distribution with different parameters based on a predefined cluster affiliation.

We choose the cluster affiliation in such a way, that the number of points affiliated to cluters *T_k_* is approximately the same along the clusters, i.e.,

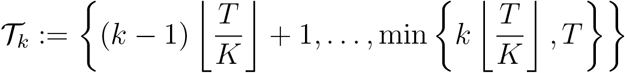

denotes the set of point indexes affiliated to *k*-th cluster. Please, notice that these sets are disjoint and union of them forms the set of all point indexes {1,…,*T*}. Using this decomposition, we generate corresponding data points for every cluster *k* = 1,…, *K* as random realisations from the multivariate normal distributions

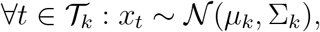

where *μ_k_* ∈ ℝ^*n*^ denotes the mean value and Σ_*k*_ ∈ ℝ^*n,n*^ a covariance matrix.

In our benchmark, we choose *K* = 4 with parameters

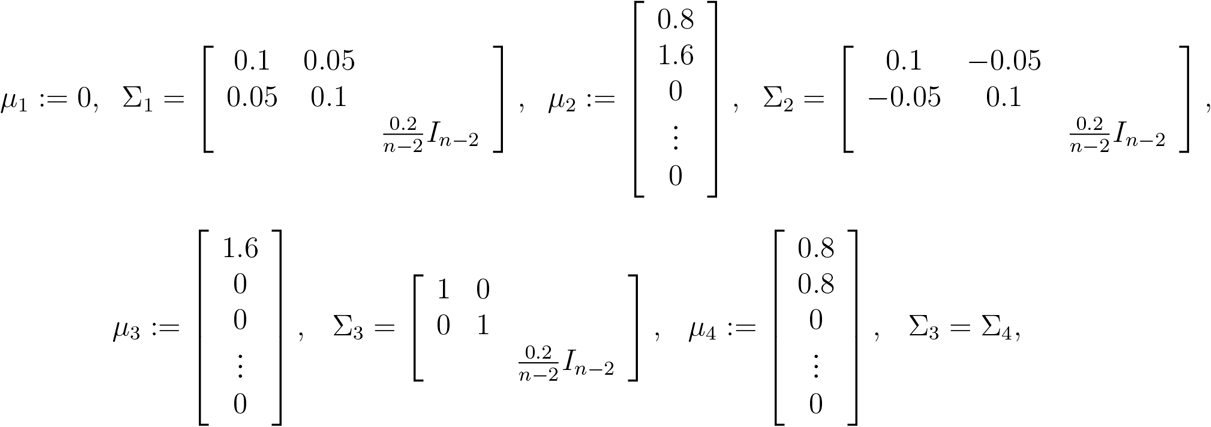

where *I*_*n*−2_ ∈ ℝ^*n*−2,*n*−2^ is identity matrix.

## General Scalable Probabilistic Approximation (SPA) formulation

The SPA optimization problem is given by

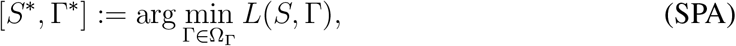

where

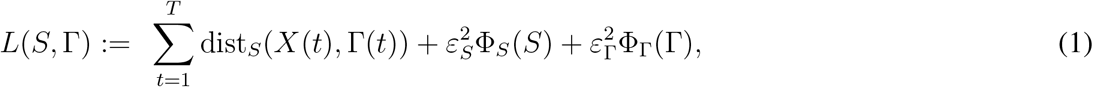

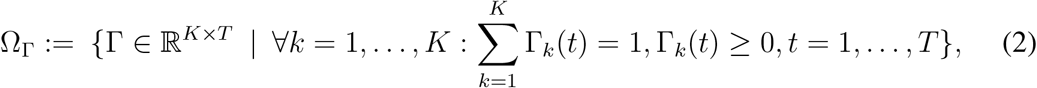

*T* denotes the number of data points, 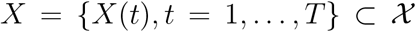 are given data from space 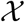 deployed with the norm ||·||, *K* > 1 denotes the number of discrete states (clusters), Γ = {Γ_*k*_(*t*), *k* = 1,…, *K*, *t* = 1,…, *T*} ⊂ Ω_Γ_ ⊂ ℝ^*K*×*T*^ are unknown cluster affiliation probability vectors, and 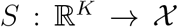 are unknown data representation vectors. We include the possibility of Tikhonov-based regularization of original ill-posed problem using the regularization functions Φ_*S*_, Φ_Γ_ with corresponding regularization parameters *ε_S_, ε*_Γ_ ≥ 0.

### Algorithm 1: General SPA algorithm.

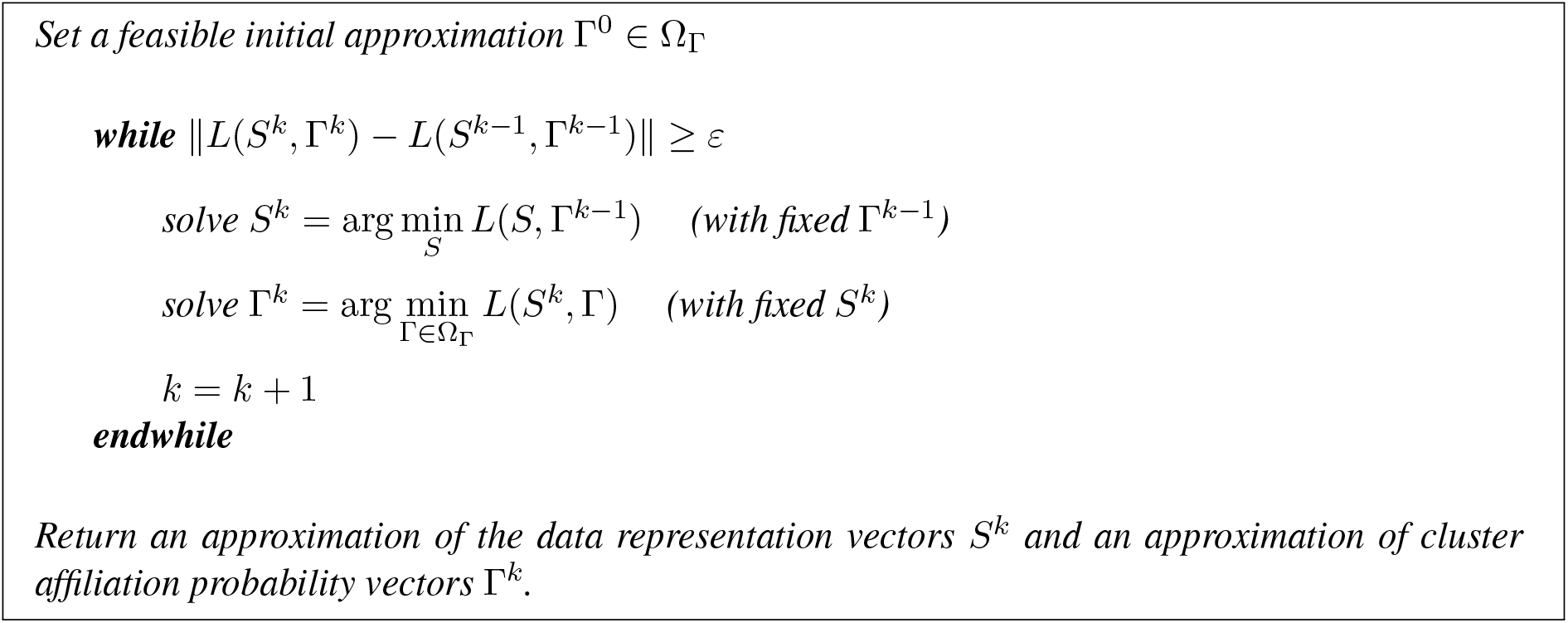

The problem (SPA) can be solved using the Algorithm 1. The idea is based on the construction of the sequence of split optimization problems. The iteration computational complexity of this algorithm is given by the complexity of the computation of inner optimization problems with fixed variables. The algorithm of this type is well-known as coordinate descent method (*10*) or alternating least-squares method (*1*). The following Lemma presents the basic convergence properties of the algorithm.

### Lemma 1.

*If the solutions of inner optimization problems in Algorithm 1 exist, then algorithm generates a sequence of approximations for the optimization problem* (SPA) *with nonincreasing objective function values, i.e*.,

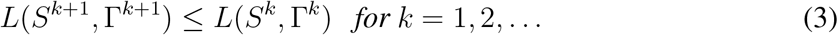

*Proof*. If the solutions of inner optimization problems exist, then the solution process of inner optimization problems provides the approximation with smaller (or the same) function value with respect to non-fixed variable, i.e. (see the Definition 1 in APPENDIX),

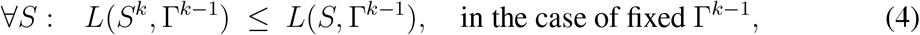

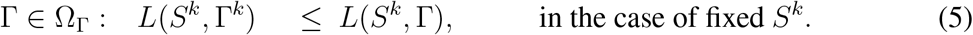

Choosing *S* = *S*^*k*−1^ in (4) and Γ = Γ^*k*−1^ in (5) we get

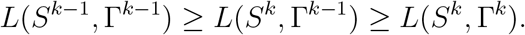

Since the objective function (1) is generally non-convex (but bounded from bellow - each distance function is non-negative), the sequence (3) can possibly converge only to the local optimum. To deal with this non-globality, one has to run the algorithm for several random initial Γ^0^ and choose the solution with the lowest function value. Such a Monte-Carlo-based approach is commonly used for solving the optimization problem with multiple local optimality points and it can be found in literature as annealing steps (*10*).

However, the convergence of the whole process still highly depends on the solvability of inner optimization problems. Following lemmas present the elementary and the most common situations when the solution exists.

### Lemma 2.

*If the distance function dist_S_ and the regularization function* Φ_*S*_ *in* (SPA) *are convex, bounded from below and continuously differentiable with respect to the variable S, then the solution with respect to S exists and can be found using the necessary optimality conditions for unconstrained problems*.

*Proof*. The Lemma is a consequence of optimization theory fundamental results, see for example (*3*).

### Lemma 3.

*If the distance function distS and the regularization function* Φ_Γ_ *in* (SPA) *are continuous in variable* Γ, *then the solution of the problem with respect to* Γ *exists*.

*Proof*. Please notice that feasible set Ω_Γ_ is compact (i.e., closed and bounded) and convex, therefore if *L* is continuous, then the existence of the solution is a consequence of Weierstrass Extreme Value Theorem (*3*).

Typically, the largest dimension parameter of the whole problem (SPA) is the number of data points *T* and the classification data-discretization process (SPA) does not reduce this number. It provides the data representation vectors *S*, whose size is determined by the size of individual data points (the dimension of vector space 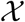) and the number of them is equal to the number of clusters *K*. We can conclude, that the optimization problem with respect to *S* is much smaller in comparison to the optimization problem with respect to the second variable Γ. The unknown Γ consists of cluster affiliation probability vector of each individual data points, i.e., its size is determined by *T* and *K*. Fortunately, the objective function *L*(1) is composed as a sum of local representation errors and therefore if the regularization function Φ_Γ_(Γ) is also additively separable (the case when it consists of the sum of local regularization functions for individual representations) then the whole minimization problem (SPA) is separable. The following Lemma presents the basic property of additively separable optimization problems.

### Lemma 4.

*If L in the optimization problem* (SPA) *is additively separable in t* (*except* Φ_*S*_(*S*)), *i.e., there exist functions L_t_*(*S*,Γ(*t*)), *t* = 1,…, *T such that*

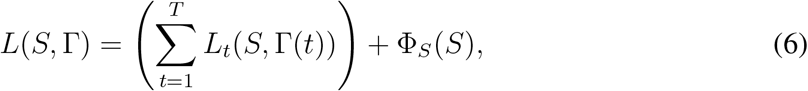

*then the solution of an optimization problem* (SPA) *with fixed S can be composed from solutions of individual problems*

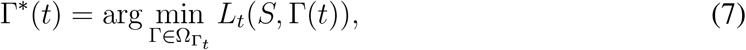

*where*

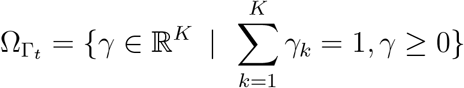

*and* Ω_Γ_1__ × ⋯ × Ω_Γ_*T*__ = Ω_Γ_ *is the decomposition of the feasible set of the original problem* (SPA).

*Proof*. The definition of optimality point of (7) reads as (see the Definition 1 in APPENDIX)

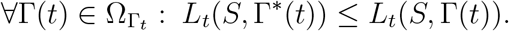

Since this inequality can be formulated for all *t* = 1,…, *T*, we can sum these *T* inequalities to obtain

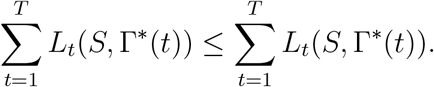

If we add term Φ_*S*_(*S*) (constant in Γ) to both sides of this inequality and use notation (6), we obtain

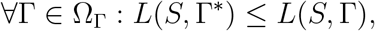

which is a definition of the optimality point of optimization problem (SPA) with respect to Γ.

The separability plays crucial role in the embarassingly parallel computations; one can solve the whole set of *T* optimization problems independently using modern multi-core architectures, see Figure S2. The Γ-problem can be splitted into smaller subsets and distributed onto separated computational nodes, which is a commonly adopted approach when working on supercomputers. Each node solves the given subset of problems without any communication with the other nodes. Moreover, if the node includes multi-core processors, then (again) each core can solve independently the part of the node subproblem. This “embarrasingly-parallel” hierarchical computation of the large-scaled problem can be exploited even more when using modern GPU architectures; in this case, the relativelly small Γ(*t*) problem (of size *K*) can be solved using just one computational thread, i.e., one computational core (please see Fig. 2c in the main manuscript).

It is necessary to mention that if the regularization function Φ_Γ_ is not separable in *T* (for example when enforcing the persistency of regime/cluster in time, see FEM-H1 and FEM-BV methods (*9*)), then the problem is not embarassingly parallel and computational nodes/cores/threads have to communicate during the solution process. However, as was demonstrated in (*12*), one can still utilize Projected Gradient methods since the projection onto separable simplexes Ω_Γ_ is still embarassingly parallel.

The following Theorem summarizes the general properties of the Algorithm (1).

### Theorem 1

(Properties of the SPA algorithm). *Let* 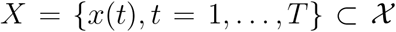 *be given data from space* 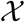, *K* > 1 *is a given number of clusters. Let* dist_*S*_, Φ_*S*_, Φ_Γ_ *be such functions that L*(*S*, Γ) *in* (SPA) *is convex, bounded from bellow and continuously differentiable with respect to the variable S and continuous in the variable* Γ.

*Then the Algorithm 1*

a. *(a) is generating a monotonically non-increasing sequence*.

*Moreover, if L*(*S*, Γ) *is additively separable problem in* Γ, *then the Algorithm 1*

*(b) scales linearly in the size T of the data statistics X*,
*(c) requires the amount of communication independent of the data size*.

*Proof*. (a) is a consequence of Lemma 2, Lemma 3, and Lemma 1. To prove (b) and (c), please notice that the solution of optimization problem with respect to *S* is independent of the number of provided data points *T*. If the assumption of separability is fullfilled, then in the case of solving the problem with respect to Γ, we can using Lemma 4 reformulate the original problem as a set of *T* independent problems, whose dimension is (again) independent of *T*.

Let us present the connection between SPA and some of the commonly used discretization (clustering) methods in following Corollaries.

### Corollary 1

(Suboptimality of K-means). *Measured in terms of squared Euclidean distance, discretisations providided by K-means are always suboptimal with respect to the discretisations obtained with* (SPA).

*Proof*. Let us consider data *X* ∈ ℝ^*n*,T^. The aim of the K-means clustering algorithm (**?**) is to optimally partition given data into *K* disjoint clusters based on the Euclidean distance from (unknown) optimal centroids of the clusters. The algorithm computes these cluster centroids *S_k_* ∈ ℝ^*n*^ and binary affiliation Γ ∈ {0,1}^*K,T*^, where Γ_*k,t*_ = 1 if *x_t_* belongs to *k*-th cluster and Γ_*k,t*_ = 0 otherwise. The corresponding optimization problem is formulated as

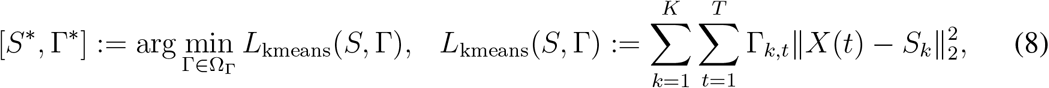

where Ω_Γ_ ⊂ {0,1}^*K,T*^ includes the condition for strict affiliation of a point into exactly one cluster, i.e.,

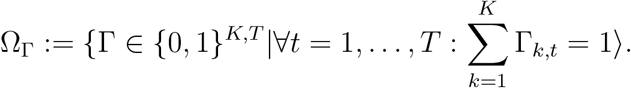

The problem (8) is solved iteratively; the feasible initial approximation of affiliations Γ is chosen randomly (the points are randomly affiliated to clusters) and afterwards, the iterative procedure solves consecutively the problems with one fixed variable. In the case of K-means, both of the subproblems have analytical solutions

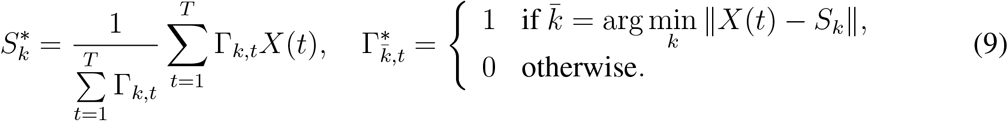

In fact, the scheme of the algorithm is the same as in the Algorithm 1 and one can easily check that if Γ is binary variable and we choose 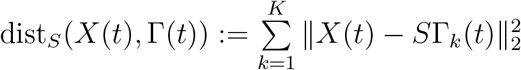 in (SPA) (in following text denoted as (SPA_2_)) then

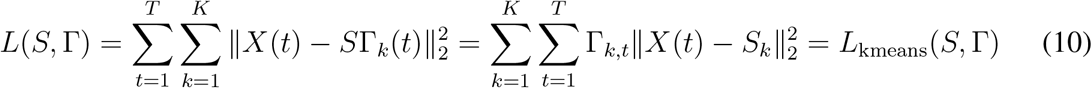

and therefore *K*-means algorithm is equivalent to (SPA_2_).

The variant of K-means algorithm with relaxed binary condition is well-known as soft K-means algorithm (**?**). In this case, Γ_*k,t*_ represents the probability that *X*(*t*) is affiliated to the *k*-th cluster. The feasible set Ω_Γ_ enforces the rows of Γ to be a corresponding discrete probability density vector, i.e., each element is continuous variable from [0, 1] and because of the law of the total probability, the sum of the elements of this vector has to be equal to one. One can easily check that Ω_Γ_ defined by (2) represents these conditions. However in the case of continuous Γ, the equality (10) does not hold. Using the Jensen’s inequality (*10*) we get

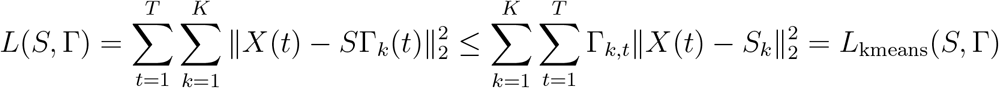

and therefore soft K-means algorithm produces only the upper estimation of the (SPA_2_) optimization problem.

### Corollary 2

(Suboptimality of FEM-BV and FEM-H1.). *Measured in terms of a squared Euclidean distance, discretisations providided by FEM-BV and FEM-H1 methods are always sub-optimal with respect to the discretisations obtained with* (SPA).

*Proof*. The family of FEM-BV and FEM-H1 methods consists of methods used for time series analysis (*9*), (*12*). The idea is to extend stationary models with clustering and additional time regularization for enforcing the model time persistency.

In time series modelling, we suppose that the measured data *x*_1_, *x*_2_,…, *x_T_* ∈ ℝ^*n*^ are described by the parametric model *ψ* and include the additive noise, i.e.,

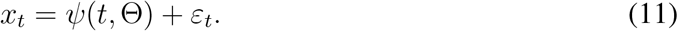

For instance one can consider autoregessive models, e.g., the Var-X model defined as

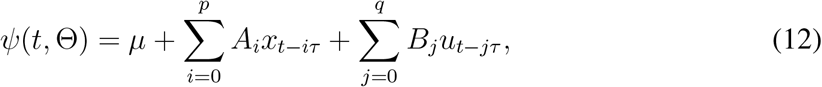

where Θ = (*μ, A*_0_,…, *A_p_*, *B*_0_,…, *B_q_*) includes all model parameters, *τ* > 0 is a discretisation time step, *p*, *q* ≥ 0 represent the size of memory, and *u_t_* denote the external factors or controls. The aim of the analysis is to find parameters of the model which fit the given data *x_t_*, *u_t_* in an optimal way, for example, one can utilize minimum least square error to formulate optimization problem

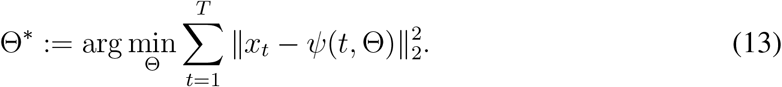

In the case of Var-X model (12) the optimization problem (13) is unconstrained quadratic programming problem and the necessary optimality conditions formulate the corresponding system of linear equations which has to be solved.

FEM-BV and FEM-H1 belong to the non-stationary models; here we suppose that the parameters of model Θ are non-stationary, i.e., they are changing (can change) in time. In general, non-stationary model without any additional assumptions, e.g., restriction of the set of permissible parameters, lead to ill-posed and biased results. In the case of FEM-BV and FEM-H1, we include the assumption of the time persistency of model parameters introducing the finite number of regimes (i.e., clusters) in which the model parameters are stationary. The switching between those regimes is realized by a hidden regime-switching process, which describes the activity of each regime in a given time. For example, if we consider stationary Var-X model (12) on each of the *K* regimes, then the corresponding optimization problem is formulated as

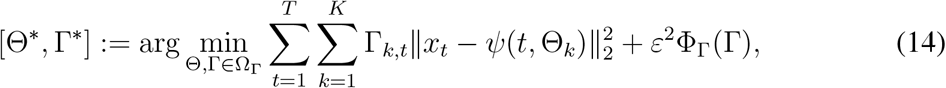

where Θ = [Θ1,…,ΘK] includes (unknown) parameters of local models on regimes and Γ_*k*_,: are model indicator functions defined in similar as in the case of **K**-means, i.e., Γ_*k,t*_ = 1 if the time series in time *t* is in *k*-th regime and and Γ_*k,t*_ = 0 otherwise. Regularization function Φ_Γ_(Γ) with regularization parameter *ε*^2^ ≥ 0 enforces the time persistency of a regime-switching process. In the case of FEM-BV, we consider Bounded variation (BV) norm defined as

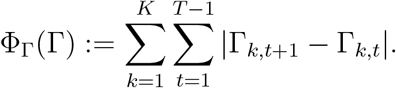

If we consider binary Γ then this value is equal to the number of switches between regimes and the regularization by this function decreases the global number of switches in the solution. The optimization problem (14) is solved using Algorithm 1, however, in this case the Γ subproblem is not separable due to non-separable regularization term and this problem of dimension *KT* has to be solved using linear programming algorithm. For extended details on the method see (*9*).

It is straightforward to verify that the formulation of FEM-BV corresponds to (SPA) with distance function defined as a local Euclidean distance between given data *X*(*t*) and the local value of model *ψ*

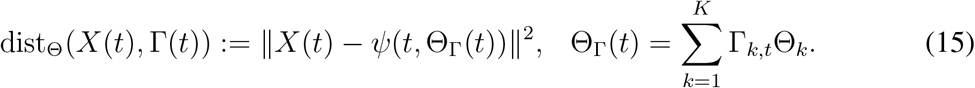

Similarly to the soft K-means clustering case considered in the Corollary 1 above, we can relax the hard clustering property (i.e., the property that each data point is exclusively affiliated to exactly one regime) considering Γ_*k,t*_ to be probability of affiliation of *X*(*t*) to *k*-th regime.

Each Γ_:,*t*_ forms the discrete probability density vector of affiliation of *X*(*t*) to regimes and a corresponding feasible set is given by (2). To include the assumption of time persistency, one can adopt the H1 half-norm

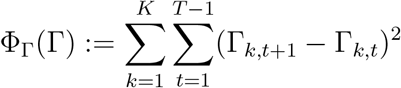

to get the FEM-H1 method, see (*9*). The problem is solved by an Algorithm 1, the corresponding Γ subproblem is non-separable convex quadratic programming problem of size *KT*, see (*12*).

Please notice that Θ depends linearly on variable Γ, the Var-X model depends linearly on parameters Θ, and the distance function dist_Θ_ is convex in variable *ψ*. Summarizing these properties we can state that distance function is convex in Γ (see (*3*) for the list of operations which preserve convexity). Using the Jensen’s inequality we get

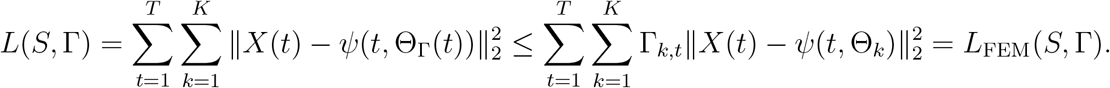

This inequality holds also when we add any regularization Φ_Γ_(Γ) to the both sides. Hence, FEM-BVandFEM-H1 algorithms produce only the upper estimation of the (SPA) optimization problem with a corresponding choice of distance function and regularization.

## SPA in the Euclidean space

We consider the data from real *n*-dimensional vector space 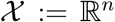 and Euclidean distance measure on 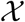 defined by

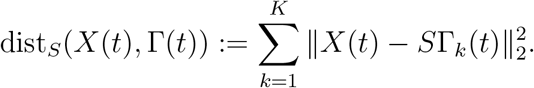

For simplicity, we compose the vectors into matrices

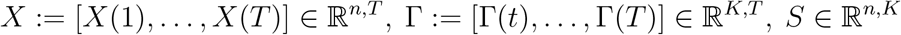

and afterwards, the corresponding optimization problem (SPA) without regularization can be written in a form

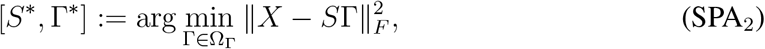

where *F* denotes Frobenius norm and the feasible set is defined by

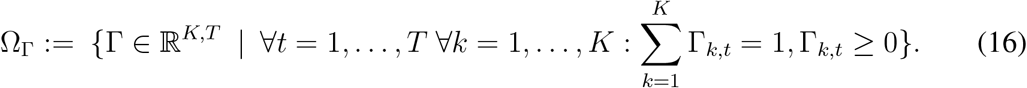

### Lemma 5.

*The solutions of problem* (SPA_2_) *are always non-unique for any K* > 1.

*Proof*. Let us consider an arbitrary solution [*S*\*, Γ*] and nonsingular matrix *R* ∈ ℝ^*K,K*^, *R* ≠ *I_K,K_* such that *R*Γ ∈ Ω_Γ_. Such a matrix always exists, e.g., we can consider a permutation matrix which permutes the rows of Γ, i.e., the indexes of clusters. Since we can write

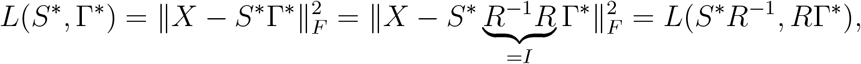

we can state that feasible [*S*\**R*^−1^, *R*Γ*] ≠ [*S*\*, Γ*] has the same (minimal) function value and therefore it also solves the problem.

## Optimality conditions

We define the Lagrange function (*10*) corresponding to the optimization problem (SPA_2_) by

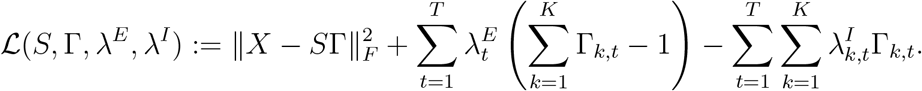

Here λ^*E*^ ∈ ℝ^*T*^ are Lagrange multipliers corresponding to equality constraints defined by the feasible set (16) and λ^*I*^ ∈ ℝ^*K,T*^ denotes the Lagrange multipliers corresponding to the nonnegativity bound constraints in (16).

The full system of Karush-Kuhn-Tucker (KKT) optimality conditions for this system will be:

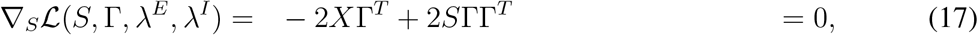

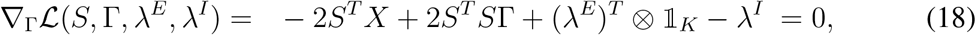

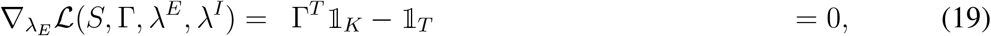

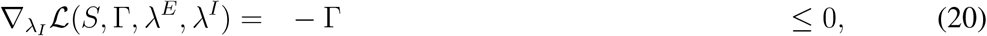

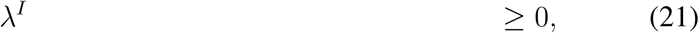

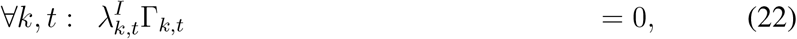

where 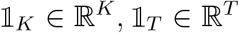 denotes the vectors of ones. Equations (17) and (18) are first-order optimality conditions, equation (19) and inequality (20) are constraints given by the definition of the feasible set (16), inequality (21) preserves the non-negativity of inequality Lagrange multipliers, and equations (22) represent the so-called complementarity conditions for inequality constraints.

## The solution of *S* subproblem

### Lemma 6

(The solution of *S*-problem). *Let* Γ ∈ Ω_Γ_ *in problem* (SPA_2_) *be fixed. Then the system of all solutions of optimization problem* (SPA_2_) *with respect to S is given by*

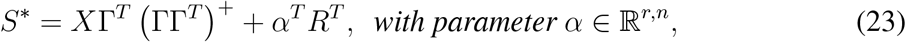

*where* (ΓΓ^*T*^)^+^ ∈ ℝ^*K,K*^ *denotes a pseudoinverse*^1^ *of the matrix* ΓΓ^*T*^, *R* ∈ ℝ^*K,r*^ *is a matrix whose columns form the basis of the null space of* Γ^*T*^, *i.e*.

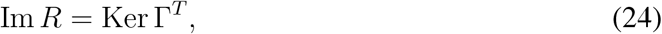

*and r* = dim Ker Γ^T^ *denotes the nullity of matrix* Γ^*T*^.

*Proof*. Please notice that the objective function of (SPA_2_) in terms of variable *S* is continuously differentiable convex matrix quadratic function. The necessary optimality condition of given unconstrained optimization problem is given by (17). This system of linear equations with multiple right-hand side vectors with symmetric positive semi-definite system matrix always has a solution. If the system matrix is non-singular, then the unique solution is given by

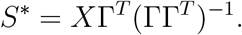

However, the non-singularity of system matrix ΓΓ^*T*^ ∈ ∈^*K,K*^ (and consequently, the existence of inverse matrix) is not guaranteed^2^, the system of all solutions is given by (23) where all solutions differ by the vector from Ker ΓΓ^*T*^, see (*8*) or (*7*).

Next we deal with the eventual ill-posedness of the optimization problem (SPA_2_) in variable *S*, or equivalently, with the ill-posedness of the system of linear equations (17). Deploying Tykhonov-regularization, we reformulate the original (SPA) problem choosing the regularization function

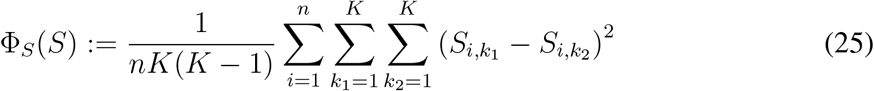

and consider regularization parameter 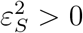. Please notice that the solution of the optimization problem in term of variable *S* is independent on the choice of regularization function Φ_Γ_. The following Lemma proves that (25) guarantees the unique solvability of *S*-problem.

### Lemma 7.

*The computational complexity of solving S subproblem in* (SPA_2_) is 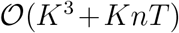, *with the memory complexity of* 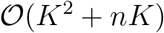.

*Proof*. The first step in solving the *S* subproblem is the assembly of the matrix ΓΓ^*T*^ and of the matrix of the right-hand side vectors *X*Γ^*T*^ in an equation (17). Let us remind that the complexity of computing matrix-matrix multiplication of general (non-sparse) matrices *A* ∈ ℝ^*n,m*^ and *B* ∈ ℝ^*m,p*^ is 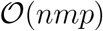, therefore in our case, the overall complexity of assembling the problem is 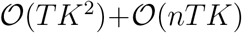. The memory required to store these two new matrices is 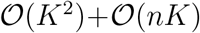.

In general, the direct methods for solving a system of linear equation *Ax* = *b, A* ∈ ℝ^*m,m*^ have the complexity of order 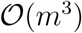. Iterative methods, like Krylov subspace algorithms, are based on the iterations where the computational complexity scaling in the leading order is dominated by the multiplication with a system matrix *A*, which is of order 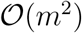. Number of iterations needed for the convergence, when using a suitable preconditioner, is usually much less than 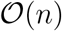. Therefore, the overall work for solving the system of linear equations is less than 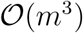. In general, numerical linear algebra algorithms for this purpose are using the auxiliary vectors of dimension ℝ^*m*^, whose number is independent on the dimension of the problem. Therefore, the amount of additional memory used for solving the system of linear equations is of the order 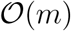.

Applying these general results to *S* subproblem which consists of *T* linear systems of dimension *K*, we obtain the total computational complexity 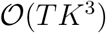 and a memory complexity 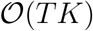. Since the system matrix is the same for all subsystems, therefore one can compute pseudoinverse and use (23) directly, which will lead to the total computational complexity of 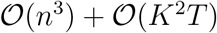. In practial applications the computation of pseudoinverse is typically much slower than solving the system of linear equations.

### Corollary 3.

*In the case of K-means algorithm, evaluation of an analytical solution S*\* (9) *consists of computing two sums with the computational complexity* 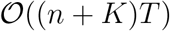. *To compute the sums, one has to use an additional auxiliary vector of dimension* 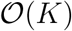.

### Lemma 8

(*S*-problem with regularization). *Let* Γ ∈ Ω_Γ_ *in a problem* (SPA_2_) *with an additional regularization function* (25) *be fixed. Then, for any* 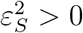 *the problem with respect to S has a unique solution given by*

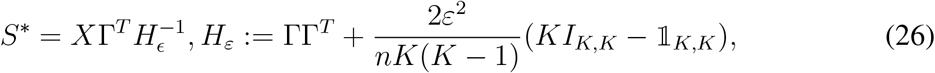

*where I_K,K_* ∈ ℝ^*K,K*^ *is an identity matrix and* 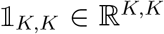 *is a matrix full of ones. Moreover the spectrum of regularized Hessian matrix H_ε_ can be estimated by*

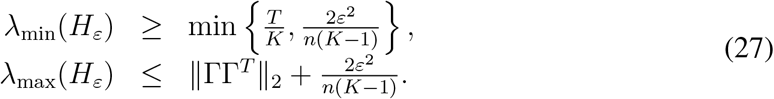

*Proof*. The gradient of the original objective function *L* in (SPA_2_) without regularization is given by the left-hand side of (17). Let us focus on the gradient of regularization function whose components are given by (for every *i* ∈ {1,…, *n*}, *k* ∈ {1,…, *K*})

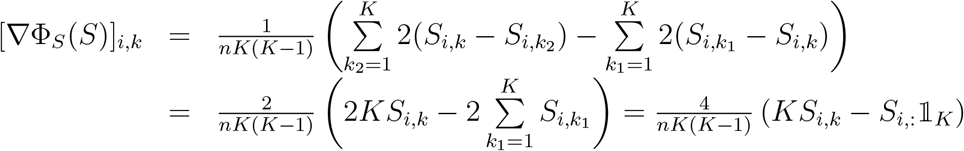

where 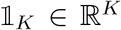 is a column vector of ones. It is easy to see that the whole gradient can be written as

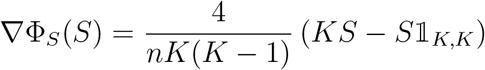

and therefore the necessary optimality condition of the regularized problem is given by the solution of a regularized linear system of equations

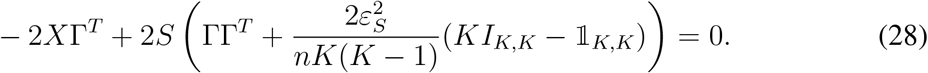

It remains to show that the system matrix is non-singular for any 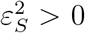 and therefore we will be able to multiply the whole equation with the matrix inverse to obtain a unique solution.

Please notice that the matrix 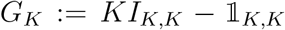 is a Laplacian matrix of a complete graph on *K* nodes, therefore it is symmetric positive semidefinite and 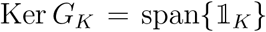, see (*5*).

For the simplicity, let us denote 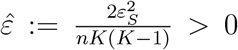. For any non-zero *y* ∈ ℝ^*K*^ we can differentiate two cases

- if *y* ∉ Ker *G_K_* then *y^T^ G_K_y* = *Ky^T^y* (the spectrum of complete graph Laplace matrix is composed from one zero eigenvalue and eigenvalues of value *K* with multiplicity *K* − 1, see (*5*)) and

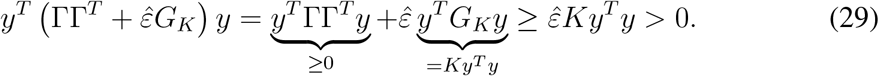
- if *y* ∈ Ker 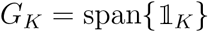 then there exists a non-zero *α* ∈ ℝ such that non-zero *y* can be written as 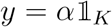. Using the equality constraints of the feasible set Ω_Γ_ (16) written in a form 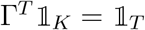 we can state that

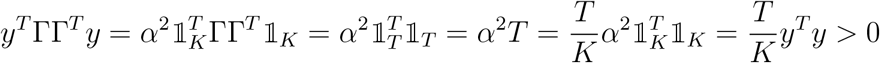

and consequently

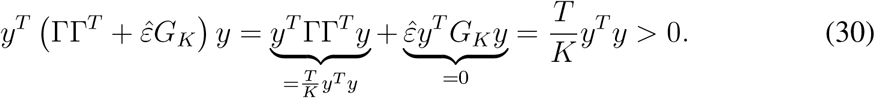 This proves that 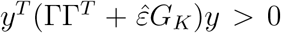 for any *y* ≠ 0, i.e., that the system matrix in (28) is symmetric positive definite and therefore there exists a unique solution of this system given by (26). This also proves that the original objective function of a problem (SPA_2_) with regularization (25) with respect to *S* is for any fixed 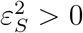 strictly convex and the optimization problem with bounded closed convex feasible set (16) has a unique minimizer. Since for any symmetric matrix and any non-zero *y* it holds *y^T^Ay* ≥ λ_min_(*A*)*y^T^y*, we can combine (29) and (30) to prove the lower estimation in (27). To prove upper estimation, one can use the property of norm and eigenvalues of complete graph Laplace matrix

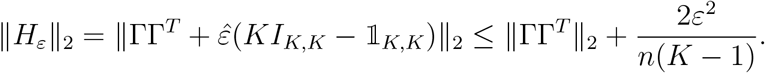

### Lemma 9

(Uniqueness of a reconstruction with the fixed Γ). *Let* [*S*^1*^, Γ^1*^] *and* [*S*^2*^, Γ^2*^] *be two solutions of* (SPA_2_) *for given data X. Let us denote the appropriate reconstructions by X*^rec1^:= *S*^1*^Γ^1*^ *and X*^rec2^:= *S*^2*^Γ^2*^. *If* Γ^1*^ = Γ^2*^ *then X*^rec1^ = *X*^rec2^.

*Proof*. From the optimality conditions, *S*^1*^ and *S*^2*^ solves (SPA_2_) with fixed Γ:= Γ^1*^ = Γ^2*^. All solutions of corresponding QP differ by a vector from kernel of Hessian matrix (see (*7*), (*12*), and (23)) and using Lemma 21 we get

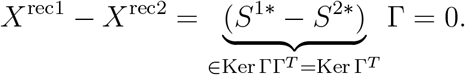

### Lemma 10

(Derivative of solution with a fixed Γ). *Let* Γ ∈ Ω_Γ_ *in problem* (SPA_2_) *with additional regularization function* (25) *be fixed and let S*\*(*X*) *be solution* (26) *for any X. Then for any j* = 1,…,*n and t* = 1,…,*T*

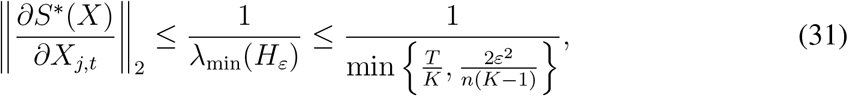

*where* λ_min_(*H_ε_*) *is the smallest eigenvalue of regularized Hessian matrix H_ε_ given by* (26) *and further estimated using* (27).

*Proof*. We use the derivative definition

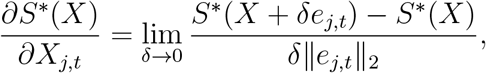

where *e_j,t_* ∈ ℝ^*n,T*^ is a standard basis vector with elements defined by

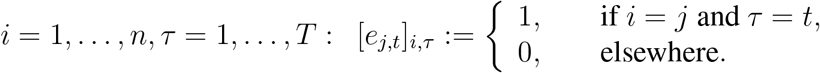

Using the solution (26), the norm can be estimated by

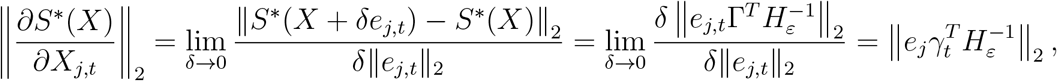

where *e_j_* ∈ ℝ^*n*^ is vector of standard basis and *γ_t_*:= Γ_:,*t*_. Using the property of the norm, we can further estimate

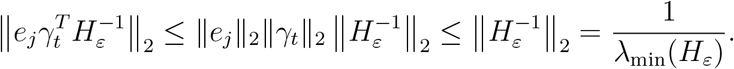

### Corollary 4.

*In the case of K-means, the indicator functions* Γ *are binary and*

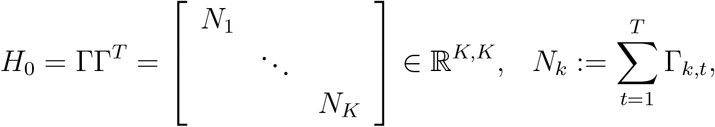

*where N_k_* ≥ 0 *denotes the number of points affiliated to k-th cluster. The eigenvalues of diagonal matrix H*_0_ *are equal to the values on the diagonal, therefore upper estimation* (31) *depends only on the inverse value of the smallest cluster size; it is independent on both of the data size and number of clusters*.

## The solution of Γ subproblem

In this Section, we suppose that in the optimization problem (SPA_2_) the variable *S* is fixed and it remains to solve the problem in a variable Γ only (the second optimization problem of Algorithm 1). In this case, the objective function is additively separable and it can be written in the form of separable Quadratic Programming (QP) problems with linear equality and bound constraints.

### Lemma 11.

*The solution of* (SPA_2_) *with fixed S is equivalent to the solution of T independent QP problems*

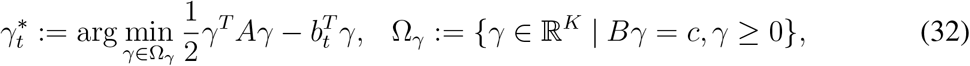

*where*

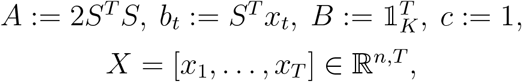

*and the original solution of* (SPA_2_) *can be composed as*

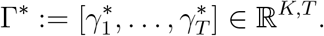

*Proof*. From the definition of Frobenius norm and matrix-matrix multiplication we have

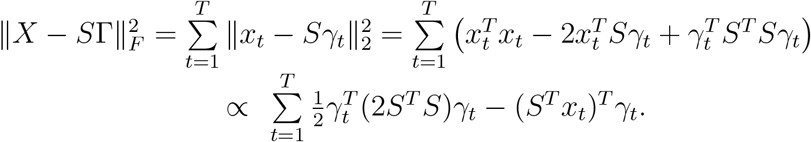

Moreover, it is easy to check that the composition of Ω_*γ*_ for all *γ_t_,t* = 1,…, *T* forms the original feasible set Ω_Γ_. Then using Lemma 4 the problem can be rewritten as the solution of the separated subproblems.

From the computational point of view, the Γ-problem is more challenging since one has to deal with optimization problems on the fasible set described by the combination of linear equality constraints and bound constraints. In the case of QP (32), the subproblems can be solved by the Interior-Point methods or by the Augumented Lagrangian methods combined with Active-set approach (*10*), (*7*). In our implementation we use the fact that the feasible set Ω_*γ*_ is the simplex of size *K*. Since the objective function is continuously differentiable, then one can use Projected Gradient Descent methods, for example Spectral projected gradient method for QP (*2*), (*12*).

### Lemma 12.

*The computational complexity of decreasing the objective function in* Γ *for a fixed A in* (SPA_2_) *is* 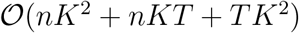, *with a memory complexity of* 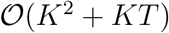.

*Proof*. The complexity of assembling this QP problem is given by the complexity of a matrix-matrix multiplications *S^T^S* and *S^T^X*, which is 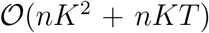. These objects require a memory of the order 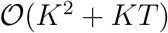.

The number of iterations required for solving this QP problem on convex sets depends on the spectral properties of its Hessian matrix (*7*). Let us focus on one iteration, which will decrease the value of an objective function (32). Such a decrease can be obtained using a projected gradient descend step

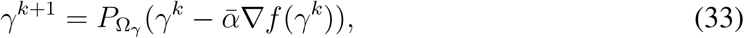

with a step-length 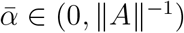. Decrease of the function value for a convex QP on a general closed convex set has been proven in (*6*), (*11*).

The computational complexity of computing the gradient in (33) is 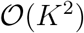 because of the Hessian matrix multiplication. Computational iteration complexity of the projection onto a simplex is of order 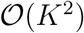 (*4*), (*12*). Since the step has to be performed for all *γ_t_*, the overal complexity is 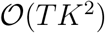. The step for each *γ_t_* requires auxiliary vectors of additional memory 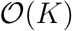, therefore a computation of the whole Γ takes additional 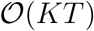 of memory.

### Corollary 5.

*In the case of K-means algorithm, the evaluation of analytical solution* Γ^*^ (9) *consists of evaluation of local error and finding the maxima for all data points. The computational complexity is* 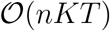 *and the size of auxiliary vectors is* 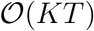.

### Lemma13.

*The computational complexity of one iteration of* (SPA_2_) *is* 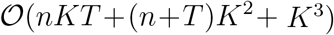, *with a memory complexity of* 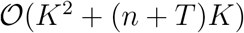.

*Proof*. The Lemma is a direct combination of Lemma 7 and Lemma 12.

### Corollary 6.

*The complexity of one iteration of K-means algorithm can be obtained combining Corollary 3 and Corollary 5. The computational complexity is* 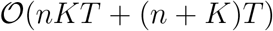 *andthe memory complexity* 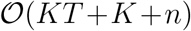. *In practical big data applications the dimension n and the statistics size T are much larger then the discretisation dimension K. It means that in such situations both K-means and SPA will have the same leading order of the computational iteration complexity* 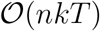 *and the same leading order of the required memory in T, being 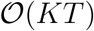. In contrast, spectral clustering methods (like LSD, PCCA+) and density-based clustering methods (like DBSCAN and “mean shift”) will have the leading order in both the computational complexity and in the required memory scaling ranging between* 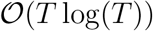 *and* 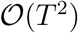.

### Lemma 14.

*Let S* ∈ ℝ^*n,K*^ *be fixed. Function γ*^*^: ℝ^*n*^ → Ω_*γ*_ *defined as*

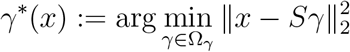

*is a continuous piecewise linear function*.

*Proof*. Let us consider arbitrary *x*_1_, *x*_2_ ∈ ℝ^*n*^ and corresponding *γ*_1_:= *γ*^*^(*x*_1_), *γ*_2_:= *γ*^*^(*x*_2_). Since both of these values solve the optimization problem, there exist appropriate Lagrange multipliers 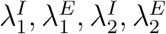 such that the KKT optimality conditions (18), (19), (20), (21), (22) are satisfied in the form

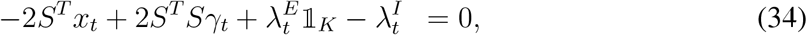

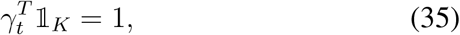

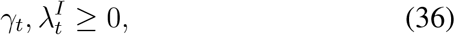

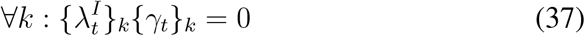

for both of the given *t* ∈ {1, 2}. Let us consider parameter *α* ∈ [0,1], build a convex combination of equations (34) and get

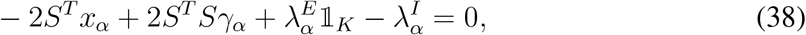

where we denoted

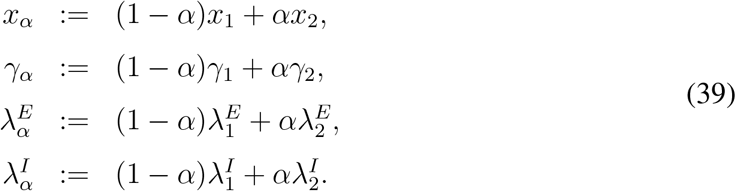

It is easy to see that (38) can be considered as the first KKT optimality condition for any *x_α_* which lies on the line connecting *x*_1_, *x*_2_. In this case, the solution *γ_α_* = *γ*^*^(*x_α_*) of the corresponding optimization problem can be built as a linear combination of *γ*_1_, *γ*_2_ with the same coeficient. The conditions (35) and (36) for *γ_α_* are also satisfied since the feasible set Ω_*γ*_ is convex (and every convex combination of points inside the convex set is also in this set) and/or one can directly check that for any *α* ∈ [0,1]

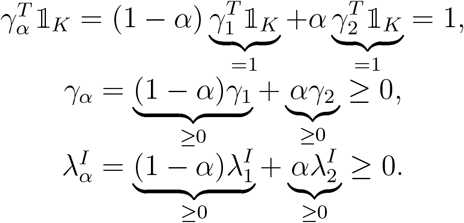

The reason why the function *γ*^*^ is not linear for general *x*_1_, *x*_2_ is the complementarity condition. If we substitute (39) into (37) for α, we obtain

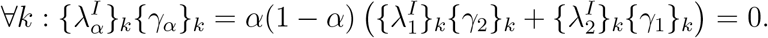

Since (36) and (37) such a condition is satisfied for all *α* ∈ [0,1] if and only if for all *k*

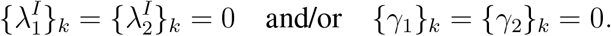

The line connecting *x*_1_, *x*_2_ can be splitted into the segments which satisfied these conditions and therefore the function *γ*^*^ is piecewise linear.

### Corollary 7.

*Let S be fixed and let us define a function*

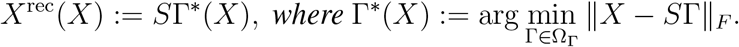

*It is easy to see that this function linearly depends on* Γ^*^(*X*) *and since this separable function is composed from linear functions (see Lemma 14) the derivative*

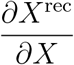

*is a piecewise constant function*.

### Lemma 15.

*Let K* = 2, *S* ∈ ℝ^*n*,2^, *x* ∈ ℝ^*n*^ *be given. Then the optimization problem*

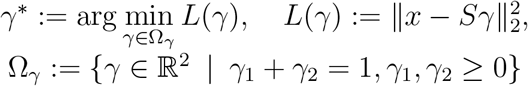

*has a solution*

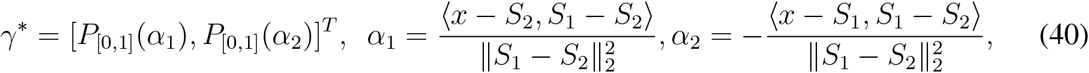

*where* P_[0,1]_(*α*) *is a projection of α* ∈ ℝ *onto interval* [0,1] *given by*

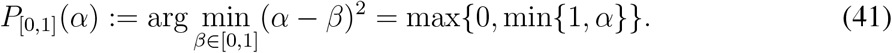

*Proof*. Let us denote the columns of matrix *S* = [*S*_1_, *S*_2_]. The KKT optimality conditions (18), (19), (20), (21), (22) form the system

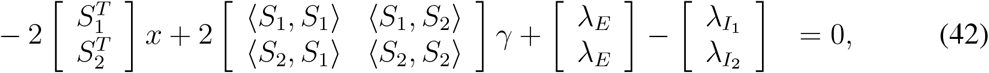

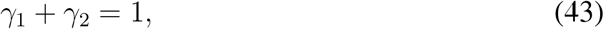

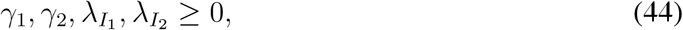

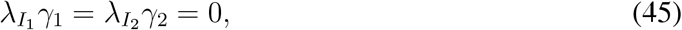

Using the equality (43), we can eliminate variable *γ*_2_ = 1 – *γ*_1_ in (42). Additionally, we can substract the equations and after some manipulations we obtain

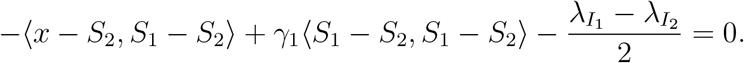

Using the notation (40) for *α*_1_ and including the remaining KKT conditions (44) and (45), we end up with the equivalent system

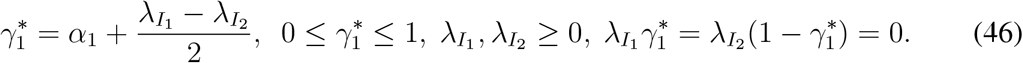

The same system of equations and inequalities can be obtained as KKT system of projection optimization problem (41); here the Lagrange function is given by

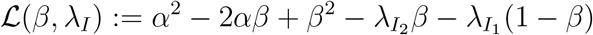

and the KKT optimality conditions can be derived and modified as

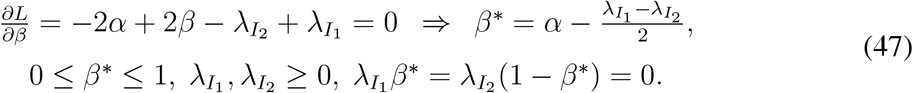

We see that if we denote the output of projection as 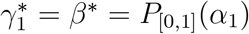 (like in the presented solution (40)) then systems (47) and (46) are the same.

The similar process can be performed to obtain 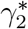, however, in this case, we use *γ*_1_ = 1 – *γ*_2_ to eliminate variable in (42).

### Lemma 16

(Uniqueness of reconstruction with fixed *S*). *Let* [*S*^1*^, Γ^1*^] *and* [*S*^2*^, Γ^2*^] *be two solutions of* (SPA_2_) *for given data X. Let us denote the appropriate reconstructions by X*^rec1^:= *S*^1*^ Γ^1*^ *and X*^rec2^:= *S*^2*^Γ^2*^. *If S*^1*^ = *S*^2*^ *then X*^rec1^ = *X*^rec2^.

*Proof*. From the optimality conditions, Γ^1*^ and Γ^2*^ solves (SPA_2_) with fixed *S*:= *S*^1*^ = *S*^2*^. All solutions of corresponding QP for every *t* = 1,…, *T* differ by a vector from kernel of Hessian matrix (see (*7*), (*12*)) and using Lemma 21 we get

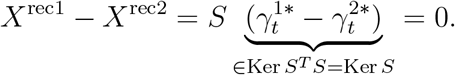

## Computing optimal discretisations for Bayesian and Markovian models

### Theorem 2.

*Let x_t_* ∈ ℝ*^n^ and y_t_* ∈ ℝ^*m*^ *be two time series of length T, X* = [*x*_1_,…, *x_T_*] ∈ ℝ^*n,T*^, *Y* = [*y*_1_,…, *y_T_*] ∈ ℝ^*m,T*^. *The solution of* (SPA_2_) *in the form*

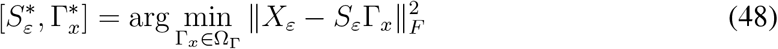

*with*

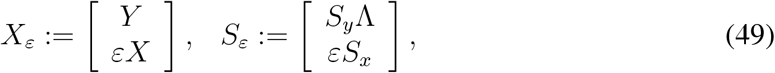

*and ε* ≥ 0 *is equivalent to the solution of* (SPA_2_) *problems*

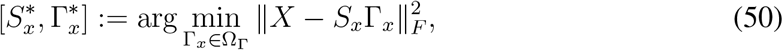

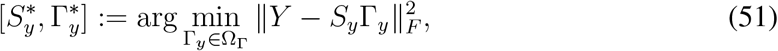

*in Tikhonov-sense with regularization parameter ε and* Λ ∈ ℝ^*K,T*^ *is left-stochastic matrix of conditional probabilities such that the discrete Bayesian and Markovian model equations*

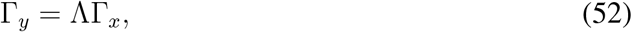

*are satisfied*.

*Proof*. The combination of problems (50) and (51) into one optimization problem using Tikhonov-based approach is given by

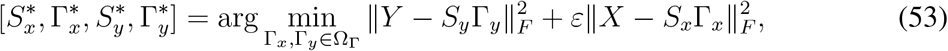

where *ε* ≥ 0 is a Tykhonov-regularisation parameter, controlling the relative importance of the X-discretisation problem with respect to the Y-discretisation problem. Substituting (52) into (53) and using the properties of Frobenius norm, we can write the objective function in form 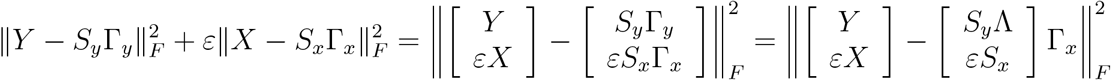.

Getting use of (49) we can reformulate optimization problem (53) into form (48).

## Feature selection with SPA in the Euclidean space

### Lemma17.

*Let S* ∈ ℝ^*n,K*^ *be given. We consider x* ∈ ℝ^*n*^ *and its small perturbation x*+*d* ∈ ℝ^*n*^. *Let us denote* 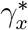 *and* 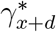 *the optimal probabilistic discretisations of x and x* + *d with respect to S, i.e*.,

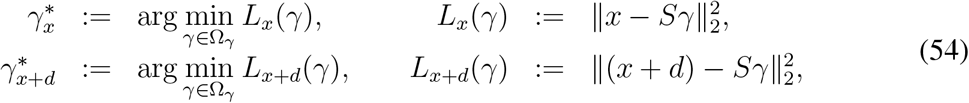

*and* 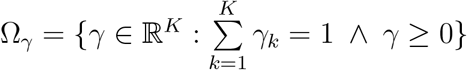 *is a feasible set. Then*

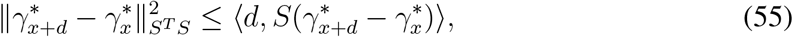

*where* 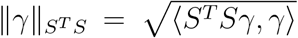 *is a seminorm on* ℝ^*K*^ *induced by the scalar product with a symmetric positive semidefinite matrix S^T^S*.

*Proof*. Using Lemma22 we state that the point *γ*^*^ is a solution of optimization problem if and only if

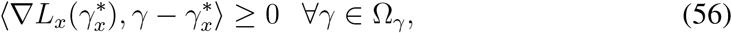

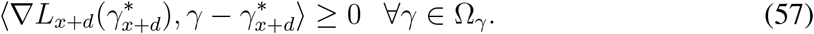

Since the feasible set is the same for both of optimization problems and consequently 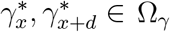, we can choose 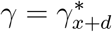 in (56) and 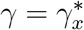 in (57). We get

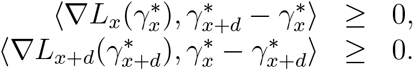

and the sum of these inequalities gives us

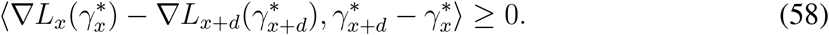

The gradient of the continuously differentiable objective functions can be computed as

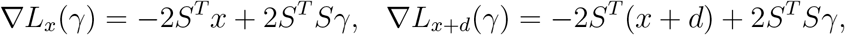

and substituted into (58) to get

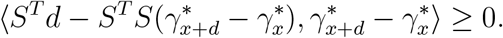

Using the properties of a scalar product, we can rewrite this inequality as (55).

### Corollary 8.

*Let us consider an arbitrary point x* ∈ ℝ^*n*^ *and its perturbation in j-th feature*

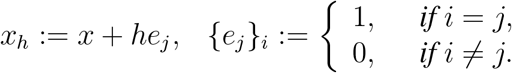

*Let us denote a so-called reconstruction of these points by* 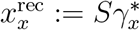 *and* 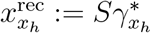. *Since the seminorm on the left-hand side of* (55) *is non-negative, we get using simple subtitution*

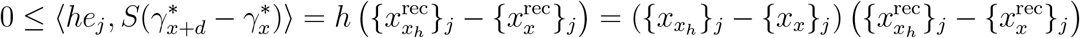

*We can conclude that the sign of the feature change in the data is the same as the sign of the feature change in corresponding reconstructions*.

### Corollary 9.

*Using Cauchy-Bunyakovsky-Schwarz inequality we can further estimate* (55) *to form*

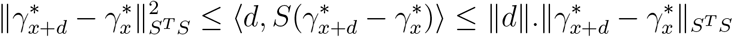

*and therefore*

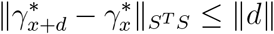

*or using the notation for x*^rec^

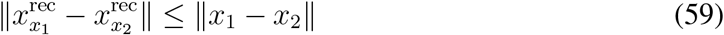

*for any x*_1_, *x*_2_ ∈ ℝ^*n*^.

*The original optimization problem can be rewritten as a projection problem to the set consisting the all possible reconstructed points* Ω_rec_ ⊂ ℝ^*n*^

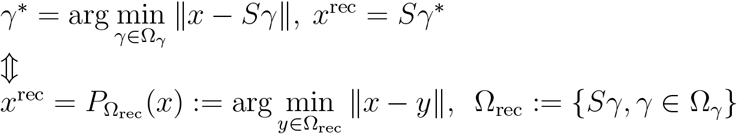

*and the projection is always non-expansive operator, i.e*.,

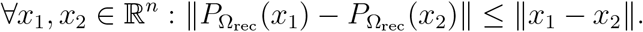

*Additionally, the distance between any* 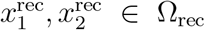 *can be bounded by the largest distance in the feasible set. In the case of the polytope* Ω_rec_, *the largest distance is given by the largest distance between the vertices stored in columns of matrix S, i.e*.,

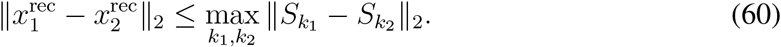

### Theorem 3.

*For sufficiently large T, let* [*S*\*, Γ^*^] *denote the solution of* (SPA_2_) *for X* ∈ ℝ^*n,T*^. *Let X*^rec^(*X*):= *S*^*^(*X*)Γ^*^(*X*) *denotes a reconstruction of the optimal discrete approximation of data X. Then for any dimension j* = 1,…,*n and any t* = 1,…,*T*

1. *if K* = 2 *then*

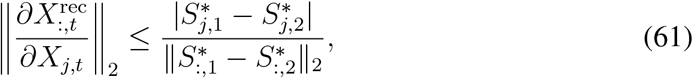
2. if K ≥ 2 *then*

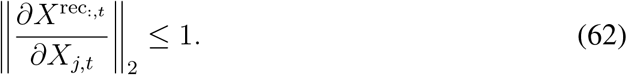

*Proof*. Using the chain rule we get

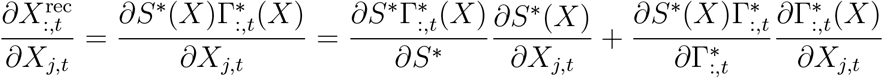

The first term represents the norm of derivate of the recontruction with fixed Γ^*^. We already proved in Lemma 10 that the upper estimation of the norm of this derivative depends on the smallest eigenvalue of matrix ΓΓ^T^. We will suppose that *T* is sufficiently large in a such way that the smallest eigenvalue is sufficiently large and therefore this norm is sufficiently small. In this case, the norm of derivative depends only on second term, i.e., we approximate

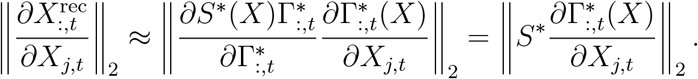

This value represents the norm of derivative of reconstruction with fixed *S*^*^, therefore in the following proof we will suppose that *S*^*^ is fixed.

1. In the case of *K* = 2, we can use an analytical solution of 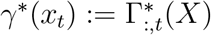 provided by the Lemma 15. Since (for given *S* = [*S*:,1,*S*:,2] ∈ ℝ^*n*,2^ and for any *x_t_* ∈ ℝ^*n*^)

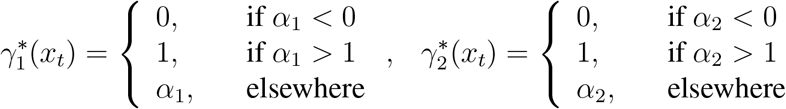

the derivatives are given by

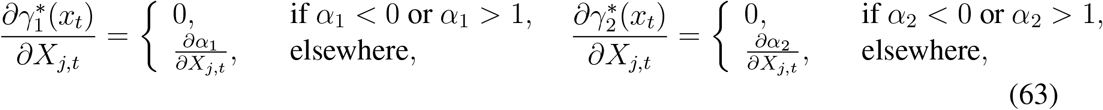

where

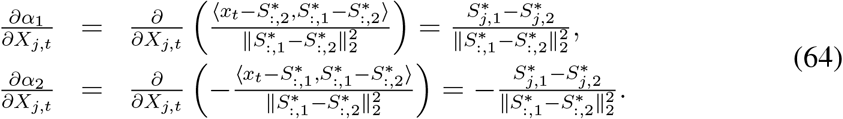 From (63), (64), and since α_1_ + α_2_ = 1 we can easily conclude that

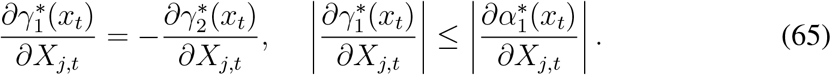 Using the linearity of derivative, the partial derivative of reconstruction 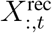 can be computed as

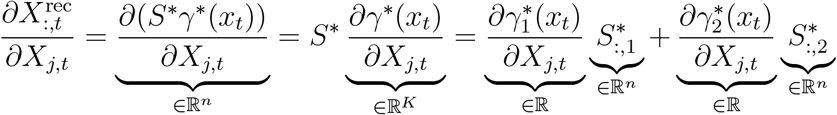

and using (65) we get

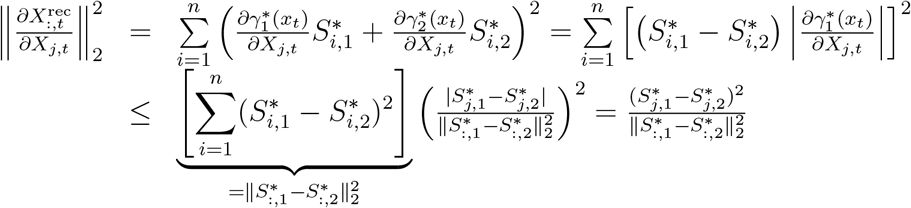
2. From a definition of the derivative we have

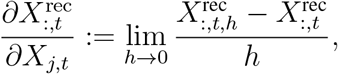

where 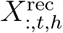 is reconstruction of point *X*_:,*t,h*_ defined as *X*_:,*t*_ with perturbated *j*-th feature, i.e.,

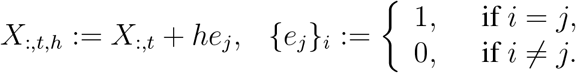 Since the reconstruction 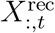 is continuous function of *X*_:,*t*_, we can write

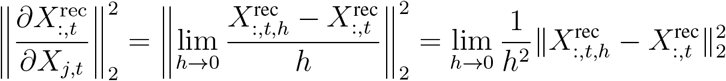 The inner norm can be estimated using (59) to get

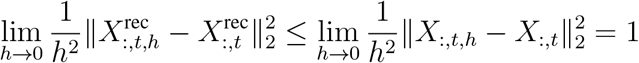

### Corollary 10.

*The previous Lemma motivates for using the regularization of S-problem* (25). *In the case of K* = 2, *such a regularization minimizes the norm of derivative* (61). *In the case of general K, this regularization modifies the resulting polytope generated by S* in a such way that this polytope is distinguishing between the features of reconstructed data, see* (60).

### Corollary 11.

*Please, notice that the dependence of reconstruction of X*^rec^ *on data X is linear and the respective derivative is piecewise constant, see Corollary after Lemma 14. In practice, we can estimate the norm in* (62) *using Euler method. Due to discontinuities in derivatives, such a method is exact for sufficiently small step h*.

---

**Figure S1:**
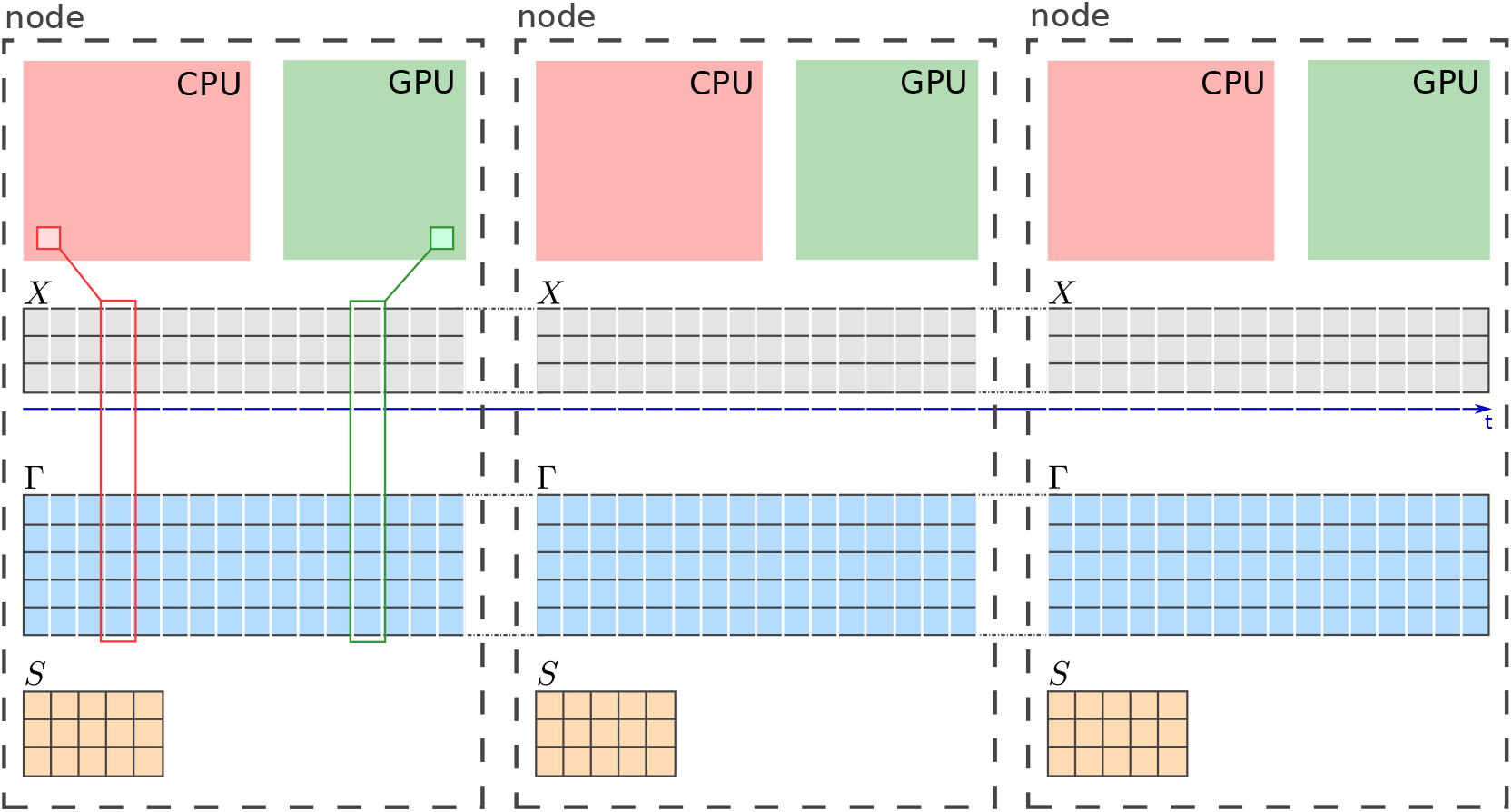
Distributed solution of Γ-problem. If objective function in (SPA), (SPA_2_) is additively separable in *t* then the solution of optimization problem with fixed *S* can be composed as a solution of individual problems (see Lemma 4 and Lemma 11). In such a case, we can distribute *T* independent problems into several computation nodes such that the each node solves its own subset of problems. This local computation can be be performed by local CPU cores and/or using GPU cores, where (again) each core solves its individual subset of local optimization problems. Additionally, if we distribute the data of the problem in the same way, then each computational resource will have an access to its own local part of memory, without any additional communication.

**Figure S2:**
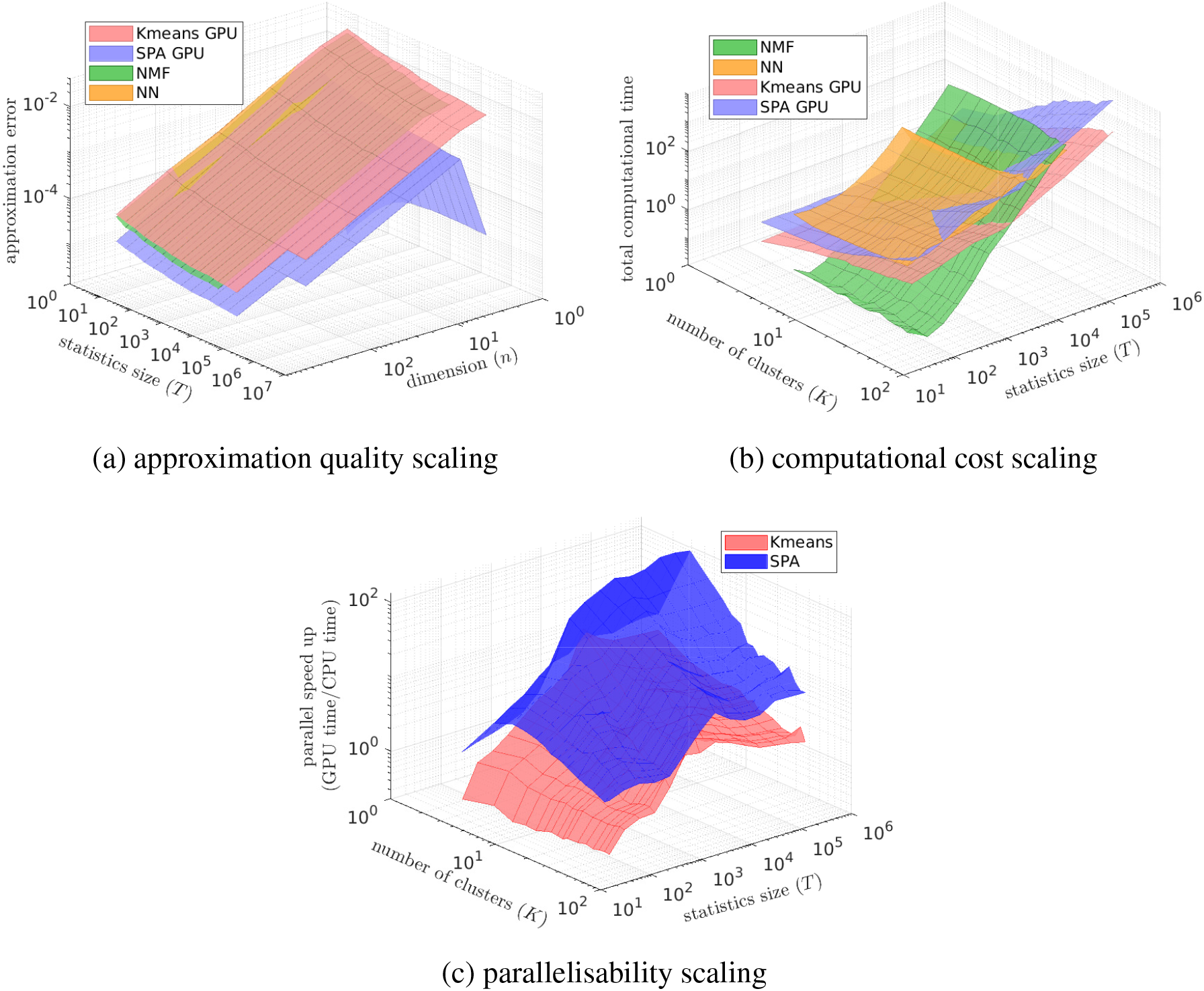
Comparing computational cost (a), discretization quality (b) and parallelizability (c): for (SPA_2_) (blue surfaces), K-means clustering (dark-green), Nonnegative Matrix Factorisation (in its probabilistic variant called Left-Stochastic Decomposition (LSD), magenta surfaces) and the Self-Organising Maps (SOM, a special form of unsupervised neuronal networks used for discretization, orange surfaces). For every combination of data dimension *n* and the data statistics length *T*, methods are applied to 50 same randomly-generated data sets and the results in each of the curves represent averages over these 50 problems. Parallel speed-up in (c) is measured as the ratio of the average times time(GPU)/time(CPU) needed to reach the same relative tolerance threshold of 10^-5^ on a single Graphics Processing Unit (GPU, ASUS TURBO-GTX1080TI-11G, with 3584 CUDA cores) for time(GPU) versus a single CPU core (Intel Core i9-7900X CPU) for time(CPU). MATLAB script Fig1_reproduce.m reproducing these results is available for open access in the repository SPA at https://github.com/SusanneGerber.

**Figure S3:**
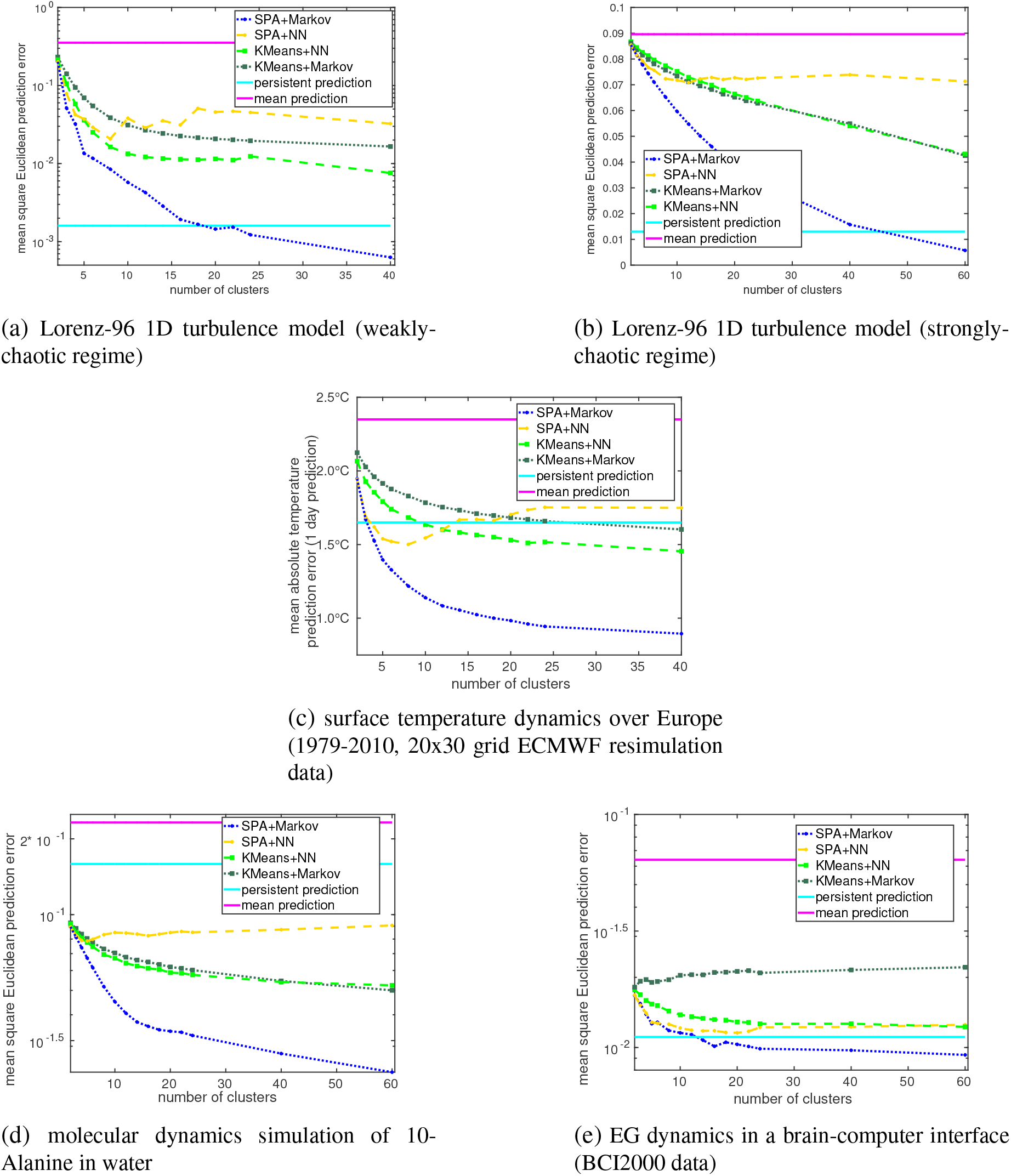
Comparison of one-time-step predictions for a combination of SPA with Markov models (based on applications of the Theorem 2, blue lines) to the one-time-step predictions obtained by the standard prediction methods. The combination of SPA with Markov models is the only prediction scheme that outperforms the persistent prediction (i.e., when the next state is predicted to be the same as the current one) for all of the considered systems.

## APPENDIX

## Definition 1.

*We say that point x^*^ is a minimizer of function f on given feasible set Ω, written as*

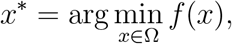

if (and only if) all points from the feasible set have larger or equal function value than f(*x*^*^), *i.e*.,

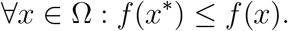

## Lemma 18.

*Let X* ∈ ℝ^*n,T*^, *a, x* ∈ ℝ^*n*^, *b* ∈ ℝ^*n*^, *A* = *A^T^* ∈ ℝ^*n,n*^. *Then*

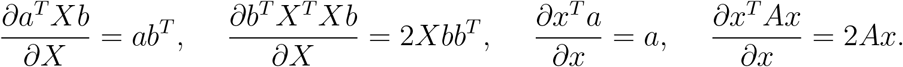

## Lemma 19.

*Let n, K, T* ∈ ℕ *and A* ∈ ℝ^*n,T*^, *B* ∈ ℕ^*K,T*^. *Then*

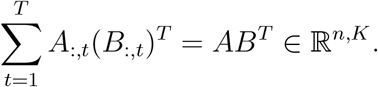

*Proof*. From the definition of matrix-vector multiplication, the components of the result on left-hand side of the equation can be written in form (for every *i* ∈ {1,…,*n*},*j* ∈ {1,…,*K*})

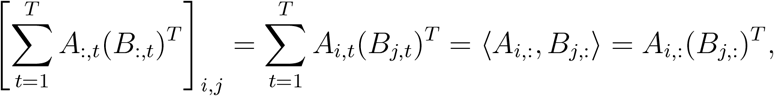

which is a value of the corresponding matrix component on right-hand side of the equation.□

## Lemma 20.

*(of four fundamental subspaces): for any B* ∈ ℝ^*n,m*^ *it holds*^3^

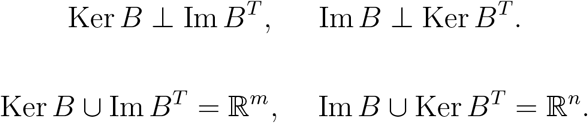

*Proof*. See Laub (*8*).

## Lemma 21.

*Let n, K, T* ∈ ℕ *and A* ∈ ℝ^*n,T*^, *B* ∈ ℝ^*K,T*^. *Then*

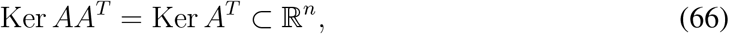

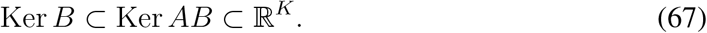

*Proof*. To prove (66), it is necessary to show that

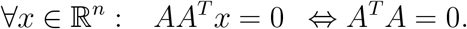

(⇐) Let us consider *x* ∈ ℝ^*m*^ such that *A^T^x* = 0. Then 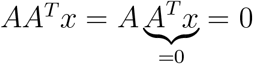 (this also proves (67))

(⇒) Let us consider *x* ∈ ℝ^*m*^ such that *AA^T^x* = 0. Using smart zero, we can write

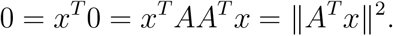

The norm of the vector is equal to zero if and only if the vector is equal to zero, therefore *A^T^x* = 0.

## Lemma 22.

*Let f*: ℝ^*n*^ → ℝ *be a continuously differentiable convex function and let* Ω ⊂ ℝ^*n*^ *be closed convex set. Then x*\* ∈ Ω *is a solution of optimization problem*

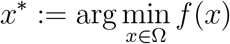

*if and only if*

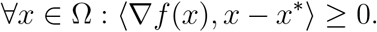

*Proof*. See (*3*), (*10*).

## Footnotes

1 i.e. the matrix such that *AA*^+^*A* = *A, A*^+^*AA*^+^ = *A*^+^, (*AA*^+^)^*T*^ = *AA*^+^, and (*A*^+^*A*)^*T*^ = *A*^+^*A*

2 Since Ker ΓΓ^*T*^ = KerΓ^*T*^ (see (*8*)) we can see that if and only if Γ has linearly independent rows, then matrix ΓΓ^*T*^ is non-singular (invertible).

3 Let 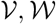 be two subspaces of vector space 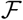 with scalar product 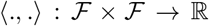. Then we say that 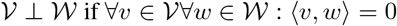. Additionally we define 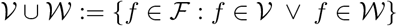.

