## Supplemental Material for "Low-cost scalable discretization, prediction and feature selection for complex systems"

Susanne Gerber,[1,†] Lukáš Pospíšil,[2,†] Mohid Navandar,[1]
Illia Horenko[2,†,*]

[1]Faculty of Biology, Johannes-Gutenberg-University of Mainz,
Anselm-Franz-von-Bentzel-Weg 3, 55128 Mainz, Germany
[2]Faculty of Informatics, Universita della Svizzera Italiana,
Via G. Buffi 13, 6900 Lugano, Switzerland

[†]These authors contributed equally to the paper.


- **Description of the synthetic data problems**
- **General Scalable Probabilistic Approximation (SPA) formulation**
  - Lemma 1 - SPA algorithm generates nonincreasing objective function
  - Lemma 2 - sufficient condition for solvability of $S$ subproblem
  - Lemma 3 - sufficient condition for solvability of $\Gamma$ subproblem
  - Lemma 4 - about separability
  - Theorem 1 - properties of SPA algorithm
  - Corollary 1 - suboptimality of K-means
  - Corollary 2 - suboptimality of FEM-BV and FEM-H1
- **SPA in the Euclidean space**
  - Lemma 5 - non-unique solution of SPA2
  - ○ **Optimality conditions**
  - ○ **The solution of $S$ subproblem**
    - Lemma 6 - analytical solution of $S$-problem
    - Lemma 7 - computational and memory complexity of $S$-problem
    - Corollary 3 - computational and memory complexity of $S$-problem in K-means
    - Lemma 8 - regularization of $S$-problem
    - Lemma 9 - uniqueness of reconstruction with fixed $\Gamma$

Lemma 10 - derivative of solution with fixed $\Gamma$
Corollary 4 - stability of solution in K-means

- **The solution of $\Gamma$ subproblem**
  Lemma 11 - about separability of QP problems
  Lemma 12 - computational and memory complexity of $\Gamma$-problem
  Corollary 5 - computational and memory complexity of $\Gamma$-problem in K-means
  Lemma 13 - complexity of one iteration of ($\text{SPA}_2$)
  Corollary 6 - comparison of leading order complexity scalings for K-means and for ($\text{SPA}_2$)
  Lemma 14 - $\Gamma$ solution is continuous piecewise linear function in $X$
  Corollary 7 - derivative of reconstruction is continuous piecewise constant
  Lemma 15 - analytical solution for $K = 2$
  Lemma 16 - uniqueness of reconstruction with fixed $S$

- **Computing optimal discretisations for Bayesian and Markovian models**
  Theorem 2 - the combination of optimal discretization with Markov model
- **Feature selection with SPA in the Euclidean space**
  Lemma 17 - the estimation of $\Gamma$ subproblem solution stability
  Corollary 8 - the consistency of change of reconstruction and original data
  Corollary 9 - projection onto optimal polytope
  Theorem 3 - $S$ subproblem regularization and feature selection
  Corollary 10 - connection between regularization and feature selection
  Corollary 11 - numerical estimation of reconstruction derivative

### Description of the synthetic data problems (used in the Figure 1 of the main manuscript)

This section provides the description of the benchmark, whose results are presented in Manuscript in the Figure 1. For a given number of data points $T > 0$ and a data dimension (number of features) $n \geq 2$, we generate the random data $X = [x_1, \ldots, x_T] \in \mathbb{R}^{n,T}$ from multivariate normal distribution with different parameters based on a predefined cluster affiliation.

We choose the cluster affiliation in such a way, that the number of points affiliated to cluters

$T_k$ is approximately the same along the clusters, i.e.,

$$\mathcal{T}_k := \left\{ (k-1)\left\lfloor \frac{T}{K} \right\rfloor + 1, \dots, \min\left\{ k\left\lfloor \frac{T}{K} \right\rfloor, T \right\} \right\}$$

denotes the set of point indexes affiliated to $k$-th cluster. Please, notice that these sets are disjoint and union of them forms the set of all point indexes $\{1, \dots, T\}$. Using this decomposition, we generate corresponding data points for every cluster $k = 1, \dots, K$ as random realisations from the multivariate normal distributions

$$\forall t \in \mathcal{T}_k : x_t \sim \mathcal{N}(\mu_k, \Sigma_k),$$

where $\mu_k \in \mathbb{R}^n$ denotes the mean value and $\Sigma_k \in \mathbb{R}^{n,n}$ a covariance matrix.

In our benchmark, we choose $K = 4$ with parameters

$$\mu_1 := 0, \;\; \Sigma_1 = \begin{bmatrix} 0.1 & 0.05 & \\ 0.05 & 0.1 & \\ & & \frac{0.2}{n-2} I_{n-2} \end{bmatrix}, \;\; \mu_2 := \begin{bmatrix} 0.8 \\ 1.6 \\ 0 \\ \vdots \\ 0 \end{bmatrix}, \;\; \Sigma_2 = \begin{bmatrix} 0.1 & -0.05 & \\ -0.05 & 0.1 & \\ & & \frac{0.2}{n-2} I_{n-2} \end{bmatrix},$$

$$\mu_3 := \begin{bmatrix} 1.6 \\ 0 \\ 0 \\ \vdots \\ 0 \end{bmatrix}, \;\; \Sigma_3 = \begin{bmatrix} 1 & 0 & \\ 0 & 1 & \\ & & \frac{0.2}{n-2} I_{n-2} \end{bmatrix}, \;\; \mu_4 := \begin{bmatrix} 0.8 \\ 0.8 \\ 0 \\ \vdots \\ 0 \end{bmatrix}, \;\; \Sigma_3 = \Sigma_4,$$

where $I_{n-2} \in \mathbb{R}^{n-2,n-2}$ is identity matrix.

### General Scalable Probabilistic Approximation (SPA) formulation

The SPA optimization problem is given by

$$[S^*, \Gamma^*] := \arg\min_{\Gamma \in \Omega_\Gamma} L(S, \Gamma), \tag{SPA}$$

where

$$L(S,\Gamma) := \sum_{t=1}^{T} \mathrm{dist}_S(X(t),\Gamma(t)) + \varepsilon_S^2 \Phi_S(S) + \varepsilon_\Gamma^2 \Phi_\Gamma(\Gamma), \quad (1)$$

$$\Omega_\Gamma := \{\Gamma \in \mathbb{R}^{K\times T} \mid \forall k=1,\dots,K : \sum_{k=1}^{K} \Gamma_k(t) = 1, \Gamma_k(t) \geq 0, t=1,\dots,T\}, \quad (2)$$

$T$ denotes the number of data points, $X = \{X(t), t = 1,\dots,T\} \subset \mathcal{X}$ are given data from space $\mathcal{X}$ deployed with the norm $\|\cdot\|$, $K > 1$ denotes the number of discrete states (clusters), $\Gamma = \{\Gamma_k(t), k = 1,\dots,K, t = 1,\dots,T\} \subset \Omega_\Gamma \subset \mathbb{R}^{K\times T}$ are unknown cluster affiliation probability vectors, and $S : \mathbb{R}^K \to \mathcal{X}$ are unknown data representation vectors. We include the possibility of Tikhonov-based regularization of original ill-posed problem using the regularization functions $\Phi_S$, $\Phi_\Gamma$ with corresponding regularization parameters $\varepsilon_S, \varepsilon_\Gamma \geq 0$.

*Set a feasible initial approximation* $\Gamma^0 \in \Omega_\Gamma$

***while*** $\|L(S^k,\Gamma^k) - L(S^{k-1},\Gamma^{k-1})\| \geq \varepsilon$

*solve* $S^k = \arg\min_S L(S,\Gamma^{k-1})$ *(with fixed* $\Gamma^{k-1}$*)*

*solve* $\Gamma^k = \arg\min_{\Gamma\in\Omega_\Gamma} L(S^k,\Gamma)$ *(with fixed* $S^k$*)*

$k = k+1$

***endwhile***

*Return an approximation of the data representation vectors* $S^k$ *and an approximation of cluster affiliation probability vectors* $\Gamma^k$.

Algorithm 1: **General SPA algorithm.**

The problem (SPA) can be solved using the Algorithm 1. The idea is based on the construction of the sequence of split optimization problems. The iteration computational complexity of this algorithm is given by the complexity of the computation of inner optimization problems with fixed variables. The algorithm of this type is well-known as coordinate descent method (*10*) or alternating least-squares method (*1*). The following Lemma presents the basic convergence properties of the algorithm.

**Lemma 1.** *If the solutions of inner optimization problems in Algorithm 1 exist, then algorithm generates a sequence of approximations for the optimization problem* (SPA) *with nonincreasing objective function values, i.e.,*

$$L(S^{k+1}, \Gamma^{k+1}) \leq L(S^k, \Gamma^k) \quad \textit{for } k = 1, 2, \ldots \tag{3}$$

*Proof.* If the solutions of inner optimization problems exist, then the solution process of inner optimization problems provides the approximation with smaller (or the same) function value with respect to non-fixed variable, i.e. (see the Definition 1 in APPENDIX),

$$\forall S : \quad L(S^k, \Gamma^{k-1}) \;\leq\; L(S, \Gamma^{k-1}), \quad \text{in the case of fixed } \Gamma^{k-1}, \tag{4}$$

$$\Gamma \in \Omega_\Gamma : \quad L(S^k, \Gamma^k) \quad \leq \; L(S^k, \Gamma), \qquad \text{in the case of fixed } S^k. \tag{5}$$

Choosing $S = S^{k-1}$ in (4) and $\Gamma = \Gamma^{k-1}$ in (5) we get

$$L(S^{k-1}, \Gamma^{k-1}) \geq L(S^k, \Gamma^{k-1}) \geq L(S^k, \Gamma^k).$$

□

Since the objective function (1) is generally non-convex (but bounded from bellow - each distance function is non-negative), the sequence (3) can possibly converge only to the local optimum. To deal with this non-globality, one has to run the algorithm for several random initial $\Gamma^0$ and choose the solution with the lowest function value. Such a Monte-Carlo-based approach is commonly used for solving the optimization problem with multiple local optimality points and it can be found in literature as annealing steps (*10*).

However, the convergence of the whole process still highly depends on the solvability of inner optimization problems. Following lemmas present the elementary and the most common situations when the solution exists.

**Lemma 2.** *If the distance function* $\mathrm{dist}_S$ *and the regularization function* $\Phi_S$ *in* (SPA) *are convex, bounded from below and continuously differentiable with respect to the variable* $S$*, then the solution with respect to* $S$ *exists and can be found using the necessary optimality conditions for unconstrained problems.*

*Proof.* The Lemma is a consequence of optimization theory fundamental results, see for example (*3*). □

**Lemma 3.** *If the distance function* $\mathrm{dist}_S$ *and the regularization function* $\Phi_\Gamma$ *in* (SPA) *are continuous in variable* $\Gamma$*, then the solution of the problem with respect to* $\Gamma$ *exists.*

*Proof.* Please notice that feasible set $\Omega_\Gamma$ is compact (i.e., closed and bounded) and convex, therefore if $L$ is continuous, then the existence of the solution is a consequence of Weierstrass Extreme Value Theorem (*3*). □

Typically, the largest dimension parameter of the whole problem (SPA) is the number of data points $T$ and the classification data-discretization process (SPA) does not reduce this number. It provides the data representation vectors $S$, whose size is determined by the size of individual data points (the dimension of vector space $\mathcal{X}$) and the number of them is equal to the number of clusters $K$. We can conclude, that the optimization problem with respect to $S$ is much smaller in comparison to the optimization problem with respect to the second variable $\Gamma$. The unknown $\Gamma$ consists of cluster affiliation probability vector of each individual data points, i.e., its size is determined by $T$ and $K$. Fortunately, the objective function $L$ (1) is composed as a sum of local representation errors and therefore if the regularization function $\Phi_\Gamma(\Gamma)$ is also additively separable (the case when it consists of the sum of local regularization functions for individual representations) then the whole minimization problem (SPA) is separable. The following Lemma presents the basic property of additively separable optimization problems.

**Lemma 4.** *If $L$ in the optimization problem (SPA) is additively separable in $t$ (except $\Phi_S(S)$), i.e., there exist functions $L_t(S, \Gamma(t)), t = 1, \dots, T$ such that*

$$L(S, \Gamma) = \left( \sum_{t=1}^{T} L_t(S, \Gamma(t)) \right) + \Phi_S(S), \quad (6)$$

*then the solution of an optimization problem (SPA) with fixed $S$ can be composed from solutions of individual problems*

$$\Gamma^*(t) = \arg \min_{\Gamma \in \Omega_{\Gamma_t}} L_t(S, \Gamma(t)), \quad (7)$$

*where*

$$\Omega_{\Gamma_t} = \{\gamma \in \mathbb{R}^K \ | \ \sum_{k=1}^{K} \gamma_k = 1, \gamma \geq 0\}$$

*and $\Omega_{\Gamma_1} \times \dots \times \Omega_{\Gamma_T} = \Omega_\Gamma$ is the decomposition of the feasible set of the original problem* (SPA).

*Proof.* The definition of optimality point of (7) reads as (see the Definition 1 in APPENDIX)

$$\forall \Gamma(t) \in \Omega_{\Gamma_t} : \ L_t(S, \Gamma^*(t)) \leq L_t(S, \Gamma(t)).$$

Since this inequality can be formulated for all $t = 1, \dots, T$, we can sum these $T$ inequalities to obtain

$$\sum_{t=1}^{T} L_t(S, \Gamma^*(t)) \leq \sum_{t=1}^{T} L_t(S, \Gamma^*(t)).$$

If we add term $\Phi_S(S)$ (constant in $\Gamma$) to both sides of this inequality and use notation (6), we obtain

$$\forall \Gamma \in \Omega_\Gamma : L(S, \Gamma^*) \leq L(S, \Gamma),$$

which is a definition of the optimality point of optimization problem (SPA) with respect to $\Gamma$. □

The separability plays crucial role in the embarassingly parallel computations; one can solve the whole set of $T$ optimization problems independently using modern multi-core architectures, see Figure S2. The $\Gamma$-problem can be splitted into smaller subsets and distributed onto separated computational nodes, which is a commonly adopted approach when working on supercomputers. Each node solves the given subset of problems without any communication with the other nodes. Moreover, if the node includes multi-core processors, then (again) each core can solve independently the part of the node subproblem. This "embarrasingly-parallel" hierarchical computation of the large-scaled problem can be exploited even more when using modern GPU architectures; in this case, the relativelly small $\Gamma(t)$ problem (of size $K$) can be solved using just one computational thread, i.e., one computational core (please see Fig. 2c in the main manuscript).

It is necessary to mention that if the regularization function $\Phi_\Gamma$ is not separable in $T$ (for example when enforcing the persistency of regime/cluster in time, see FEM-H1 and FEM-BV methods (*9*)), then the problem is not embarassingly parallel and computational nodes/cores/threads have to communicate during the solution process. However, as was demonstrated in (*12*), one can still utilize Projected Gradient methods since the projection onto separable simplexes $\Omega_\Gamma$ is still embarassingly parallel.

The following Theorem summarizes the general properties of the Algorithm (1).

**Theorem 1** (Properties of the SPA algorithm)**.** *Let* $X = \{x(t), t = 1, \ldots, T\} \subset \mathcal{X}$ *be given data from space* $\mathcal{X}$*,* $K > 1$ *is a given number of clusters. Let* $\mathrm{dist}_S$*,* $\Phi_S$*,* $\Phi_\Gamma$ *be such functions that* $L(S, \Gamma)$ *in* (SPA) *is convex, bounded from bellow and continuously differentiable with respect to the variable* $S$ *and continuous in the variable* $\Gamma$.

*Then the Algorithm 1*

*(a) is generating a monotonically non-increasing sequence.*

*Moreover, if* $L(S, \Gamma)$ *is additively separable problem in* $\Gamma$*, then the Algorithm 1*

*(b) scales linearly in the size* $T$ *of the data statistics* $X$*,*

*(c) requires the amount of communication independent of the data size.*

*Proof.* (a) is a consequence of Lemma 2, Lemma 3, and Lemma 1. To prove (b) and (c), please notice that the solution of optimization problem with respect to $S$ is independent of the number of provided data points $T$. If the assumption of separability is fullfilled, then in the case of solving the problem with respect to $\Gamma$, we can using Lemma 4 reformulate the original problem as a set of $T$ independent problems, whose dimension is (again) independent of $T$. □

Let us present the connection between SPA and some of the commonly used discretization (clustering) methods in following Corollaries.

**Corollary 1** (Suboptimality of K-means)**.** *Measured in terms of squared Euclidean distance, discretisations providided by K-means are always suboptimal with respect to the discretisations obtained with* (SPA).

*Proof.* Let us consider data $X \in \mathbb{R}^{n,T}$. The aim of the K-means clustering algorithm (**?**) is to optimally partition given data into $K$ disjoint clusters based on the Euclidean distance from (unknown) optimal centroids of the clusters. The algorithm computes these cluster centroids $S_k \in \mathbb{R}^n$ and binary affiliation $\Gamma \in \{0,1\}^{K,T}$, where $\Gamma_{k,t} = 1$ if $x_t$ belongs to $k$-th cluster and $\Gamma_{k,t} = 0$ otherwise. The corresponding optimization problem is formulated as

$$[S^*, \Gamma^*] := \arg \min_{\Gamma \in \Omega_\Gamma} L_{\text{kmeans}}(S, \Gamma), \quad L_{\text{kmeans}}(S, \Gamma) := \sum_{k=1}^{K} \sum_{t=1}^{T} \Gamma_{k,t} \| X(t) - S_k \|_2^2, \tag{8}$$

where $\Omega_\Gamma \subset \{0,1\}^{K,T}$ includes the condition for strict affiliation of a point into exactly one cluster, i.e.,

$$\Omega_\Gamma := \{\Gamma \in \{0,1\}^{K,T} | \forall t = 1, \dots, T : \sum_{k=1}^{K} \Gamma_{k,t} = 1\rangle.$$

The problem (8) is solved iteratively; the feasible initial approximation of affiliations $\Gamma$ is chosen randomly (the points are randomly affiliated to clusters) and afterwards, the iterative procedure solves consecutively the problems with one fixed variable. In the case of K-means, both of the subproblems have analytical solutions

$$S_k^* = \frac{1}{\sum\limits_{t=1}^{T} \Gamma_{k,t}} \sum_{t=1}^{T} \Gamma_{k,t} X(t), \quad \Gamma_{\bar{k},t}^* = \begin{cases} 1 & \text{if } \bar{k} = \arg\min\limits_{k} \|X(t) - S_k\|, \\ 0 & \text{otherwise.} \end{cases} \tag{9}$$

In fact, the scheme of the algorithm is the same as in the Algorithm 1 and one can easily check that if $\Gamma$ is binary variable and we choose $\mathrm{dist}_S(X(t), \Gamma(t)) := \sum\limits_{k=1}^{K} \|X(t) - S\Gamma_k(t)\|_2^2$ in (SPA) (in following text denoted as ($\text{SPA}_2$)) then

$$L(S,\Gamma) = \sum_{t=1}^{T} \sum_{k=1}^{K} \|X(t) - S\Gamma_k(t)\|_2^2 = \sum_{k=1}^{K} \sum_{t=1}^{T} \Gamma_{k,t} \|X(t) - S_k\|_2^2 = L_{\text{kmeans}}(S,\Gamma) \tag{10}$$

and therefore K-means algorithm is equivalent to ($\text{SPA}_2$).

The variant of K-means algorithm with relaxed binary condition is well-known as soft K-means algorithm (**?**). In this case, $\Gamma_{k,t}$ represents the probability that $X(t)$ is affiliated to the $k$-th cluster. The feasible set $\Omega_\Gamma$ enforces the rows of $\Gamma$ to be a corresponding discrete probability density vector, i.e., each element is continuous variable from $[0, 1]$ and because of the law of the total probability, the sum of the elements of this vector has to be equal to one. One can easily check that $\Omega_\Gamma$ defined by (2) represents these conditions. However in the case of continuous $\Gamma$, the equality (10) does not hold. Using the Jensen's inequality (*10*) we get

$$L(S,\Gamma) = \sum_{t=1}^{T} \sum_{k=1}^{K} \|X(t) - S\Gamma_k(t)\|_2^2 \leq \sum_{k=1}^{K} \sum_{t=1}^{T} \Gamma_{k,t} \|X(t) - S_k\|_2^2 = L_{\text{kmeans}}(S,\Gamma)$$

and therefore soft K-means algorithm produces only the upper estimation of the ($\text{SPA}_2$) optimization problem. □

**Corollary 2** (Suboptimality of FEM-BV and FEM-H1.)**.** *Measured in terms of a squared Euclidean distance, discretisations providided by FEM-BV and FEM-H1 methods are always suboptimal with respect to the discretisations obtained with* (SPA).

*Proof.* The family of FEM-BV and FEM-H1 methods consists of methods used for time series analysis (*9*), (*12*). The idea is to extend stationary models with clustering and additional time regularization for enforcing the model time persistency.

In time series modelling, we suppose that the measured data $x_1, x_2, \ldots, x_T \in \mathbb{R}^n$ are described by the parametric model $\psi$ and include the additive noise, i.e.,

$$x_t = \psi(t, \Theta) + \varepsilon_t. \tag{11}$$

For instance one can consider autoregessive models, e.g., the Var-X model defined as

$$\psi(t, \Theta) = \mu + \sum_{i=0}^{p} A_i x_{t-i\tau} + \sum_{j=0}^{q} B_j u_{t-j\tau}, \tag{12}$$

where $\Theta = (\mu, A_0, \ldots, A_p, B_0, \ldots, B_q)$ includes all model parameters, $\tau > 0$ is a discretisation time step, $p, q \geq 0$ represent the size of memory, and $u_t$ denote the external factors or controls. The aim of the analysis is to find parameters of the model which fit the given data $x_t, u_t$ in an optimal way, for example, one can utilize minimum least square error to formulate optimization problem

$$\Theta^* := \arg\min_{\Theta} \sum_{t=1}^{T} \|x_t - \psi(t, \Theta)\|_2^2. \tag{13}$$

In the case of Var-X model (12) the optimization problem (13) is unconstrained quadratic programming problem and the necessary optimality conditions formulate the corresponding system of linear equations which has to be solved.

FEM-BV and FEM-H1 belong to the non-stationary models; here we suppose that the parameters of model $\Theta$ are non-stationary, i.e., they are changing (can change) in time. In general, non-stationary model without any additional assumptions, e.g., restriction of the set of permissible parameters, lead to ill-posed and biased results. In the case of FEM-BV and FEM-H1, we include the assumption of the time persistency of model parameters introducing the finite number of regimes (i.e., clusters) in which the model parameters are stationary. The switching between those regimes is realized by a hidden regime-switching process, which describes the activity of each regime in a given time. For example, if we consider stationary Var-X model (12) on each of the $K$ regimes, then the corresponding optimization problem is formulated as

$$[\Theta^*, \Gamma^*] := \arg \min_{\Theta, \Gamma \in \Omega_\Gamma} \sum_{t=1}^{T} \sum_{k=1}^{K} \Gamma_{k,t} \|x_t - \psi(t, \Theta_k)\|_2^2 + \varepsilon^2 \Phi_\Gamma(\Gamma), \tag{14}$$

where $\Theta = [\Theta_1, \ldots, \Theta_K]$ includes (unknown) parameters of local models on regimes and $\Gamma_{k,:}$ are model indicator functions defined in similar as in the case of K-means, i.e., $\Gamma_{k,t} = 1$ if the time series in time $t$ is in $k$-th regime and and $\Gamma_{k,t} = 0$ otherwise. Regularization function $\Phi_\Gamma(\Gamma)$ with regularization parameter $\varepsilon^2 \geq 0$ enforces the time persistency of a regime-switching process. In the case of FEM-BV, we consider Bounded variation (BV) norm defined as

$$\Phi_\Gamma(\Gamma) := \sum_{k=1}^{K} \sum_{t=1}^{T-1} |\Gamma_{k,t+1} - \Gamma_{k,t}|.$$

If we consider binary $\Gamma$ then this value is equal to the number of switches between regimes and the regularization by this function decreases the global number of switches in the solution. The optimization problem (14) is solved using Algorithm 1, however, in this case the $\Gamma$ subproblem is not separable due to non-separable regularization term and this problem of dimension $KT$ has to be solved using linear programming algorithm. For extended details on the method see (*9*).

It is straightforward to verify that the formulation of FEM-BV corresponds to (SPA) with distance function defined as a local Euclidean distance between given data $X(t)$ and the local value of model $\psi$

$$\text{dist}_\Theta(X(t), \Gamma(t)) := \|X(t) - \psi(t, \Theta_\Gamma(t))\|^2, \quad \Theta_\Gamma(t) = \sum_{k=1}^{K} \Gamma_{k,t} \Theta_k. \tag{15}$$

Similarly to the soft K-means clustering case considered in the Corollary 1 above, we can relax the hard clustering property (i.e., the property that each data point is exclusively affiliated to exactly one regime) considering $\Gamma_{k,t}$ to be probability of affiliation of $X(t)$ to $k$-th regime.

Each $\Gamma_{:,t}$ forms the discrete probability density vector of affiliation of $X(t)$ to regimes and a corresponding feasible set is given by (2). To include the assumption of time persistency, one can adopt the H1 half-norm

$$\Phi_\Gamma(\Gamma) := \sum_{k=1}^{K} \sum_{t=1}^{T-1} (\Gamma_{k,t+1} - \Gamma_{k,t})^2$$

to get the FEM-H1 method, see (*9*). The problem is solved by an Algorithm 1, the corresponding $\Gamma$ subproblem is non-separable convex quadratic programming problem of size $KT$, see (*12*).

Please notice that $\Theta$ depends linearly on variable $\Gamma$, the Var-X model depends linearly on parameters $\Theta$, and the distance function $\mathrm{dist}_\Theta$ is convex in variable $\psi$. Summarizing these properties we can state that distance function is convex in $\Gamma$ (see (*3*) for the list of operations which preserve convexity). Using the Jensen's inequality we get

$$L(S,\Gamma) = \sum_{t=1}^{T} \sum_{k=1}^{K} \|X(t) - \psi(t, \Theta_\Gamma(t))\|_2^2 \leq \sum_{t=1}^{T} \sum_{k=1}^{K} \Gamma_{k,t} \|X(t) - \psi(t, \Theta_k)\|_2^2 = L_{\mathrm{FEM}}(S,\Gamma).$$

This inequality holds also when we add any regularization $\Phi_\Gamma(\Gamma)$ to the both sides. Hence, FEM-BV and FEM-H1 algorithms produce only the upper estimation of the (SPA) optimization problem with a corresponding choice of distance function and regularization. □

### SPA in the Euclidean space

We consider the data from real $n$-dimensional vector space $\mathcal{X} := \mathbb{R}^n$ and Euclidean distance measure on $\mathcal{X}$ defined by

$$\mathrm{dist}_S(X(t), \Gamma(t)) := \sum_{k=1}^{K} \|X(t) - S\Gamma_k(t)\|_2^2.$$

For simplicity, we compose the vectors into matrices

$$X := [X(1), \ldots, X(T)] \in \mathbb{R}^{n,T},\ \Gamma := [\Gamma(t), \ldots, \Gamma(T)] \in \mathbb{R}^{K,T},\ S \in \mathbb{R}^{n,K}$$

and afterwards, the corresponding optimization problem (SPA) without regularization can be written in a form

$$[S^*, \Gamma^*] := \arg \min_{\Gamma \in \Omega_\Gamma} \|X - S\Gamma\|_F^2, \tag{SPA$_2$}$$

where $F$ denotes Frobenius norm and the feasible set is defined by

$$\Omega_\Gamma := \{\Gamma \in \mathbb{R}^{K,T} \mid \forall t = 1, \dots, T \; \forall k = 1, \dots, K : \sum_{k=1}^{K} \Gamma_{k,t} = 1, \Gamma_{k,t} \geq 0\}. \tag{16}$$

**Lemma 5.** *The solutions of problem* (SPA$_2$) *are always non-unique for any* $K > 1$.

*Proof.* Let us consider an arbitrary solution $[S^*, \Gamma^*]$ and nonsingular matrix $R \in \mathbb{R}^{K,K}, R \neq I_{K,K}$ such that $R\Gamma \in \Omega_\Gamma$. Such a matrix always exists, e.g., we can consider a permutation matrix which permutes the rows of $\Gamma$, i.e., the indexes of clusters. Since we can write

$$L(S^*, \Gamma^*) = \|X - S^*\Gamma^*\|_F^2 = \|X - S^* \underbrace{R^{-1}R}_{=I} \Gamma^*\|_F^2 = L(S^*R^{-1}, R\Gamma^*),$$

we can state that feasible $[S^*R^{-1}, R\Gamma^*] \neq [S^*, \Gamma^*]$ has the same (minimal) function value and therefore it also solves the problem. □

#### Optimality conditions

We define the Lagrange function (*10*) corresponding to the optimization problem (SPA$_2$) by

$$\mathcal{L}(S, \Gamma, \lambda^E, \lambda^I) := \|X - S\Gamma\|_F^2 + \sum_{t=1}^{T} \lambda_t^E \left( \sum_{k=1}^{K} \Gamma_{k,t} - 1 \right) - \sum_{t=1}^{T} \sum_{k=1}^{K} \lambda_{k,t}^I \Gamma_{k,t}.$$

Here $\lambda^E \in \mathbb{R}^T$ are Lagrange multipliers corresponding to equality constraints defined by the feasible set (16) and $\lambda^I \in \mathbb{R}^{K,T}$ denotes the Lagrange multipliers corresponding to the non-negativity bound constraints in (16).

The full system of Karush-Kuhn-Tucker (KKT) optimality conditions for this system will be:

$$\nabla_S \mathcal{L}(S,\Gamma,\lambda^E,\lambda^I) = \; -2X\Gamma^T + 2S\Gamma\Gamma^T = 0, \tag{17}$$

$$\nabla_\Gamma \mathcal{L}(S,\Gamma,\lambda^E,\lambda^I) = \; -2S^TX + 2S^TS\Gamma + (\lambda^E)^T \otimes \mathbb{1}_K - \lambda^I = 0, \tag{18}$$

$$\nabla_{\lambda_E} \mathcal{L}(S,\Gamma,\lambda^E,\lambda^I) = \; \Gamma^T\mathbb{1}_K - \mathbb{1}_T = 0, \tag{19}$$

$$\nabla_{\lambda_I} \mathcal{L}(S,\Gamma,\lambda^E,\lambda^I) = \; -\Gamma \leq 0, \tag{20}$$

$$\lambda^I \geq 0, \tag{21}$$

$$\forall k,t: \; \lambda^I_{k,t}\Gamma_{k,t} = 0, \tag{22}$$

where $\mathbb{1}_K \in \mathbb{R}^K, \mathbb{1}_T \in \mathbb{R}^T$ denotes the vectors of ones. Equations (17) and (18) are first-order optimality conditions, equation (19) and inequality (20) are constraints given by the definition of the feasible set (16), inequality (21) preserves the non-negativity of inequality Lagrange multipliers, and equations (22) represent the so-called complementarity conditions for inequality constraints.

#### The solution of $S$ subproblem

**Lemma 6** (The solution of $S$-problem)**.** *Let $\Gamma \in \Omega_\Gamma$ in problem* (SPA$_2$) *be fixed. Then the system of all solutions of optimization problem* (SPA$_2$) *with respect to $S$ is given by*

$$S^* = X\Gamma^T\left(\Gamma\Gamma^T\right)^+ + \alpha^TR^T, \;\; \textit{with parameter } \alpha \in \mathbb{R}^{r,n}, \tag{23}$$

*where $\left(\Gamma\Gamma^T\right)^+ \in \mathbb{R}^{K,K}$ denotes a pseudoinverse*[1] *of the matrix $\Gamma\Gamma^T$, $R \in \mathbb{R}^{K,r}$ is a matrix whose columns form the basis of the null space of $\Gamma^T$, i.e.*

$$\operatorname{Im} R = \operatorname{Ker} \Gamma^T, \tag{24}$$

*and $r = \dim \operatorname{Ker} \Gamma^T$ denotes the nullity of matrix $\Gamma^T$.*

[1] i.e. the matrix such that $AA^+A = A$, $A^+AA^+ = A^+$, $(AA^+)^T = AA^+$, and $(A^+A)^T = A^+A$

*Proof.* Please notice that the objective function of (SPA$_2$) in terms of variable $S$ is continuously differentiable convex matrix quadratic function. The necessary optimality condition of given unconstrained optimization problem is given by (17). This system of linear equations with multiple right-hand side vectors with symmetric positive semi-definite system matrix always has a solution. If the system matrix is non-singular, then the unique solution is given by

$$S^* = X\Gamma^T(\Gamma\Gamma^T)^{-1}.$$

However, the non-singularity of system matrix $\Gamma\Gamma^T \in \mathbb{R}^{K,K}$ (and consequently, the existence of inverse matrix) is not guaranteed[2], the system of all solutions is given by (23) where all solutions differ by the vector from $\operatorname{Ker}\Gamma\Gamma^T$, see (*8*) or (*7*). □

Next we deal with the eventual ill-posedness of the optimization problem (SPA$_2$) in variable $S$, or equivalently, with the ill-posedness of the system of linear equations (17). Deploying Tykhonov-regularization, we reformulate the original (SPA) problem choosing the regularization function

$$\Phi_S(S) := \frac{1}{nK(K-1)} \sum_{i=1}^{n} \sum_{k_1=1}^{K} \sum_{k_2=1}^{K} \left(S_{i,k_1} - S_{i,k_2}\right)^2 \tag{25}$$

and consider regularization parameter $\varepsilon_S^2 > 0$. Please notice that the solution of the optimization problem in term of variable $S$ is independent on the choice of regularization function $\Phi_\Gamma$. The following Lemma proves that (25) guarantees the unique solvability of $S$-problem.

**Lemma 7.** *The computational complexity of solving $S$ subproblem in* (SPA$_2$) *is $\mathcal{O}(K^3 + KnT)$, with the memory complexity of $\mathcal{O}(K^2 + nK)$.*

*Proof.* The first step in solving the $S$ subproblem is the assembly of the matrix $\Gamma\Gamma^T$ and of the matrix of the right-hand side vectors $X\Gamma^T$ in an equation (17). Let us remind that the complexity

[2] Since $\operatorname{Ker}\Gamma\Gamma^T = \operatorname{Ker}\Gamma^T$ (see (*8*)) we can see that if and only if $\Gamma$ has linearly independent rows, then matrix $\Gamma\Gamma^T$ is non-singular (invertible).

of computing matrix-matrix multiplication of general (non-sparse) matrices $A \in \mathbb{R}^{n,m}$ and $B \in \mathbb{R}^{m,p}$ is $\mathcal{O}(nmp)$, therefore in our case, the overall complexity of assembling the problem is $\mathcal{O}(TK^2)+\mathcal{O}(nTK)$. The memory required to store these two new matrices is $\mathcal{O}(K^2)+\mathcal{O}(nK)$.

In general, the direct methods for solving a system of linear equation $Ax = b, A \in \mathbb{R}^{m,m}$ have the complexity of order $\mathcal{O}(m^3)$. Iterative methods, like Krylov subspace algorithms, are based on the iterations where the computational complexity scaling in the leading order is dominated by the multiplication with a system matrix $A$, which is of order $\mathcal{O}(m^2)$. Number of iterations needed for the convergence, when using a suitable preconditioner, is usually much less than $\mathcal{O}(n)$. Therefore, the overall work for solving the system of linear equations is less than $\mathcal{O}(m^3)$. In general, numerical linear algebra algorithms for this purpose are using the auxiliary vectors of dimension $\mathbb{R}^m$, whose number is independent on the dimension of the problem. Therefore, the amount of additional memory used for solving the system of linear equations is of the order $\mathcal{O}(m)$.

Applying these general results to $S$ subproblem which consists of $T$ linear systems of dimension $K$, we obtain the total computational complexity $\mathcal{O}(TK^3)$ and a memory complexity $\mathcal{O}(TK)$. Since the system matrix is the same for all subsystems, therefore one can compute pseudoinverse and use (23) directly, which will lead to the total computational complexity of $\mathcal{O}(n^3) + \mathcal{O}(K^2T)$. In practial applications the computation of pseudoinverse is typically much slower than solving the system of linear equations.

□

**Corollary 3.** *In the case of K-means algorithm, evaluation of an analytical solution $S^*$ (9) consists of computing two sums with the computational complexity $\mathcal{O}((n+K)T)$. To compute the sums, one has to use an additional auxiliary vector of dimension $\mathcal{O}(K)$.*

**Lemma 8** ($S$-problem with regularization)**.** *Let $\Gamma \in \Omega_\Gamma$ in a problem* ($\text{SPA}_2$) *with an additional regularization function* (25) *be fixed. Then, for any $\varepsilon_S^2 > 0$ the problem with respect to $S$ has a* *unique solution given by*

$$S^* = X\Gamma^T H_\epsilon^{-1}, H_\varepsilon := \Gamma\Gamma^T + \frac{2\varepsilon^2}{nK(K-1)}(KI_{K,K} - \mathbb{1}_{K,K}), \tag{26}$$

*where* $I_{K,K} \in \mathbb{R}^{K,K}$ *is an identity matrix and* $\mathbb{1}_{K,K} \in \mathbb{R}^{K,K}$ *is a matrix full of ones. Moreover the spectrum of regularized Hessian matrix* $H_\varepsilon$ *can be estimated by*

$$\begin{array}{rcl} \lambda_{\min}(H_\varepsilon) & \geq & \min\left\{\frac{T}{K}, \frac{2\varepsilon^2}{n(K-1)}\right\}, \\ \lambda_{\max}(H_\varepsilon) & \leq & \|\Gamma\Gamma^T\|_2 + \frac{2\varepsilon^2}{n(K-1)}. \end{array} \tag{27}$$

*Proof.* The gradient of the original objective function $L$ in ($\text{SPA}_2$) without regularization is given by the left-hand side of (17). Let us focus on the gradient of regularization function whose components are given by (for every $i \in \{1,\dots,n\}, k \in \{1,\dots,K\}$)

$$\begin{array}{rcl} [\nabla\Phi_S(S)]_{i,k} & = & \frac{1}{nK(K-1)}\left(\sum\limits_{k_2=1}^{K} 2(S_{i,k} - S_{i,k_2}) - \sum\limits_{k_1=1}^{K} 2(S_{i,k_1} - S_{i,k})\right) \\ & = & \frac{2}{nK(K-1)}\left(2KS_{i,k} - 2\sum\limits_{k_1=1}^{K} S_{i,k_1}\right) = \frac{4}{nK(K-1)}\left(KS_{i,k} - S_{i,:}\mathbb{1}_K\right) \end{array}$$

where $\mathbb{1}_K \in \mathbb{R}^K$ is a column vector of ones. It is easy to see that the whole gradient can be written as

$$\nabla\Phi_S(S) = \frac{4}{nK(K-1)}\left(KS - S\mathbb{1}_{K,K}\right)$$

and therefore the necessary optimality condition of the regularized problem is given by the solution of a regularized linear system of equations

$$-2X\Gamma^T + 2S\left(\Gamma\Gamma^T + \frac{2\varepsilon_S^2}{nK(K-1)}(KI_{K,K} - \mathbb{1}_{K,K})\right) = 0. \tag{28}$$

It remains to show that the system matrix is non-singular for any $\varepsilon_S^2 > 0$ and therefore we will be able to multiply the whole equation with the matrix inverse to obtain a unique solution.

Please notice that the matrix $G_K := KI_{K,K} - \mathbb{1}_{K,K}$ is a Laplacian matrix of a complete graph on $K$ nodes, therefore it is symmetric positive semidefinite and $\operatorname{Ker} G_K = \operatorname{span}\{\mathbb{1}_K\}$, see (5).

For the simplicity, let us denote $\hat{\varepsilon} := \frac{2\varepsilon_S^2}{nK(K-1)} > 0$. For any non-zero $y \in \mathbb{R}^K$ we can differentiate two cases

- if $y \notin \operatorname{Ker} G_K$ then $y^T G_K y = K y^T y$ (the spectrum of complete graph Laplace matrix is composed from one zero eigenvalue and eigenvalues of value $K$ with multiplicity $K-1$, see (5)) and

$$y^T \left(\Gamma\Gamma^T + \hat{\varepsilon} G_K\right) y = \underbrace{y^T \Gamma\Gamma^T y}_{\geq 0} + \hat{\varepsilon} \underbrace{y^T G_K y}_{= K y^T y} \geq \hat{\varepsilon} K y^T y > 0. \tag{29}$$

- if $y \in \operatorname{Ker} G_K = \operatorname{span}\{\mathbb{1}_K\}$ then there exists a non-zero $\alpha \in \mathbb{R}$ such that non-zero $y$ can be written as $y = \alpha \mathbb{1}_K$. Using the equality constraints of the feasible set $\Omega_\Gamma$ (16) written in a form $\Gamma^T \mathbb{1}_K = \mathbb{1}_T$ we can state that

$$y^T \Gamma\Gamma^T y = \alpha^2 \mathbb{1}_K^T \Gamma\Gamma^T \mathbb{1}_K = \alpha^2 \mathbb{1}_T^T \mathbb{1}_T = \alpha^2 T = \frac{T}{K} \alpha^2 \mathbb{1}_K^T \mathbb{1}_K = \frac{T}{K} y^T y > 0$$

  and consequently

$$y^T \left(\Gamma\Gamma^T + \hat{\varepsilon} G_K\right) y = \underbrace{y^T \Gamma\Gamma^T y}_{= \frac{T}{K} y^T y} + \underbrace{\hat{\varepsilon} y^T G_K y}_{= 0} = \frac{T}{K} y^T y > 0. \tag{30}$$

This proves that $y^T(\Gamma\Gamma^T + \hat{\varepsilon} G_K) y > 0$ for any $y \neq 0$, i.e., that the system matrix in (28) is symmetric positive definite and therefore there exists a unique solution of this system given by (26). This also proves that the original objective function of a problem ($\text{SPA}_2$) with regularization (25) with respect to $S$ is for any fixed $\varepsilon_S^2 > 0$ strictly convex and the optimization problem with bounded closed convex feasible set (16) has a unique minimizer. Since for any symmetric matrix and any non-zero $y$ it holds $y^T A y \geq \lambda_{\min}(A) y^T y$, we can combine (29) and (30) to prove the lower estimation in (27). To prove upper estimation, one can use the property of norm and eigenvalues of complete graph Laplace matrix

$$\|H_\varepsilon\|_2 = \|\Gamma\Gamma^T + \hat{\varepsilon}(K I_{K,K} - \mathbb{1}_{K,K})\|_2 \leq \|\Gamma\Gamma^T\|_2 + \frac{2\varepsilon^2}{n(K-1)}.$$

□

**Lemma 9** (Uniqueness of a reconstruction with the fixed $\Gamma$)**.** *Let $[S^{1*}, \Gamma^{1*}]$ and $[S^{2*}, \Gamma^{2*}]$ be two solutions of (SPA$_2$) for given data $X$. Let us denote the appropriate reconstructions by $X^{\text{rec1}} := S^{1*}\Gamma^{1*}$ and $X^{\text{rec2}} := S^{2*}\Gamma^{2*}$. If $\Gamma^{1*} = \Gamma^{2*}$ then $X^{\text{rec1}} = X^{\text{rec2}}$.*

*Proof.* From the optimality conditions, $S^{1*}$ and $S^{2*}$ solves (SPA$_2$) with fixed $\Gamma := \Gamma^{1*} = \Gamma^{2*}$. All solutions of corresponding QP differ by a vector from kernel of Hessian matrix (see (*7*), (*12*), and (23)) and using Lemma 21 we get

$$X^{\text{rec1}} - X^{\text{rec2}} = \underbrace{(S^{1*} - S^{2*})}_{\in \text{Ker}\, \Gamma\Gamma^T = \text{Ker}\, \Gamma^T} \Gamma = 0.$$

□

**Lemma 10** (Derivative of solution with a fixed $\Gamma$)**.** *Let $\Gamma \in \Omega_\Gamma$ in problem (SPA$_2$) with additional regularization function (25) be fixed and let $S^*(X)$ be solution (26) for any $X$. Then for any $j = 1, \dots, n$ and $t = 1, \dots, T$*

$$\left\| \frac{\partial S^*(X)}{\partial X_{j,t}} \right\|_2 \leq \frac{1}{\lambda_{\min}(H_\varepsilon)} \leq \frac{1}{\min\left\{ \frac{T}{K}, \frac{2\varepsilon^2}{n(K-1)} \right\}}, \tag{31}$$

*where $\lambda_{\min}(H_\varepsilon)$ is the smallest eigenvalue of regularized Hessian matrix $H_\varepsilon$ given by (26) and further estimated using (27).*

*Proof.* We use the derivative definition

$$\frac{\partial S^*(X)}{\partial X_{j,t}} = \lim_{\delta \to 0} \frac{S^*(X + \delta e_{j,t}) - S^*(X)}{\delta \|e_{j,t}\|_2},$$

where $e_{j,t} \in \mathbb{R}^{n,T}$ is a standard basis vector with elements defined by

$$i = 1, \dots, n, \tau = 1, \dots, T: \quad [e_{j,t}]_{i,\tau} := \begin{cases} 1, & \text{if } i = j \text{ and } \tau = t, \\ 0, & \text{elsewhere.} \end{cases}$$

Using the solution (26), the norm can be estimated by

$$\left\| \frac{\partial S^*(X)}{\partial X_{j,t}} \right\|_2 = \lim_{\delta \to 0} \frac{\|S^*(X + \delta e_{j,t}) - S^*(X)\|_2}{\delta \|e_{j,t}\|_2} = \lim_{\delta \to 0} \frac{\delta \left\| e_{j,t} \Gamma^T H_\varepsilon^{-1} \right\|_2}{\delta \|e_{j,t}\|_2} = \left\| e_j \gamma_t^T H_\varepsilon^{-1} \right\|_2,$$

where $e_j \in \mathbb{R}^n$ is vector of standard basis and $\gamma_t := \Gamma_{:,t}$. Using the property of the norm, we can further estimate

$$\left\|e_j \gamma_t^T H_\varepsilon^{-1}\right\|_2 \leq \|e_j\|_2 \|\gamma_t\|_2 \left\|H_\varepsilon^{-1}\right\|_2 \leq \left\|H_\varepsilon^{-1}\right\|_2 = \frac{1}{\lambda_{\min}(H_\varepsilon)}.$$

□

**Corollary 4.** *In the case of K-means, the indicator functions $\Gamma$ are binary and*

$$H_0 = \Gamma\Gamma^T = \begin{bmatrix} N_1 & & \\ & \ddots & \\ & & N_K \end{bmatrix} \in \mathbb{R}^{K,K}, \quad N_k := \sum_{t=1}^{T} \Gamma_{k,t},$$

*where $N_k \geq 0$ denotes the number of points affiliated to $k$-th cluster. The eigenvalues of diagonal matrix $H_0$ are equal to the values on the diagonal, therefore upper estimation (31) depends only on the inverse value of the smallest cluster size; it is independent on both of the data size and number of clusters.*

#### The solution of $\Gamma$ subproblem

In this Section, we suppose that in the optimization problem ($\text{SPA}_2$) the variable $S$ is fixed and it remains to solve the problem in a variable $\Gamma$ only (the second optimization problem of Algorithm 1). In this case, the objective function is additively separable and it can be written in the form of separable Quadratic Programming (QP) problems with linear equality and bound constraints.

**Lemma 11.** *The solution of ($\text{SPA}_2$) with fixed $S$ is equivalent to the solution of $T$ independent QP problems*

$$\gamma_t^* := \arg\min_{\gamma \in \Omega_\gamma} \frac{1}{2}\gamma^T A \gamma - b_t^T \gamma, \quad \Omega_\gamma := \{\gamma \in \mathbb{R}^K \mid B\gamma = c, \gamma \geq 0\}, \tag{32}$$

*where*

$$A := 2S^T S,\ b_t := S^T x_t,\ B := \mathbb{1}_K^T,\ c := 1,$$
$$X = [x_1, \ldots, x_T] \in \mathbb{R}^{n,T},$$

*and the original solution of* (SPA$_2$) *can be composed as*

$$\Gamma^* := [\gamma_1^*, \dots, \gamma_T^*] \in \mathbb{R}^{K,T}.$$

*Proof.* From the definition of Frobenius norm and matrix-matrix multiplication we have

$$\|X - S\Gamma\|_F^2 = \sum_{t=1}^{T} \|x_t - S\gamma_t\|_2^2 = \sum_{t=1}^{T} \left( x_t^T x_t - 2x_t^T S\gamma_t + \gamma_t^T S^T S\gamma_t \right)$$
$$\propto \sum_{t=1}^{T} \tfrac{1}{2}\gamma_t^T (2S^T S)\gamma_t - (S^T x_t)^T \gamma_t.$$

Moreover, it is easy to check that the composition of $\Omega_\gamma$ for all $\gamma_t, t = 1, \dots, T$ forms the original feasible set $\Omega_\Gamma$. Then using Lemma 4 the problem can be rewritten as the solution of the separated subproblems. □

From the computational point of view, the $\Gamma$-problem is more challenging since one has to deal with optimization problems on the fasible set described by the combination of linear equality constraints and bound constraints. In the case of QP (32), the subproblems can be solved by the Interior-Point methods or by the Augumented Lagrangian methods combined with Active-set approach (*10*), (*7*). In our implementation we use the fact that the feasible set $\Omega_\gamma$ is the simplex of size $K$. Since the objective function is continuously differentiable, then one can use Projected Gradient Descent methods, for example Spectral projected gradient method for QP (*2*), (*12*).

**Lemma 12.** *The computational complexity of decreasing the objective function in* $\Gamma$ *for a fixed* $A$ *in* (SPA$_2$) *is* $\mathcal{O}(nK^2 + nKT + TK^2)$*, with a memory complexity of* $\mathcal{O}(K^2 + KT)$*.*

*Proof.* The complexity of assembling this QP problem is given by the complexity of a matrix-matrix multiplications $S^T S$ and $S^T X$, which is $\mathcal{O}(nK^2 + nKT)$. These objects require a memory of the order $\mathcal{O}(K^2 + KT)$.

The number of iterations required for solving this QP problem on convex sets depends on the spectral properties of its Hessian matrix (*7*). Let us focus on one iteration, which will decrease the value of an objective function (32). Such a decrease can be obtained using a projected gradient descend step

$$\gamma^{k+1} = P_{\Omega_\gamma}(\gamma^k - \bar{\alpha}\nabla f(\gamma^k)), \tag{33}$$

with a step-length $\bar{\alpha} \in (0, \|A\|^{-1})$. Decrease of the function value for a convex QP on a general closed convex set has been proven in (*6*), (*11*).

The computational complexity of computing the gradient in (33) is $\mathcal{O}(K^2)$ because of the Hessian matrix multiplication. Computational iteration complexity of the projection onto a simplex is of order $\mathcal{O}(K^2)$ (*4*), (*12*). Since the step has to be performed for all $\gamma_t$, the overal complexity is $\mathcal{O}(TK^2)$. The step for each $\gamma_t$ requires auxiliary vectors of additional memory $\mathcal{O}(K)$, therefore a computation of the whole $\Gamma$ takes additional $\mathcal{O}(KT)$ of memory. □

**Corollary 5.** *In the case of K-means algorithm, the evaluation of analytical solution* $\Gamma^*$ (9) *consists of evaluation of local error and finding the maxima for all data points. The computational complexity is* $\mathcal{O}(nKT)$ *and the size of auxiliary vectors is* $\mathcal{O}(KT)$.

**Lemma 13.** *The computational complexity of one iteration of* ($\text{SPA}_2$) *is* $\mathcal{O}(nKT+(n+T)K^2+K^3)$*, with a memory complexity of* $\mathcal{O}(K^2+(n+T)K)$.

*Proof.* The Lemma is a direct combination of Lemma 7 and Lemma 12. □

**Corollary 6.** *The complexity of one iteration of K-means algorithm can be obtained combining Corollary 3 and Corollary 5. The computational complexity is* $\mathcal{O}(nKT+(n+K)T)$ *and the memory complexity* $\mathcal{O}(KT+K+n)$*. In practical big data applications the dimension* $n$ *and the statistics size* $T$ *are much larger then the discretisation dimension* $K$*. It means that in such situations both K-means and SPA will have the same leading order of the computational iteration complexity* $\mathcal{O}(nkT)$ *and the same leading order of the required memory in* $T$*, being* $\mathcal{O}(KT)$*. In contrast, spectral clustering methods (like LSD, PCCA+) and density-based clustering meth-*

*ods (like DBSCAN and "mean shift") will have the leading order in both the computational complexity and in the required memory scaling ranging between $\mathcal{O}(T\log(T))$ and $\mathcal{O}(T^2)$.*

**Lemma 14.** *Let $S \in \mathbb{R}^{n,K}$ be fixed. Function $\gamma^* : \mathbb{R}^n \to \Omega_\gamma$ defined as*

$$\gamma^*(x) := \arg\min_{\gamma \in \Omega_\gamma} \|x - S\gamma\|_2^2$$

*is a continuous piecewise linear function.*

*Proof.* Let us consider arbitrary $x_1, x_2 \in \mathbb{R}^n$ and corresponding $\gamma_1 := \gamma^*(x_1), \gamma_2 := \gamma^*(x_2)$. Since both of these values solve the optimization problem, there exist appropriate Lagrange multipliers $\lambda_1^I, \lambda_1^E, \lambda_2^I, \lambda_2^E$ such that the KKT optimality conditions (18), (19), (20), (21), (22) are satisfied in the form

$$-2S^T x_t + 2S^T S\gamma_t + \lambda_t^E \mathbb{1}_K - \lambda_t^I = 0, \tag{34}$$

$$\gamma_t^T \mathbb{1}_K = 1, \tag{35}$$

$$\gamma_t, \lambda_t^I \geq 0, \tag{36}$$

$$\forall k : \{\lambda_t^I\}_k \{\gamma_t\}_k = 0 \tag{37}$$

for both of the given $t \in \{1, 2\}$. Let us consider parameter $\alpha \in [0, 1]$, build a convex combination of equations (34) and get

$$-2S^T x_\alpha + 2S^T S\gamma_\alpha + \lambda_\alpha^E \mathbb{1}_K - \lambda_\alpha^I = 0, \tag{38}$$

where we denoted

$$\begin{aligned} x_\alpha &:= (1-\alpha)x_1 + \alpha x_2, \\ \gamma_\alpha &:= (1-\alpha)\gamma_1 + \alpha\gamma_2, \\ \lambda_\alpha^E &:= (1-\alpha)\lambda_1^E + \alpha\lambda_2^E, \\ \lambda_\alpha^I &:= (1-\alpha)\lambda_1^I + \alpha\lambda_2^I. \end{aligned} \tag{39}$$

It is easy to see that (38) can be considered as the first KKT optimality condition for any $x_\alpha$ which lies on the line connecting $x_1, x_2$. In this case, the solution $\gamma_\alpha = \gamma^*(x_\alpha)$ of the corresponding optimization problem can be built as a linear combination of $\gamma_1, \gamma_2$ with the same coeficient. The conditions (35) and (36) for $\gamma_\alpha$ are also satisified since the feasible set $\Omega_\gamma$ is convex (and every convex combination of points inside the convex set is also in this set) and/or one can directly check that for any $\alpha \in [0, 1]$

$$\gamma_\alpha^T \mathbb{1}_K = (1-\alpha)\underbrace{\gamma_1^T \mathbb{1}_K}_{=1} + \alpha \underbrace{\gamma_2^T \mathbb{1}_K}_{=1} = 1,$$
$$\gamma_\alpha = \underbrace{(1-\alpha)\gamma_1}_{\geq 0} + \underbrace{\alpha\gamma_2}_{\geq 0} \geq 0,$$
$$\lambda_\alpha^I = \underbrace{(1-\alpha)\lambda_1^I}_{\geq 0} + \underbrace{\alpha\lambda_2^I}_{\geq 0} \geq 0.$$

The reason why the function $\gamma^*$ is not linear for general $x_1, x_2$ is the complementarity condition. If we substitute (39) into (37) for $\alpha$, we obtain

$$\forall k : \{\lambda_\alpha^I\}_k \{\gamma_\alpha\}_k = \alpha(1-\alpha)\left(\{\lambda_1^I\}_k\{\gamma_2\}_k + \{\lambda_2^I\}_k\{\gamma_1\}_k\right) = 0.$$

Since (36) and (37) such a condition is satisfied for all $\alpha \in [0, 1]$ if and only if for all $k$

$$\{\lambda_1^I\}_k = \{\lambda_2^I\}_k = 0 \quad \text{and/or} \quad \{\gamma_1\}_k = \{\gamma_2\}_k = 0.$$

The line connecting $x_1, x_2$ can be splitted into the segments which satisfied these conditions and therefore the function $\gamma^*$ is piecewise linear. □

**Corollary 7.** *Let $S$ be fixed and let us define a function*

$$X^{\mathrm{rec}}(X) := S\Gamma^*(X), \text{ where } \Gamma^*(X) := \arg\min_{\Gamma \in \Omega_\Gamma} \|X - S\Gamma\|_F.$$

*It is easy to see that this function linearly depends on $\Gamma^*(X)$ and since this separable function is composed from linear functions (see Lemma 14) the derivative*

$$\frac{\partial X^{\mathrm{rec}}}{\partial X}$$

*is a piecewise constant function.*

**Lemma 15.** *Let $K = 2$, $S \in \mathbb{R}^{n,2}, x \in \mathbb{R}^n$ be given. Then the optimization problem*

$$\gamma^* := \arg\min_{\gamma \in \Omega_\gamma} L(\gamma), \quad L(\gamma) := \|x - S\gamma\|_2^2,$$
$$\Omega_\gamma := \{\gamma \in \mathbb{R}^2 \mid \gamma_1 + \gamma_2 = 1, \gamma_1, \gamma_2 \geq 0\}$$

*has a solution*

$$\gamma^* = [P_{[0,1]}(\alpha_1), P_{[0,1]}(\alpha_2)]^T, \quad \alpha_1 = \frac{\langle x - S_2, S_1 - S_2 \rangle}{\|S_1 - S_2\|_2^2}, \alpha_2 = -\frac{\langle x - S_1, S_1 - S_2 \rangle}{\|S_1 - S_2\|_2^2}, \tag{40}$$

*where $P_{[0,1]}(\alpha)$ is a projection of $\alpha \in \mathbb{R}$ onto interval $[0,1]$ given by*

$$P_{[0,1]}(\alpha) := \arg\min_{\beta \in [0,1]} (\alpha - \beta)^2 = \max\{0, \min\{1, \alpha\}\}. \tag{41}$$

*Proof.* Let us denote the columns of matrix $S = [S_1, S_2]$. The KKT optimality conditions (18), (19), (20), (21), (22) form the system

$$-2\begin{bmatrix} S_1^T \\ S_2^T \end{bmatrix} x + 2\begin{bmatrix} \langle S_1, S_1 \rangle & \langle S_1, S_2 \rangle \\ \langle S_2, S_1 \rangle & \langle S_2, S_2 \rangle \end{bmatrix} \gamma + \begin{bmatrix} \lambda_E \\ \lambda_E \end{bmatrix} - \begin{bmatrix} \lambda_{I_1} \\ \lambda_{I_2} \end{bmatrix} = 0, \tag{42}$$

$$\gamma_1 + \gamma_2 = 1, \tag{43}$$

$$\gamma_1, \gamma_2, \lambda_{I_1}, \lambda_{I_2} \geq 0, \tag{44}$$

$$\lambda_{I_1}\gamma_1 = \lambda_{I_2}\gamma_2 = 0, \tag{45}$$

Using the equality (43), we can eliminate variable $\gamma_2 = 1 - \gamma_1$ in (42). Additionally, we can substract the equations and after some manipulations we obtain

$$-\langle x - S_2, S_1 - S_2 \rangle + \gamma_1 \langle S_1 - S_2, S_1 - S_2 \rangle - \frac{\lambda_{I_1} - \lambda_{I_2}}{2} = 0.$$

Using the notation (40) for $\alpha_1$ and including the remaining KKT conditions (44) and (45), we end up with the equivalent system

$$\gamma_1^* = \alpha_1 + \frac{\lambda_{I_1} - \lambda_{I_2}}{2}, \;\; 0 \leq \gamma_1^* \leq 1, \;\; \lambda_{I_1}, \lambda_{I_2} \geq 0, \;\; \lambda_{I_1}\gamma_1^* = \lambda_{I_2}(1 - \gamma_1^*) = 0. \tag{46}$$

The same system of equations and inequalities can be obtained as KKT system of projection optimization problem (41); here the Lagrange function is given by

$$\mathcal{L}(\beta, \lambda_I) := \alpha^2 - 2\alpha\beta + \beta^2 - \lambda_{I_2}\beta - \lambda_{I_1}(1-\beta)$$

and the KKT optimality conditions can be derived and modified as

$$\begin{array}{c} \frac{\partial L}{\partial \beta} = -2\alpha + 2\beta - \lambda_{I_2} + \lambda_{I_1} = 0 \;\; \Rightarrow \;\; \beta^* = \alpha - \frac{\lambda_{I_1} - \lambda_{I_2}}{2}, \\ 0 \leq \beta^* \leq 1, \; \lambda_{I_1}, \lambda_{I_2} \geq 0, \; \lambda_{I_1}\beta^* = \lambda_{I_2}(1-\beta^*) = 0. \end{array} \tag{47}$$

We see that if we denote the output of projection as $\gamma_1^* = \beta^* = P_{[0,1]}(\alpha_1)$ (like in the presented solution (40)) then systems (47) and (46) are the same.

The similar process can be performed to obtain $\gamma_2^*$, however, in this case, we use $\gamma_1 = 1 - \gamma_2$ to eliminate variable in (42). ☐

**Lemma 16** (Uniqueness of reconstruction with fixed $S$)**.** *Let* $[S^{1*}, \Gamma^{1*}]$ *and* $[S^{2*}, \Gamma^{2*}]$ *be two solutions of* (SPA$_2$) *for given data* $X$*. Let us denote the appropriate reconstructions by* $X^{\text{rec}1} := S^{1*}\Gamma^{1*}$ *and* $X^{\text{rec}2} := S^{2*}\Gamma^{2*}$*. If* $S^{1*} = S^{2*}$ *then* $X^{\text{rec}1} = X^{\text{rec}2}$*.*

*Proof.* From the optimality conditions, $\Gamma^{1*}$ and $\Gamma^{2*}$ solves (SPA$_2$) with fixed $S := S^{1*} = S^{2*}$. All solutions of corresponding QP for every $t = 1, \dots, T$ differ by a vector from kernel of Hessian matrix (see (*7*), (*12*)) and using Lemma 21 we get

$$X^{\text{rec}1} - X^{\text{rec}2} = S \underbrace{(\gamma_t^{1*} - \gamma_t^{2*})}_{\in \text{Ker}\, S^T S = \text{Ker}\, S} = 0.$$

☐

### Computing optimal discretisations for Bayesian and Markovian models

**Theorem 2.** *Let $x_t \in \mathbb{R}^n$ and $y_t \in \mathbb{R}^m$ be two time series of length $T$, $X = [x_1, \dots, x_T] \in \mathbb{R}^{n,T}, Y = [y_1, \dots, y_T] \in \mathbb{R}^{m,T}$. The solution of* (SPA$_2$) *in the form*

$$[S_\varepsilon^*, \Gamma_x^*] = \arg \min_{\Gamma_x \in \Omega_\Gamma} \|X_\varepsilon - S_\varepsilon \Gamma_x\|_F^2 \tag{48}$$

*with*

$$X_\varepsilon := \begin{bmatrix} Y \\ \varepsilon X \end{bmatrix}, \quad S_\varepsilon := \begin{bmatrix} S_y \Lambda \\ \varepsilon S_x \end{bmatrix}, \tag{49}$$

*and $\varepsilon \geq 0$ is equivalent to the solution of* (SPA$_2$) *problems*

$$[S_x^*, \Gamma_x^*] := \arg \min_{\Gamma_x \in \Omega_\Gamma} \|X - S_x \Gamma_x\|_F^2, \tag{50}$$

$$[S_y^*, \Gamma_y^*] := \arg \min_{\Gamma_y \in \Omega_\Gamma} \|Y - S_y \Gamma_y\|_F^2, \tag{51}$$

*in Tikhonov-sense with regularization parameter $\varepsilon$ and $\Lambda \in \mathbb{R}^{K,T}$ is left-stochastic matrix of conditional probabilities such that the discrete Bayesian and Markovian model equations*

$$\Gamma_y = \Lambda \Gamma_x, \tag{52}$$

*are satisfied.*

*Proof.* The combination of problems (50) and (51) into one optimization problem using Tikhonov-based approach is given by

$$[S_x^*, \Gamma_x^*, S_y^*, \Gamma_y^*] = \arg \min_{\Gamma_x, \Gamma_y \in \Omega_\Gamma} \|Y - S_y \Gamma_y\|_F^2 + \varepsilon \|X - S_x \Gamma_x\|_F^2, \tag{53}$$

where $\varepsilon \geq 0$ is a Tykhonov-regularisation parameter, controlling the relative importance of the X-discretisation problem with respect to the Y-discretisation problem. Substituting (52) into (53) and using the properties of Frobenius norm, we can write the objective function in form

$$\|Y - S_y \Gamma_y\|_F^2 + \varepsilon \|X - S_x \Gamma_x\|_F^2 = \left\| \begin{bmatrix} Y \\ \varepsilon X \end{bmatrix} - \begin{bmatrix} S_y \Gamma_y \\ \varepsilon S_x \Gamma_x \end{bmatrix} \right\|_F^2 = \left\| \begin{bmatrix} Y \\ \varepsilon X \end{bmatrix} - \begin{bmatrix} S_y \Lambda \\ \varepsilon S_x \end{bmatrix} \Gamma_x \right\|_F^2.$$

Getting use of (49) we can reformulate optimization problem (53) into form (48). □

#### Feature selection with SPA in the Euclidean space

**Lemma 17.** *Let $S \in \mathbb{R}^{n,K}$ be given. We consider $x \in \mathbb{R}^n$ and its small perturbation $x+d \in \mathbb{R}^n$. Let us denote $\gamma_x^*$ and $\gamma_{x+d}^*$ the optimal probabilistic discretisations of $x$ and $x+d$ with respect to $S$, i.e.,*

$$\begin{array}{rclrcl} \gamma_x^* & := & \arg\min\limits_{\gamma \in \Omega_\gamma} L_x(\gamma), & L_x(\gamma) & := & \|x - S\gamma\|_2^2, \\ \gamma_{x+d}^* & := & \arg\min\limits_{\gamma \in \Omega_\gamma} L_{x+d}(\gamma), & L_{x+d}(\gamma) & := & \|(x+d) - S\gamma\|_2^2, \end{array} \tag{54}$$

*and $\Omega_\gamma = \{\gamma \in \mathbb{R}^K : \sum\limits_{k=1}^{K} \gamma_k = 1 \ \wedge \ \gamma \geq 0\}$ is a feasible set. Then*

$$\|\gamma_{x+d}^* - \gamma_x^*\|_{S^TS}^2 \leq \langle d, S(\gamma_{x+d}^* - \gamma_x^*)\rangle, \tag{55}$$

*where $\|\gamma\|_{S^TS} = \sqrt{\langle S^TS\gamma, \gamma\rangle}$ is a seminorm on $\mathbb{R}^K$ induced by the scalar product with a symmetric positive semidefinite matrix $S^TS$.*

*Proof.* Using Lemma 22 we state that the point $\gamma^*$ is a solution of optimization problem if and only if

$$\langle \nabla L_x(\gamma_x^*), \gamma - \gamma_x^* \rangle \geq 0 \quad \forall \gamma \in \Omega_\gamma, \tag{56}$$

$$\langle \nabla L_{x+d}(\gamma_{x+d}^*), \gamma - \gamma_{x+d}^* \rangle \geq 0 \quad \forall \gamma \in \Omega_\gamma. \tag{57}$$

Since the feasible set is the same for both of optimization problems and consequently $\gamma_x^*, \gamma_{x+d}^* \in \Omega_\gamma$, we can choose $\gamma = \gamma_{x+d}^*$ in (56) and $\gamma = \gamma_x^*$ in (57). We get

$$\begin{array}{rcl} \langle \nabla L_x(\gamma_x^*), \gamma_{x+d}^* - \gamma_x^* \rangle & \geq & 0, \\ \langle \nabla L_{x+d}(\gamma_{x+d}^*), \gamma_x^* - \gamma_{x+d}^* \rangle & \geq & 0. \end{array}$$

and the sum of these inequalities gives us

$$\langle \nabla L_x(\gamma_x^*) - \nabla L_{x+d}(\gamma_{x+d}^*), \gamma_{x+d}^* - \gamma_x^* \rangle \geq 0. \tag{58}$$

The gradient of the continuously differentiable objective functions can be computed as

$$\nabla L_x(\gamma) = -2S^Tx + 2S^TS\gamma, \quad \nabla L_{x+d}(\gamma) = -2S^T(x+d) + 2S^TS\gamma,$$

and substituted into (58) to get

$$\langle S^T d - S^T S(\gamma^*_{x+d} - \gamma^*_x), \gamma^*_{x+d} - \gamma^*_x \rangle \geq 0.$$

Using the properties of a scalar product, we can rewrite this inequality as (55). □

**Corollary 8.** *Let us consider an arbitrary point $x \in \mathbb{R}^n$ and its perturbation in $j$-th feature*

$$x_h := x + he_j, \quad \{e_j\}_i := \begin{cases} 1, & \text{if } i = j, \\ 0, & \text{if } i \neq j. \end{cases}$$

*Let us denote a so-called reconstruction of these points by $x^{\text{rec}}_x := S\gamma^*_x$ and $x^{\text{rec}}_{x_h} := S\gamma^*_{x_h}$. Since the seminorm on the left-hand side of (55) is non-negative, we get using simple subtitution*

$$0 \leq \langle he_j, S(\gamma^*_{x+d} - \gamma^*_x)\rangle = h\left(\{x^{\text{rec}}_{x_h}\}_j - \{x^{\text{rec}}_x\}_j\right) = (\{x_{x_h}\}_j - \{x_x\}_j)\left(\{x^{\text{rec}}_{x_h}\}_j - \{x^{\text{rec}}_x\}_j\right)$$

*We can conclude that the sign of the feature change in the data is the same as the sign of the feature change in corresponding reconstructions.*

**Corollary 9.** *Using Cauchy-Bunyakovsky-Schwarz inequality we can further estimate (55) to form*

$$\|\gamma^*_{x+d} - \gamma^*_x\|^2_{S^TS} \leq \langle d, S(\gamma^*_{x+d} - \gamma^*_x)\rangle \leq \|d\|.\|\gamma^*_{x+d} - \gamma^*_x\|_{S^TS}$$

*and therefore*

$$\|\gamma^*_{x+d} - \gamma^*_x\|_{S^TS} \leq \|d\|$$

*or using the notation for $x^{\text{rec}}$*

$$\|x^{\text{rec}}_{x_1} - x^{\text{rec}}_{x_2}\| \leq \|x_1 - x_2\| \tag{59}$$

*for any $x_1, x_2 \in \mathbb{R}^n$.*

*The original optimization problem can be rewritten as a projection problem to the set consisting the all possible reconstructed points $\Omega_{\text{rec}} \subset \mathbb{R}^n$*

$$\begin{aligned} &\gamma^* = \arg\min_{\gamma \in \Omega_\gamma} \|x - S\gamma\|, \; x^{\text{rec}} = S\gamma^* \\ &\Updownarrow \\ &x^{\text{rec}} = P_{\Omega_{\text{rec}}}(x) := \arg\min_{y \in \Omega_{\text{rec}}} \|x - y\|, \;\; \Omega_{\text{rec}} := \{S\gamma, \gamma \in \Omega_\gamma\} \end{aligned}$$

*and the projection is always non-expansive operator, i.e.,*

$$\forall x_1, x_2 \in \mathbb{R}^n : \|P_{\Omega_{\text{rec}}}(x_1) - P_{\Omega_{\text{rec}}}(x_2)\| \leq \|x_1 - x_2\|.$$

*Additionally, the distance between any $x_1^{\text{rec}}, x_2^{\text{rec}} \in \Omega_{\text{rec}}$ can be bounded by the largest distance in the feasible set. In the case of the polytope $\Omega_{\text{rec}}$, the largest distance is given by the largest distance between the vertices stored in columns of matrix $S$, i.e.,*

$$\|x_1^{\text{rec}} - x_2^{\text{rec}}\|_2 \leq \max_{k_1,k_2} \|S_{k_1} - S_{k_2}\|_2. \tag{60}$$

**Theorem 3.** *For sufficiently large $T$, let $[S^*, \Gamma^*]$ denote the solution of (SPA$_2$) for $X \in \mathbb{R}^{n,T}$. Let $X^{\text{rec}}(X) := S^*(X)\Gamma^*(X)$ denotes a reconstruction of the optimal discrete approximation of data $X$. Then for any dimension $j = 1, \dots, n$ and any $t = 1, \dots, T$*

*1.) if $K = 2$ then*

$$\left\| \frac{\partial X^{\text{rec}}_{:,t}}{\partial X_{j,t}} \right\|_2 \leq \frac{|S^*_{j,1} - S^*_{j,2}|}{\|S^*_{:,1} - S^*_{:,2}\|_2}, \tag{61}$$

*2.) if $K \geq 2$ then*

$$\left\| \frac{\partial X^{\text{rec}_{:,t}}}{\partial X_{j,t}} \right\|_2 \leq 1. \tag{62}$$

*Proof.* Using the chain rule we get

$$\frac{\partial X^{\text{rec}}_{:,t}}{\partial X_{j,t}} = \frac{\partial S^*(X)\Gamma^*_{:,t}(X)}{\partial X_{j,t}} = \frac{\partial S^*\Gamma^*_{:,t}(X)}{\partial S^*} \frac{\partial S^*(X)}{\partial X_{j,t}} + \frac{\partial S^*(X)\Gamma^*_{:,t}}{\partial \Gamma^*_{:,t}} \frac{\partial \Gamma^*_{:,t}(X)}{\partial X_{j,t}}$$

The first term represents the norm of derivate of the recontruction with fixed $\Gamma^*$. We already proved in Lemma 10 that the upper estimation of the norm of this derivative depends on the smallest eigenvalue of matrix $\Gamma\Gamma^T$. We will suppose that $T$ is sufficiently large in a such way that the smallest eigenvalue is sufficiently large and therefore this norm is sufficiently small. In this case, the norm of derivative depends only on second term, i.e., we approximate

$$\left\| \frac{\partial X^{\text{rec}}_{:,t}}{\partial X_{j,t}} \right\|_2 \approx \left\| \frac{\partial S^*(X)\Gamma^*_{:,t}}{\partial \Gamma^*_{:,t}} \frac{\partial \Gamma^*_{:,t}(X)}{\partial X_{j,t}} \right\|_2 = \left\| S^* \frac{\partial \Gamma^*_{:,t}(X)}{\partial X_{j,t}} \right\|_2.$$

This value represents the norm of derivative of reconstruction with fixed $S^*$, therefore in the following proof we will suppose that $S^*$ is fixed.

1.) In the case of $K = 2$, we can use an analytical solution of $\gamma^*(x_t) := \Gamma^*_{:,t}(X)$ provided by the Lemma 15. Since (for given $S = [S_{:,1}, S_{:,2}] \in \mathbb{R}^{n,2}$ and for any $x_t \in \mathbb{R}^n$)

$$\gamma_1^*(x_t) = \begin{cases} 0, & \text{if } \alpha_1 < 0 \\ 1, & \text{if } \alpha_1 > 1 \\ \alpha_1, & \text{elsewhere} \end{cases}, \quad \gamma_2^*(x_t) = \begin{cases} 0, & \text{if } \alpha_2 < 0 \\ 1, & \text{if } \alpha_2 > 1 \\ \alpha_2, & \text{elsewhere} \end{cases}$$

the derivatives are given by

$$\frac{\partial \gamma_1^*(x_t)}{\partial X_{j,t}} = \begin{cases} 0, & \text{if } \alpha_1 < 0 \text{ or } \alpha_1 > 1, \\ \frac{\partial \alpha_1}{\partial X_{j,t}}, & \text{elsewhere,} \end{cases} \quad \frac{\partial \gamma_2^*(x_t)}{\partial X_{j,t}} = \begin{cases} 0, & \text{if } \alpha_2 < 0 \text{ or } \alpha_2 > 1, \\ \frac{\partial \alpha_2}{\partial X_{j,t}}, & \text{elsewhere,} \end{cases} \tag{63}$$

where

$$\begin{aligned} \frac{\partial \alpha_1}{\partial X_{j,t}} &= \frac{\partial}{\partial X_{j,t}} \left( \frac{\langle x_t - S^*_{:,2}, S^*_{:,1} - S^*_{:,2} \rangle}{\|S^*_{:,1} - S^*_{:,2}\|_2^2} \right) = \frac{S^*_{j,1} - S^*_{j,2}}{\|S^*_{:,1} - S^*_{:,2}\|_2^2}, \\ \frac{\partial \alpha_2}{\partial X_{j,t}} &= \frac{\partial}{\partial X_{j,t}} \left( -\frac{\langle x_t - S^*_{:,1}, S^*_{:,1} - S^*_{:,2} \rangle}{\|S^*_{:,1} - S^*_{:,2}\|_2^2} \right) = -\frac{S^*_{j,1} - S^*_{j,2}}{\|S^*_{:,1} - S^*_{:,2}\|_2^2}. \end{aligned} \tag{64}$$

From (63), (64), and since $\alpha_1 + \alpha_2 = 1$ we can easily conclude that

$$\frac{\partial \gamma_1^*(x_t)}{\partial X_{j,t}} = -\frac{\partial \gamma_2^*(x_t)}{\partial X_{j,t}}, \quad \left| \frac{\partial \gamma_1^*(x_t)}{\partial X_{j,t}} \right| \leq \left| \frac{\partial \alpha_1^*(x_t)}{\partial X_{j,t}} \right|. \tag{65}$$

Using the linearity of derivative, the partial derivative of reconstruction $X^{\text{rec}}_{:,t}$ can be computed as

$$\frac{\partial X^{\text{rec}}_{:,t}}{\partial X_{j,t}} = \underbrace{\frac{\partial (S^* \gamma^*(x_t))}{\partial X_{j,t}}}_{\in \mathbb{R}^n} = S^* \underbrace{\frac{\partial \gamma^*(x_t)}{\partial X_{j,t}}}_{\in \mathbb{R}^K} = \underbrace{\frac{\partial \gamma_1^*(x_t)}{\partial X_{j,t}}}_{\in \mathbb{R}} \underbrace{S^*_{:,1}}_{\in \mathbb{R}^n} + \underbrace{\frac{\partial \gamma_2^*(x_t)}{\partial X_{j,t}}}_{\in \mathbb{R}} \underbrace{S^*_{:,2}}_{\in \mathbb{R}^n}$$

and using (65) we get

$$\begin{aligned} \left\| \frac{\partial X^{\text{rec}}_{:,t}}{\partial X_{j,t}} \right\|_2^2 &= \sum_{i=1}^n \left( \frac{\partial \gamma_1^*(x_t)}{\partial X_{j,t}} S^*_{i,1} + \frac{\partial \gamma_2^*(x_t)}{\partial X_{j,t}} S^*_{i,2} \right)^2 = \sum_{i=1}^n \left[ \left( S^*_{i,1} - S^*_{i,2} \right) \left| \frac{\partial \gamma_1^*(x_t)}{\partial X_{j,t}} \right| \right]^2 \\ &\leq \underbrace{\left[ \sum_{i=1}^n (S^*_{i,1} - S^*_{i,2})^2 \right]}_{= \|S^*_{:,1} - S^*_{:,2}\|_2^2} \left( \frac{|S^*_{j,1} - S^*_{j,2}|}{\|S^*_{:,1} - S^*_{:,2}\|_2^2} \right)^2 = \frac{(S^*_{j,1} - S^*_{j,2})^2}{\|S^*_{:,1} - S^*_{:,2}\|_2^2} \end{aligned}$$

2.) From a definition of the derivative we have

$$\frac{\partial X^{\text{rec}}_{:,t}}{\partial X_{j,t}} := \lim_{h\to 0} \frac{X^{\text{rec}}_{:,t,h} - X^{\text{rec}}_{:,t}}{h},$$

where $X^{\text{rec}}_{:,t,h}$ is reconstruction of point $X_{:,t,h}$ defined as $X_{:,t}$ with perturbated $j$-th feature, i.e.,

$$X_{:,t,h} := X_{:,t} + he_j, \quad \{e_j\}_i := \begin{cases} 1, & \text{if } i = j, \\ 0, & \text{if } i \neq j. \end{cases}$$

Since the reconstruction $X^{\text{rec}}_{:,t}$ is continuous function of $X_{:,t}$, we can write

$$\left\| \frac{\partial X^{\text{rec}}_{:,t}}{\partial X_{j,t}} \right\|_2^2 = \left\| \lim_{h\to 0} \frac{X^{\text{rec}}_{:,t,h} - X^{\text{rec}}_{:,t}}{h} \right\|_2^2 = \lim_{h\to 0} \frac{1}{h^2} \|X^{\text{rec}}_{:,t,h} - X^{\text{rec}}_{:,t}\|_2^2$$

The inner norm can be estimated using (59) to get

$$\lim_{h\to 0} \frac{1}{h^2} \|X^{\text{rec}}_{:,t,h} - X^{\text{rec}}_{:,t}\|_2^2 \leq \lim_{h\to 0} \frac{1}{h^2} \|X_{:,t,h} - X_{:,t}\|_2^2 = 1$$

□

**Corollary 10.** *The previous Lemma motivates for using the regularization of $S$-problem (25). In the case of $K = 2$, such a regularization minimizes the norm of derivative (61). In the case of general $K$, this regularization modifies the resulting polytope generated by $S^*$ in a such way that this polytope is distinguishing between the features of reconstructed data, see (60).*

**Corollary 11.** *Please, notice that the dependence of reconstruction of $X^{\text{rec}}$ on data $X$ is linear and the respective derivative is piecewise constant, see Corollary after Lemma 14. In practice, we can estimate the norm in (62) using Euler method. Due to discontinuities in derivatives, such a method is exact for sufficiently small step $h$.*

### Figures

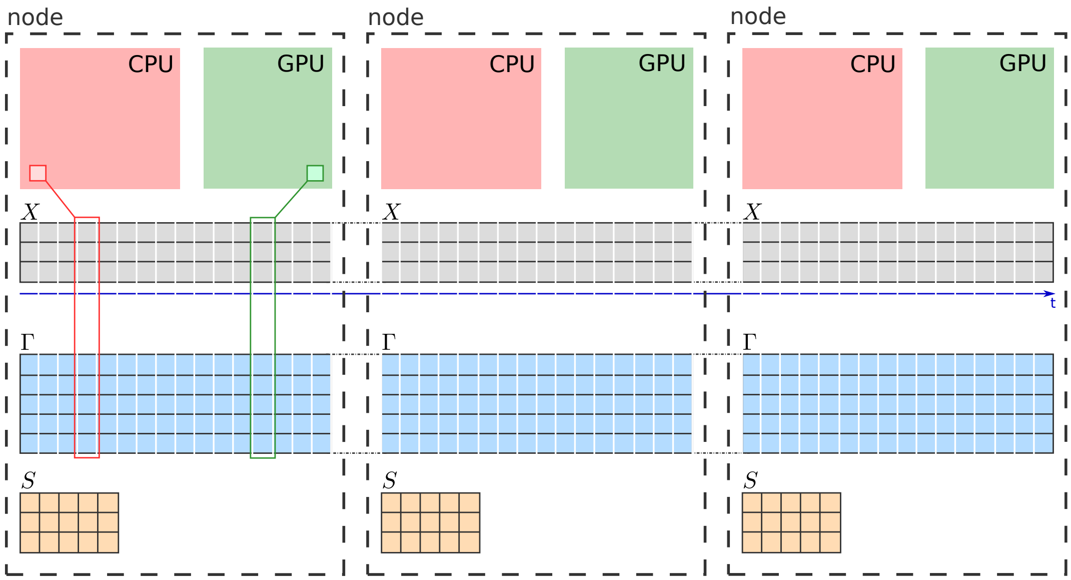


Figure S1: **Distributed solution of $\Gamma$-problem:** If objective function in (SPA), ($SPA_2$) is additively separable in $t$ then the solution of optimization problem with fixed $S$ can be composed as a solution of individual problems (see Lemma 4 and Lemma 11). In such a case, we can distribute $T$ independent problems into several computation nodes such that the each node solves its own subset of problems. This local computation can be be performed by local CPU cores and/or using GPU cores, where (again) each core solves its individual subset of local optimization problems. Additionally, if we distribute the data of the problem in the same way, then each computational resource will have an access to its own local part of memory, without any additional communication.

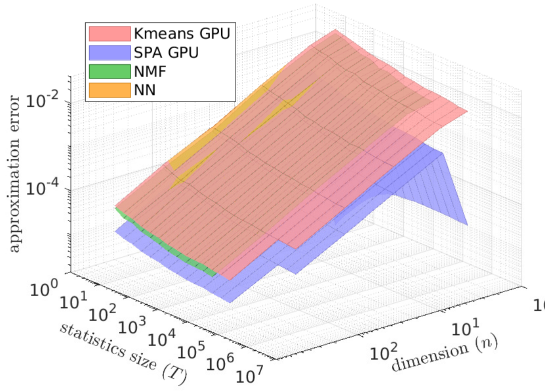


(a) approximation quality scaling

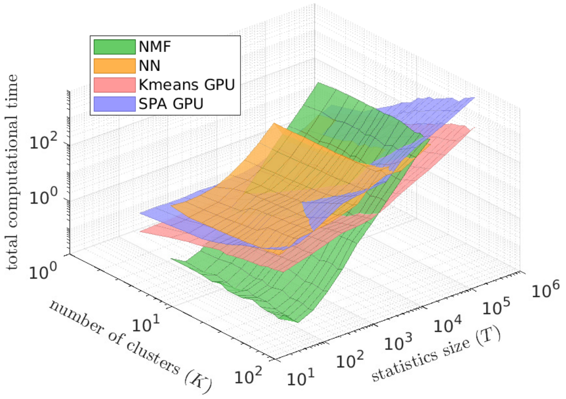


(b) computational cost scaling

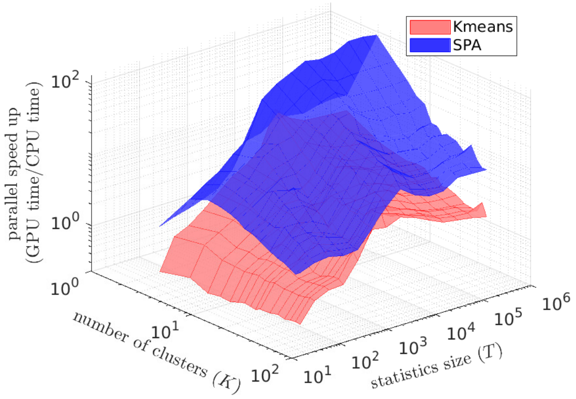


(c) parallelisability scaling

Figure S2: **Comparing computational cost (a), discretization quality (b) and parallelizability (c):** for ($SPA_2$) (blue surfaces), K-means clustering (dark-green), Nonnegative Matrix Factorisation (in its probabilistic variant called Left-Stochastic Decomposition (LSD), magenta surfaces) and the Self-Organising Maps (SOM, a special form of unsupervised neuronal networks used for discretization, orange surfaces). For every combination of data dimension $n$ and the data statistics length $T$, methods are applied to 50 same randomly-generated data sets and the results in each of the curves represent averages over these 50 problems. Parallel speed-up in (c) is measured as the ratio of the average times time(GPU)/time(CPU) needed to reach the same relative tolerance threshold of $10^{-5}$ on a single Graphics Processing Unit (GPU, ASUS TURBO-GTX1080TI-11G, with 3584 CUDA cores) for time(GPU) versus a single CPU core (Intel Core i9-7900X CPU) for time(CPU). MATLAB script Fig1_reproduce.m reproducing these results is available for open access in the repository SPA at https://github.com/SusanneGerber.

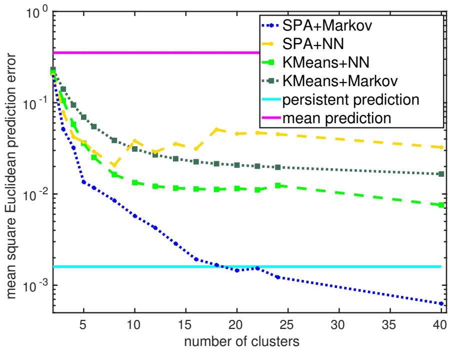


(a) Lorenz-96 1D turbulence model (weakly-chaotic regime)

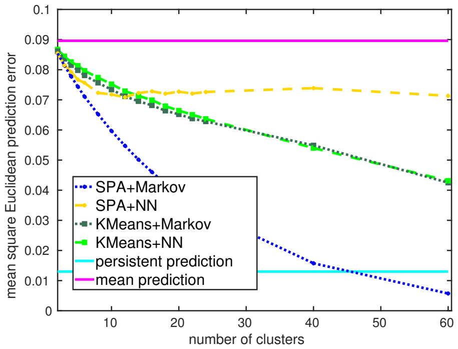


(b) Lorenz-96 1D turbulence model (strongly-chaotic regime)

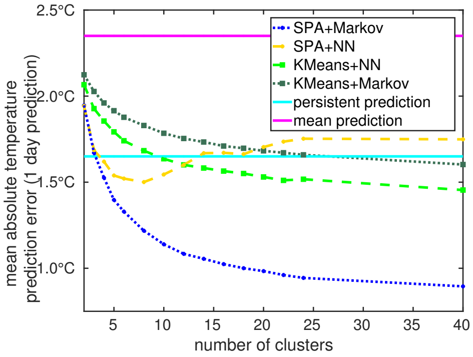


(c) surface temperature dynamics over Europe (1979-2010, 20x30 grid ECMWF resimulation data)

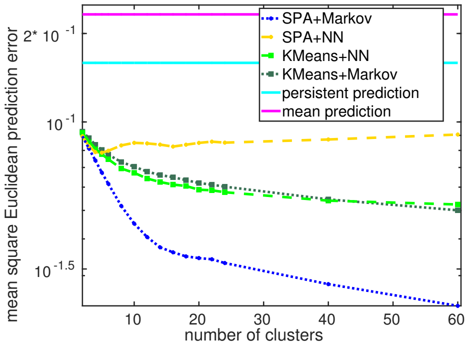


(d) molecular dynamics simulation of 10-Alanine in water

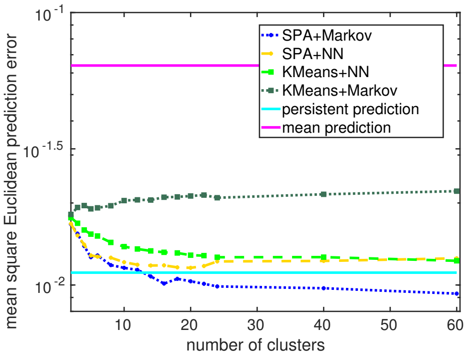


(e) EG dynamics in a brain-computer interface (BCI2000 data)

Figure S3: Comparison of one-time-step predictions for a combination of SPA with Markov models (based on applications of the Theorem 2, blue lines) to the one-time-step predictions obtained by the standard prediction methods. The combination of SPA with Markov models is the only prediction scheme that outperforms the persistent prediction (i.e., when the next state is predicted to be the same as the current one) for all of the considered systems.

### APPENDIX

**Definition 1.** *We say that point $x^*$ is a minimizer of function $f$ on given feasible set $\Omega$, written as*

$$x^* = \arg\min_{x\in\Omega} f(x),$$

*if (and only if) all points from the feasible set have larger or equal function value than $f(x^*)$, i.e.,*

$$\forall x \in \Omega : f(x^*) \leq f(x).$$

**Lemma 18.** *Let $X \in \mathbb{R}^{n,T}, a, x \in \mathbb{R}^n, b \in \mathbb{R}^n, A = A^T \in \mathbb{R}^{n,n}$. Then*

$$\frac{\partial a^T X b}{\partial X} = ab^T, \qquad \frac{\partial b^T X^T X b}{\partial X} = 2Xbb^T, \qquad \frac{\partial x^T a}{\partial x} = a, \qquad \frac{\partial x^T A x}{\partial x} = 2Ax.$$

**Lemma 19.** *Let $n, K, T \in \mathbb{N}$ and $A \in \mathbb{R}^{n,T}, B \in \mathbb{R}^{K,T}$. Then*

$$\sum_{t=1}^{T} A_{:,t}(B_{:,t})^T = AB^T \in \mathbb{R}^{n,K}.$$

*Proof.* From the definition of matrix-vector multiplication, the components of the result on left-hand side of the equation can be written in form (for every $i \in \{1,\ldots,n\}, j \in \{1,\ldots,K\}$)

$$\left[\sum_{t=1}^{T} A_{:,t}(B_{:,t})^T\right]_{i,j} = \sum_{t=1}^{T} A_{i,t}(B_{j,t})^T = \langle A_{i,:}, B_{j,:}\rangle = A_{i,:}(B_{j,:})^T,$$

which is a value of the corresponding matrix component on right-hand side of the equation. □

**Lemma 20.** *(of four fundamental subspaces): for any $B \in \mathbb{R}^{n,m}$ it holds*[3]

$$\operatorname{Ker} B \perp \operatorname{Im} B^T, \qquad \operatorname{Im} B \perp \operatorname{Ker} B^T.$$

$$\operatorname{Ker} B \cup \operatorname{Im} B^T = \mathbb{R}^m, \qquad \operatorname{Im} B \cup \operatorname{Ker} B^T = \mathbb{R}^n.$$

[3] Let $\mathcal{V}, \mathcal{W}$ be two subspaces of vector space $\mathcal{F}$ with scalar product $\langle .,. \rangle : \mathcal{F} \times \mathcal{F} \to \mathbb{R}$. Then we say that $\mathcal{V} \perp \mathcal{W}$ if $\forall v \in \mathcal{V} \forall w \in \mathcal{W} : \langle v, w \rangle = 0$. Additionally we define $\mathcal{V} \cup \mathcal{W} := \{f \in \mathcal{F} : f \in \mathcal{V} \ \vee \ f \in \mathcal{W}\}$.

*Proof.* See Laub (*8*). ☐

**Lemma 21.** *Let* $n, K, T \in \mathbb{N}$ *and* $A \in \mathbb{R}^{n,T}, B \in \mathbb{R}^{K,T}$. *Then*

$$\operatorname{Ker} AA^T = \operatorname{Ker} A^T \subset \mathbb{R}^n, \tag{66}$$

$$\operatorname{Ker} B \subset \operatorname{Ker} AB \subset \mathbb{R}^K. \tag{67}$$

*Proof.* To prove (66), it is necessary to show that

$$\forall x \in \mathbb{R}^n : \quad AA^T x = 0 \quad \Leftrightarrow A^T A = 0.$$

($\Leftarrow$) Let us consider $x \in \mathbb{R}^m$ such that $A^T x = 0$. Then $AA^T x = A \underbrace{A^T x}_{=0} = 0$ (this also proves (67))

($\Rightarrow$) Let us consider $x \in \mathbb{R}^m$ such that $AA^T x = 0$. Using smart zero, we can write

$$0 = x^T 0 = x^T AA^T x = \|A^T x\|^2.$$

The norm of the vector is equal to zero if and only if the vector is equal to zero, therefore $A^T x = 0$.

☐

**Lemma 22.** *Let* $f : \mathbb{R}^n \to \mathbb{R}$ *be a continuously differentiable convex function and let* $\Omega \subset \mathbb{R}^n$ *be closed convex set. Then* $x^* \in \Omega$ *is a solution of optimization problem*

$$x^* := \arg\min_{x \in \Omega} f(x)$$

*if and only if*

$$\forall x \in \Omega : \langle \nabla f(x), x - x^* \rangle \geq 0.$$

*Proof.* See (*3*), (*10*). ☐
